# Structurally distinct manganese-sensing riboswitch aptamers regulate different expression platform architectures

**DOI:** 10.1101/2024.12.14.628514

**Authors:** Christine N. Stephen, Danea E. Palmer, Clarisa Bautista, Tatiana V. Mishanina

## Abstract

Manganese (Mn)-sensing riboswitches protect bacteria from Mn toxicity by upregulating expression of Mn exporters. The Mn aptamers share key features but diverge in other important elements, including within the metal-binding core. Although X-ray crystal structures of isolated aptamers exist, these structural snapshots lack crucial details about how the aptamer communicates the presence or absence of ligand to the expression platform. In this work, we investigated the Mn-sensing translational riboswitches in *E. coli* (*mntP* and *alx*), which differ in aptamer secondary structure, nucleotide sequence, and pH-dependence of Mn response. We performed co-transcriptional RNA chemical probing, allowing us to visualize RNA folding intermediates that form and resolve *en route* to the final folded riboswitch. For the first time, we report that sampling of metal ions by the RNA begins before the aptamer synthesis and folding are complete. At a single-nucleotide resolution, we pinpoint the transcription window where “riboswitching” occurs in response to Mn binding and uncover key differences in how the *alx* and *mntP* riboswitches fold. Finally, we describe riboswitch-specific effects of pH, providing insights into how two members of the same riboswitch family differentially sense two distinct environmental cues: concentration of Mn and pH.

**GRAPHICAL ABSTRACT:** 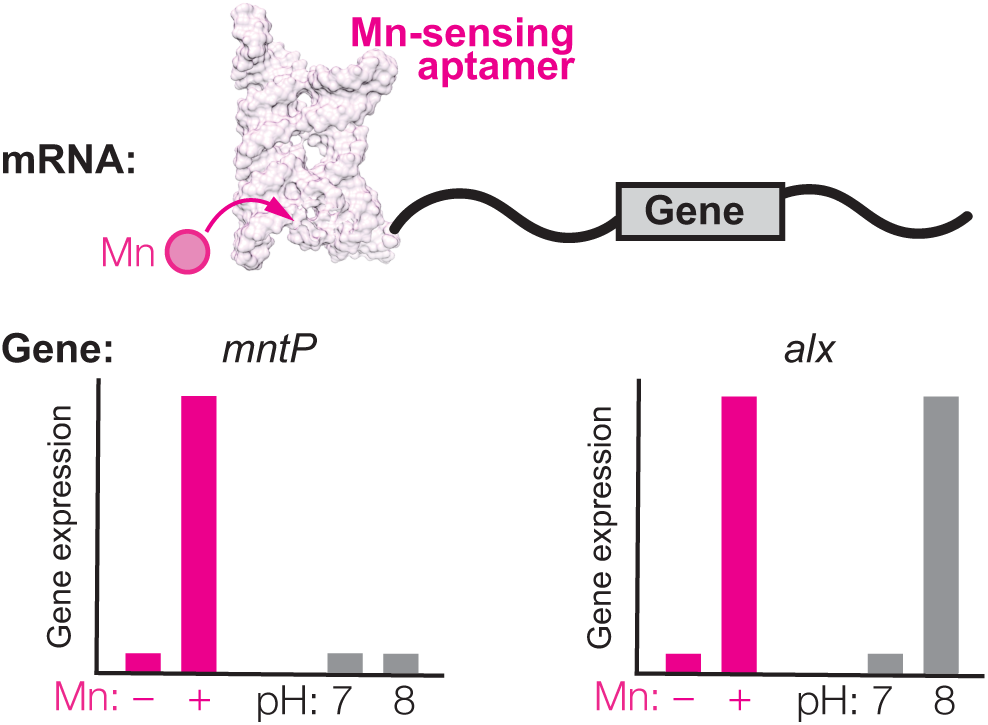

## INTRODUCTION

Transition metal ions play key cellular roles (e.g., serving as enzyme cofactors) but become harmful in excess due to misincorporation into biomolecules and metal-catalyzed formation of reactive oxygen species(1, 2). To match the need for metal ion transporters, cells use the metal’s intracellular concentration to regulate expression of the metal’s transporters at the transcription and/or translation level(3). Examples of this regulation include metal-binding transcription regulators(4, 5) or direct interaction of the metal with a riboswitch encoded in the mRNA(6–8). Riboswitches are 5’-untranslated regions of mRNA that promote or inhibit expression of the downstream gene by changing their folding upon binding a ligand(9). A riboswitch is comprised of a ligand-binding aptamer overlapping with an expression platform, whose folding determines the transcriptional and/or translational fate of the mRNA(9). Many classes of metal-sensing riboswitches have been identified, such as magnesium(7), nickel(8), cobalt(8), and manganese (Mn)(6, 10).

The *yybP-ykoY* Mn-sensing riboswitch family is the largest metal-sensing family, with >1, 000 unique representatives in different bacterial phyla(11). Crystal structures, single-molecule FRET (smFRET), and molecular dynamics (MD) simulations of isolated Mn-sensing aptamers provided valuable insights into the overall RNA fold, metal binding site, and RNA dynamics in the presence or absence of Mn(12, 13). However, these studies lack the expression platform of the riboswitch – the “business end” that determines the gene expression outcome. Consequently, it remains unknown how Mn binding to the aptamer actuates the differential folding of the downstream RNA to affect its transcription or translation. Further, studies to date were performed on pre-synthesized RNA, missing the aspect of active transcription that is critical to riboswitch folding(14–17).

We addressed these gaps by investigating the two Mn-sensing translational riboswitches in *E. coli* which regulate translation of the *alx* and *mntP* genes, with the *alx* riboswitch upregulating gene expression in response to alkaline pH and increased Mn concentration ([Mn])(18, 19) (Fig. 1A). Binding of Mn to either the *mntP* or *alx* aptamers promotes a translationally active conformation in the expression platform, increasing expression of the Alx and MntP Mn exporters(10, 19, 20) (Fig. 1A-B). Multiple structures of Mn-sensing riboswitch aptamers established their core structural features, consisting of two coaxially stacked “legs” connected by a four-way junction (4WJ)(6, 12, 21) (Fig. 1C). The right leg (L1, P1.1, P1.2, and P2; Fig. 1C) extrudes a conserved adenosine in L1 to form a cross-helix A-minor interaction with P3.1 in the left leg (L3, P3.1, P3.2, and P4; Fig. 1C). The L3 loop in the left leg stacks below the conserved adenosine in the A-minor interaction, forming a phosphate-rich metal-binding hot spot (Fig. 1C). X-ray crystal structures show two metal-binding sites in the aptamer, with one occupied by Mg^2+^ (M_A_) and the other by Mn^2+^ (M_B_)(6, 12, 21) (Fig. 1C). The P1.1 “switch helix” is a key element of the *yybP-ykoY* aptamer as its 3’-half alternatively pairs within the aptamer or the expression platform depending on whether Mn is bound, with consequences to folding of the rest of the expression platform.(12) (see Fig. 1B for *alx* and *mntP* alternative P1.1 pairing).

**Figure 1.**
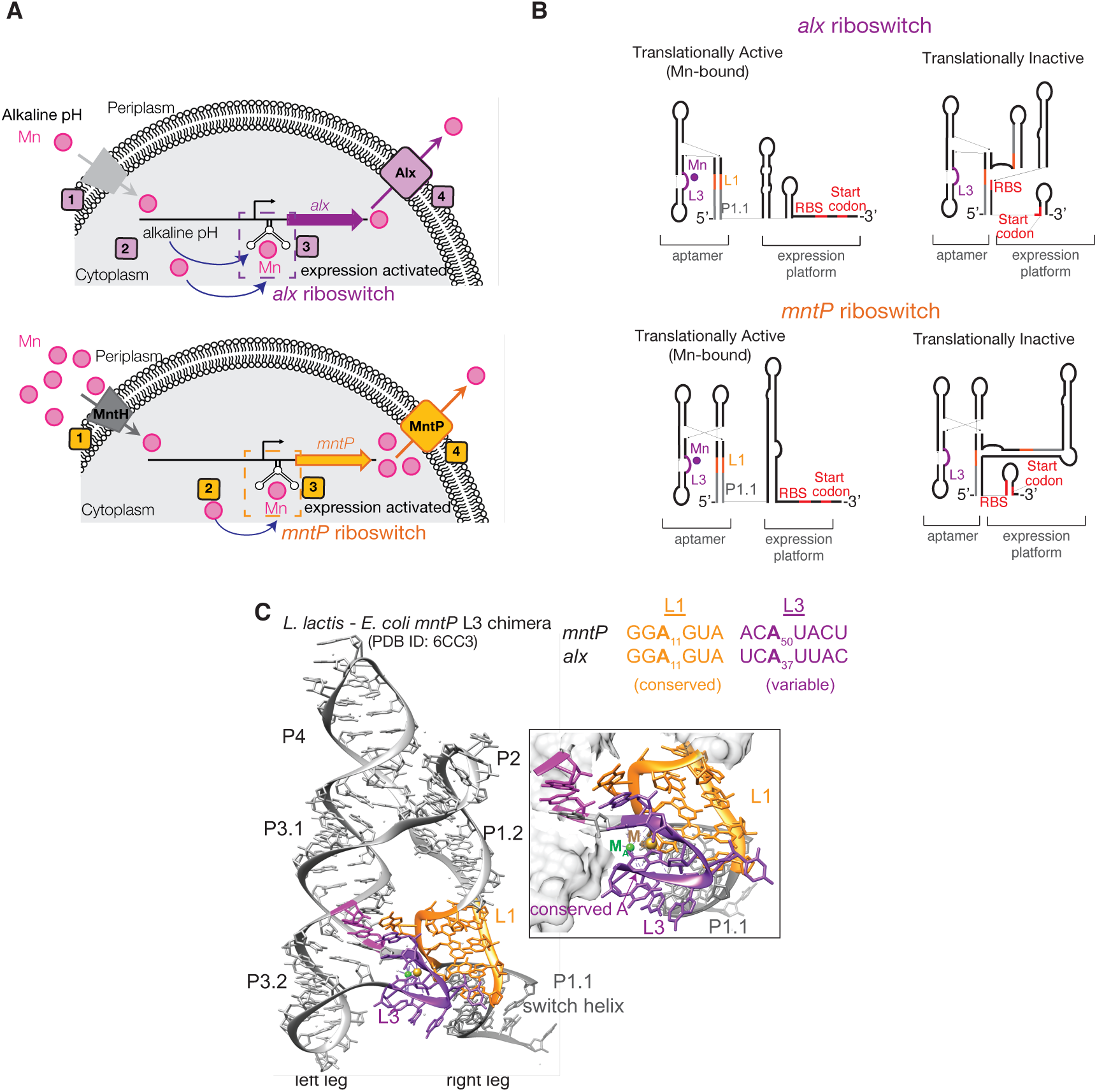
Introduction to the *E. coli alx* and *mntP* Mn-sensing translational riboswitches. **A.** Cartoon model of *alx* and *mntP* riboregulation of their respective genes in *E. coli*. (Top panel) At alkaline pH, the cytosolic [Mn] increases in *E. coli*, resulting in the *alx* riboswitch turning “on” expression of Alx, an Mn exporter. (Bottom panel) An increase in environmental [Mn] results in the *mntP* riboswitch turning “on” expression of MntP, the major Mn exporter in *E. coli*. **B.** Cartoon models of the *alx* and *mntP* riboswitches in their translationally active vs. inactive conformations. L3 (purple), L1 (orange), P1.1 (gray), and RBS/start codon (red) are highlighted. **C.** *L. lactis* Mn-sensing aptamer crystal structure with the *E. coli mntP* L3 sequence (PDB 1D: 6CC3). “Left leg” and “right leg” refer to coaxially stacked helices in the aptamer. L3 (purple), L1 (orange), P1.1 (gray), and G-C receptor base pair in cross-helix A-minor (magenta) are highlighted. Mg^2+^ in the M_A_ site and Cd^2+^ in the M_B_ site (captured instead of Mn^2+^) are green and yellow spheres, respectively. Sequence alignment of *alx* and *mntP* L1 and L3 are orange and yellow, respectively.

Despite sharing conserved features for Mn-sensing, the *alx* and *mntP* riboswitches differ in their global architectures (Fig. 2A). The *alx*-like riboswitches are exclusively present in the *Enterobacteriales* order, members of which are often found in environments that can alkalinize, such as water, soil, and in association with other organisms(22, 23). Upon alkaline pH challenge, the *E. coli* cytosol alkalinizes and then recovers to pH 7.8(24); in turn, cytosol alkalinization increases cytoplasmic [Mn](19), potentially explaining *alx* translational activation at alkaline pH. Initial studies of the *alx* and *mntP* riboswitches put forward alternative secondary structure models based on chemical probing with pre-synthesized RNA(10, 18), prompting us to ask (i) how these alternative structures fold and respond to Mn during transcription and (ii) why alkaline pH uniquely affects *alx* structure. To answer these fundamental questions, we applied modern RNA chemical probing methods to *alx* and *mntP*, where ribonucleotides are modified by either a selective 2’-hydroxyl (SHAPE) reagent or dimethyl sulfate (DMS) and detected by high-throughput mutational profiling (MaP)(25).

**Figure 2.**
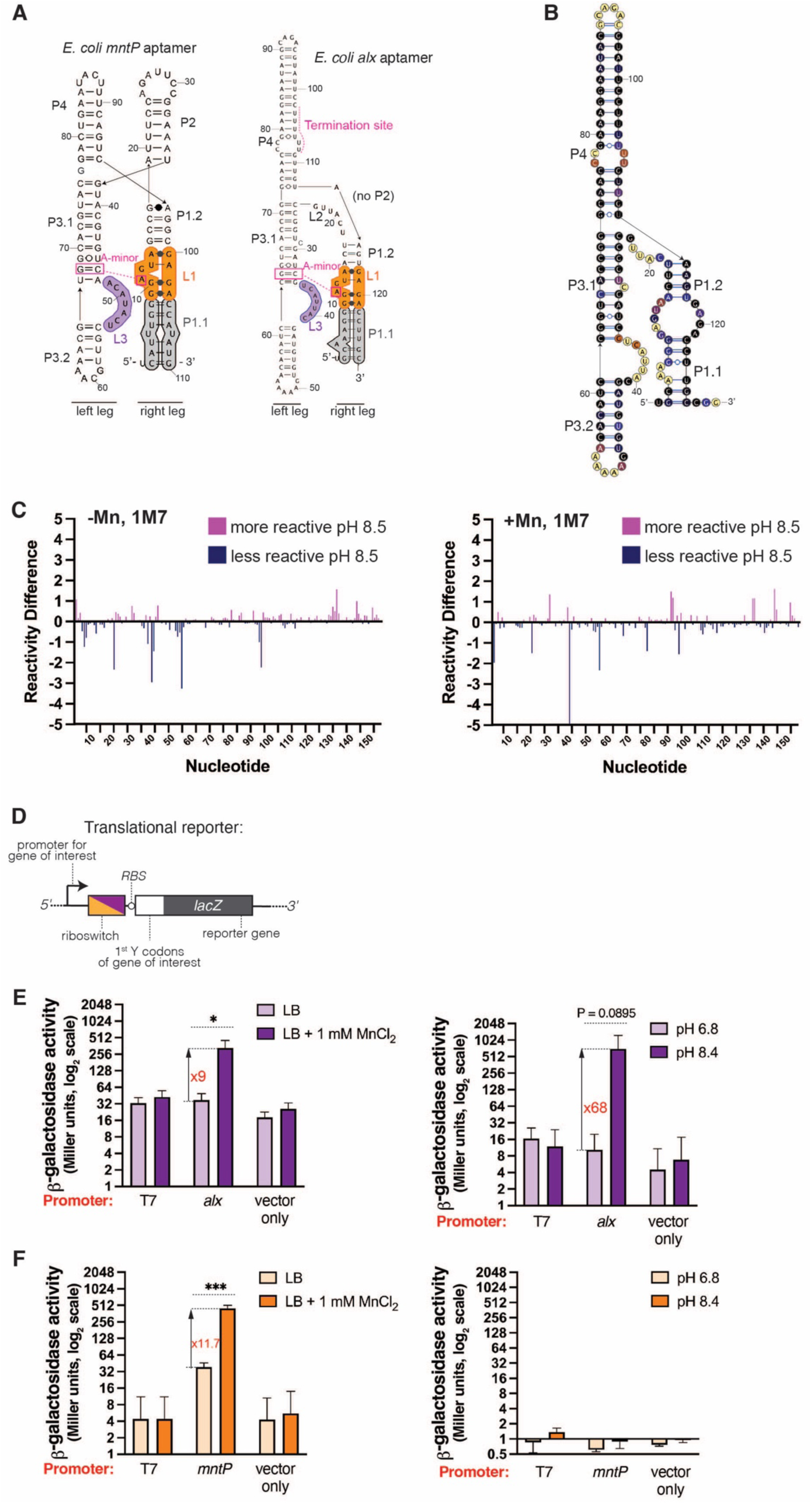
Aptamer architectures and riboswitch-controlled translational response *in vivo*. **A.** *mntP* and *alx* aptamer structure models showing the *mntP* 4WJ and *alx* 3WJ elements. The key Mn-sensing loops (L1 is orange and L3 is purple) and the P1.1 switch helix (gray) are highlighted. **B.** Secondary structure model of equilibrium-refolded (pH 7.2 –Mn) *alx* aptamer modeled based on reactivity with 1M7. **C.** Individual nucleotide reactivity differences when modified with 1M7 at neutral (7.2) vs alkaline pH (8.5), measured in the presence or absence of Mn. **D.** Translational reporter construct design for *alx* and *mntP* translational reporters with the native or T7 promoter. **E.** Translational response of *alx* to supplemented Mn (left) or alkaline pH (right) as a function of promoter. Statistical analysis done using a parametric unpaired t-test to assess significance. **F.** Translational response of *mntP* to supplemented Mn (left) or alkaline pH (right) as a function of promoter. Statistical analysis done using a parametric unpaired t-test to assess significance.

In this work, we performed DMS-MaP in a co-transcriptional format using established methods(26, 27), getting to the heart of how Mn binding to the aptamer is “communicated” to the expression platform. Unexpectedly, we uncovered evidence that both *alx* and *mntP* aptamer folding intermediates begin sampling for divalents before the full aptamer domain has emerged from RNAP, challenging the notion that a fully folded aptamer is required for ligand sensing. Further, a critical RNA loop that participates in Mn sensing diverges in sequence between *alx* and *mntP* and may hold the key to how pH differentially affects riboregulation by these riboswitches. Using gene reporter assays, we show that alkaline pH alone cannot rearrange the *alx* riboswitch into a translationally active conformation; instead, the *alx* riboswitch senses the increased [Mn] brought about by alkaline pH. Lastly, our co-transcriptional chemical probing data uncovered distinct structural features in the *alx* and *mntP* expression platforms, which are potentially coupled to the cellular need for the Alx and MntP exporter proteins under increased cytosolic [Mn].

## MATERIAL AND METHODS

The source and catalog number (if applicable) for all key reagents are provided as a Supplementary Table 1.

All oligonucleotides were purchased from either Integrated DNA Technologies or Eton Bioscience. A summary of all oligonucleotides used in this work including sequences, modifications, and purifications is presented in Supplementary Table 2.

Procedures for protein preparations, construction of plasmids and bacterial strains, and preparation of DNA templates are provided in the Supplemental Methods. Details for all DNA templates, strains, and plasmids are tabulated in Supplementary Tables 3-6.

### Single-round *in vitro* transcription for TECprobe-VL experiments

To prepare the holoRNAP, *E. coli* RNAP (6 µM) and 0^70^ (12 µM) were combined in 1x TECprobe transcription buffer (100 mM Tris-Cl, pH 7.2 or 8.5, 50 mM KCl, and 1 mM DTT) and incubated for 30 min at 37 °C. The single-round *in vitro* transcription reactions for TECprobe-VL experiments were assembled by first combining 1x transcription buffer, 40 nM DNA template (*alx* riboswitch, Template 3, Supplementary Table 3) or 50 nM DNA template (*mntP* riboswitch, Template 6, Supplementary Table 3) with randomly distributed internal biotin-11 dNMPs (AAT BioQuest/Biotium), 0.5 µM holoRNAP, and 500 µM ATP, UTP, CTP, and GTP. The transcription buffer omitted magnesium, which was a part of the start solution added later to initiate transcription elongation. Tris-Cl was added at a final concentration of 100 mM to minimize pH changes during the chemical probing reaction, as described previously(27). The reaction was incubated for 10 min at 37 °C to form open promoter complexes. Next, SAvPhire Monomeric Streptavidin (Sigma-Aldrich) was added to the reaction at a final concentration of 5 µM, followed by incubation for 10 min at 37 °C to allow streptavidin binding to biotinylation sites on the DNA template. Transcription was initiated by adding 10X start solution (100 mM MgCl_2_, 100 µg/ml rifampicin) to the reaction to a final concentration of 10 mM MgCl_2_ and 10 µg/ml rifampicin to inhibit reinitiation. After addition of start solution, the final reaction volume was 60 µl. To test the effect of the Mn ligand on riboswitch structural rearrangements, MnCl_2_ was added to a final concentration of 1 mM in the transcription reactions and was introduced alongside the start solution. The transcription reaction was incubated for 30 s at 37 °C before proceeding to the chemical probing, which is described in the next section.

Single-round *in vitro* transcription for the single-length TECprobe-SL experiments followed the same protocol as outlined above for variable length TECprobe-VL experiments. The DNA templates used for TECprobe-SL experiments contained a terminal biotin modification that was introduced by a 5’ biotinylated reverse primer in the PCR (*alx* riboswitch, Template 7, Supplementary table 3).

### RNA chemical probing during TECprobe-VL experiments

Benzoyl cyanide (BzCN, Sigma-Aldrich) probing was performed by splitting the 60-µl transcription reaction into 25 µl aliquots and mixing with either 2.78 µl of 400 mM BzCN dissolved in anhydrous DMSO (Invitrogen) to a final concentration of 17.7 mM [(+) sample] or with anhydrous DMSO [(-) sample]. Since the BzCN probing reaction is complete within ∼1 s, no additional incubation was required before quenching the reactions with 250 µl of TRIzol reagent (Invitrogen).

DMS probing was performed by splitting the 60-µl transcription reaction into 25 µl aliquots and mixing with either 2.78 µl of 6.5% v/v DMS (Sigma-Aldrich) in 200 proof ethanol [(+) channel] or with 200 proof ethanol [(-) channel]. The samples were incubated for 5 min at 37 °C. The DMS probing reaction was quenched by adding 6.67 µl of 14.5 µM 2-mercaptoethanol and incubating the sample at 37 °C for 1 min. *In vitro* transcription reactions were then quenched by addition of 250 µl of TRIzol reagent.

### RNA purification post-TECprobe-VL probing

The RNA was purified from the quenched *in vitro* transcription reactions using the following protocol. First, 50 µl of chloroform was added to the samples and the samples were mixed by vortexing followed by centrifugation at 18, 000*g* and 4 °C for 5 min. Next, the aqueous layer was transferred to a new tube and nucleic acids were precipitated by addition of 105 µl cold isopropanol and 1 µl GlycoBlue Coprecipitant (Invitrogen). The samples were incubated at room temperature for 15 min and centrifuged at 18, 000*g* and 4 °C for 15 min. The supernatant was discarded and the pellet containing the precipitated nucleic acids was washed by addition of 200 µl 70% ethanol. Following the 70% ethanol wash, the samples were centrifuged at 18, 000*g* and 4 °C for 2 min and the supernatant was discarded. The pelleted nucleic acids were resuspended in 25 µl of 1x DNase buffer (10 mM Tris-Cl, pH 7.5, 2.5 mM MgCl_2_) and mixed with 0.75 µl of Turbo DNase (Invitrogen) to remove residual DNA template. The samples were incubated at 37 °C for 15 min and DNase digestion was quenched by addition of 250 µl TRIzol reagent (Invitrogen). A second TRIzol extraction was performed as described above, except the pellet containing the DMS-modified RNA transcripts was resuspended in 5 µl of 10% (v/v) DMSO.

### RNA 3’ adapter ligation onto the TECprobe-VL purified transcripts

The 9N_VRA3 adapter oligonucleotide (0139, Supplementary Table 2) was pre-adenylated with the 5’ DNA Adenylation Kit (New England Biolabs) using the following protocol. A 100 µl master mix was prepared that consisted of the 1X DNA Adenylation Buffer (New England Biolabs), 100 µM ATP, 5 µM 9N_VRA3 oligo, and 5 µM Mth RNA Ligase (New England Biolabs). The 100 µl master mix was split into 4 aliquots (25 µl each) in PCR tubes and incubated in a thermal cycler set to 65 °C for 1 h. To clean up the adenylation reactions, reactions were pooled to 50 µl followed by addition of 500 µl TRIzol (Invitrogen) and 100 µl of chloroform to each 50 µl pool. The samples were vortexed to mix and centrifuged at 18, 000*g* and 4 °C for 5 min. The aqueous phase was transferred to a new tube and the adapter was ethanol precipitated. The pellet was resuspended in Tris-Cl, pH 8.0, and its concentration was quantified with the Qubit ssDNA Assay Kit (Invitrogen) and a Qubit 3 Fluorometer. The pre-adenylated adapter was diluted to 1 µM, divided into small aliquots, and stored at -20 °C.

The RNA 3’ adapter ligation reactions were performed using the following protocol. Purified RNA redissolved in 5 µl of 10% (v/v) DMSO was combined with 1x T4 RNA Ligase Buffer (New England Biolabs), 0.5 U/µl SuperaseIN (Invitrogen), 0.05 µM pre-adenylated 9N_VRA3 adapter, 15 % PEG 8000, and 5 U/µl T4 RNA Ligase 2, truncated KQ (New England Biolabs). The final volume for the ligation reactions was 20 µl and the reactions were incubated at 25 °C for 2 h.

### SPRI bead purification of adapter-ligated RNA

A solid-phase reversible immobilization (SPRI) bead purification protocol was used to deplete excess 9N_VRA3 adapter oligonucleotide. The SPRI beads were prepared with SpeedBead Magnetic Carboxylate Beads (Cytiva) using the protocol from Jolivet and Foley(28). The ligation reactions were combined with 17.5 µl of water, 40 µl anhydrous isopropanol, and 22.5 µl of the SPRI beads. The samples were vortexed and incubated at room temperature for 5 min. The samples were then placed on a magnetic separation rack and the beads collected on the tube wall before disposing of the supernatant. Keeping the tubes on the magnetic stand, the beads were washed twice with 200 µl of 70% ethanol. After discarding the second ethanol wash, the tubes were placed in a dry bath set to 37 °C to evaporate the residual ethanol. The beads were resuspended in 25 µl of elution buffer (20 mM Tris-Cl pH 8.0, 50 mM KCl, 1 mM DTT, 0.1 mM EDTA, 0.1% Triton X-100, 10 mM MgCl_2_) and the samples were incubated for 3 minutes at room temperature to elute the RNA. The samples were placed on a magnetic rack and beads collected on the tube wall before aliquoting the supernatant containing the RNA into a fresh tube.

### TECprobe-VL cDNA synthesis and cleanup

The reverse transcription reactions were performed via capture of RNA transcripts on streptavidin beads using a 5’ biotinylated reverse transcription primer (0159, Supplementary table 2), which anneals to the 9N_VRA3 sequence that was appended to the intermediate riboswitch transcripts. Dynabeads MyOne Streptavidin C1 beads (Invitrogen) were equilibrated as described previously(27). 5 µl of 500 nM RT primer was combined with the purified RNA samples and annealed in a thermal cycler using the following program: 70 °C for 5 min, ramp to 50 °C at 0.1 °C /s, 50 °C for 5 min, ramp to 40 °C at 0.1 °C /s, 40 °C for 5 min, cool to 25 °C. The equilibrated streptavidin beads were placed on a magnetic stand to allow the beads to collect on the tube wall and the supernatant was discarded. The streptavidin beads were then resuspended with the samples containing RNA annealed to the 5’ biotinylated RT primer and incubated at room temperature for 15 min on an end-over-end rotator. The samples were placed on a magnetic stand and the beads were allowed to collect on the tube wall before discarding the supernatant. The beads were resuspended in 19.5 µl of reverse transcription master mix and incubated in a thermal cycler set to 42 °C for 2 min before addition of the reverse transcriptase. Upon addition of 0.5 µl of SuperScript II Reverse Transcriptase (Invitrogen) to each sample, the 20 µl RT reactions contained 1x first strand buffer (50 mM Tris-Cl, pH 8.0, 75 mM KCl), 500 µM dNTPs, 10 mM DTT, 2 % formamide, 10 ng/µl Extreme Thermostable Single-Stranded DNA Binding Protein (ET SSB) (New England Biolabs), 0.1 % Triton-X-100, and 3 mM MnCl_2_. The reverse transcription reactions were incubated at 42 °C for 180 min followed by a 70 °C incubation for 50 min to inactivate the SuperScript II and then cooled to 12 °C. Next, the RNAs were degraded by adding 1.25 U RNase H (New England Biolabs) and 12.5 U RNase I_f_ (New England Biolabs) to each sample and the samples were incubated at 37 °C for 20 min followed by a 70 °C incubation for 20 min to inactivate the RNases. The samples were placed on a magnetic stand and the beads were allowed to collect on the tube wall. The supernatant was discarded and the beads were resuspended in 75 µl of Tris Wash Buffer (10 mM Tris-Cl, pH 8.0, 0.05% Triton X-100) to wash the beads. The samples were placed on a magnetic stand and the wash was discarded. The bead-bound cDNA was resuspended in 25 µl of the Tris Wash Buffer and stored at -20 °C.

### Chemical probing of equilibrium-refolded intermediate transcripts

Single-round *in vitro* transcription using randomly biotinylated DNA templates was executed as described for TECprobe-VL experiments above. To quench transcription, 600 µl of TRIzol was added to the reactions. Next, 120 µl of chloroform was added to the samples and the samples were vortexed to mix followed by centrifugation at 18, 000*g* and 4 °C for 5 min. The aqueous layer was transferred into a new tube and nucleic acids were isopropanol precipitated as described for TECprobe-VL experiments except a larger volume of isopropanol was used to account for the larger sample volume. The pellets containing the intermediate RNA transcripts were resuspended in 30 µl of 1x TECprobe transcription buffer (100 mM Tris-Cl, pH 7.2 or 8.5, 50 mM KCl, 1 mM DTT), placed on a heat block set to 95 °C for 2 min, and snap-cooled on ice for 1 min. To each sample, 30 µl of 2x equilibration buffer (200 mM Tris-Cl, pH 7.2 or 8.5, 100 mM KCl, 2 mM DTT, 20 mM MgCl_2_) was added, which contained either no Mn or 2 mM MnCl_2_ (1 mM final in 60 µl reaction), and the samples were incubated at 37 °C for 20 min. The RNAs were then probed with DMS as described for TECprobe-VL experiments, and the DMS-probed RNA was purified as described for TECprobe-VL experiments.

### Preparation of TECprobe-VL dsDNA libraries for sequencing

cDNA libraries were PCR-amplified for high throughput sequencing using the following protocol. Separate PCR reactions were performed for the (+) and (-) samples, which contained 1x Q5 Reaction Buffer, 1x Q5 GC Enhancer, 200 µM dNTPs, 250 nM RPI Indexing Primer (0163-0178, 0235-0248, Supplementary table 2), 250 nM dRP1_NoMod.R reverse primer (0162, Supplementary Table 2), 10 nM SC1Brdg_MINUS or SC1Brdg_PLUS channel barcode oligo (0160, 0161, Supplementary Table 2), 12 µl of bead-bound cDNA library, and 0.02 U/µl Q5 High-Fidelity DNA Polymerase (New England Biolabs). Amplification was performed with the following thermal cycler program: 98 °C for 30 s, [98 °C for 10s, 62 °C for 20 s, 72 °C for 20s] x 22 cycles, 72 °C for 2 min, hold at 12 °C.

### High-throughput DNA sequencing of TECprobe-VL libraries

Sequencing of the TECprobe-VL and equilibrium refolded intermediate transcript libraries was performed by Novogene Co. on a Novaseq X Plus system with 2 x 150 Paired End (PE) reads and a 20% PhiX spike in. The TECprobe-VL and equilibrium refolded intermediate transcript libraries were sequenced at a depth of ∼30-60 million PE reads.

Sequencing of TECprobe-SL libraries was performed by Novogene Co. on a Novaseq X Plus system with 2 x 150 PE reads. No PhiX spike in was added since these libraries were sequenced in a partial lane. The TECprobe-SL libraries were sequenced at a depth of ∼10 million PE reads. Sequencing data for TECprobe-VL and TECprobe-SL experiments is publicly available in the Sequencing Read Archive (SRA) under project number PRJNA1197522 and BioSamples are tabulated in Supplementary Table 7.

### TECprobe-VL data processing and visualization

Sequence read pre-processing, target generation using cotrans_preprocessor, and shapemapper2 run script generation were performed exactly as described in the published protocol using the publicly available TECprobe data processing tools (https://github.com/e-strobel-lab/TECtools). TECprobe visualization tools was used to assemble reactivity matrices and analyze the correlation between replicate data sets (https://github.com/e-strobel-lab/TECprobe_visualization). The compile_sm2_output script was used to assemble the reactivity profiles for intermediate transcripts into matrix format and to normalize the dataset. The normalized reactivity matrices were visualized in heatmap format using GraphPad Prism.

For the *alx* riboswitch, replicate data sets were required to obtain adequate sequencing depth for longer transcripts, as there was a strong bias in representation towards shorter transcripts for the amplified cDNA libraries. Plots with fraction of aligned reads per transcript length and alignment rates for *alx* co-transcriptional TECprobe-VL experiments are presented in SI Fig. 21. Additionally, the presence of two intrinsic termination sites in the *alx* riboswitch sequence (110 and 190 nt) results in a clear drop in sequencing depth for intermediate transcripts past these sites (SI Fig. 19A). For the co-transcriptional DMS-MaP libraries, 5-7 independent replicates were pooled per chemical probing condition to increase data quality for longer transcript lengths (see SI Figs 22-25 for individual and concatenated heatmaps). Plots with fraction of aligned reads per transcript length and alignment rates for *alx* equilibrium refolded TECprobe-VL experiments are presented in SI Fig. 26. For the equilibrium refolded *alx* TECprobe-VL experiments, 2 independent replicates were pooled per chemical probing condition to increase data quality for longer transcript lengths (see SI Figs 27-30 for individual and concatenated heatmaps). To further increase sequencing depth when necessary, multiple independent PCR amplifications were performed with the same cDNA library and sequenced; reads were subsequently pooled together for data processing.

For the *mntP* riboswitch, two replicate data sets were concatenated and analyzed together after assessing the correlation between replicates using the plot_cotrans_correlation script available in TECprobe visualization tools. Plots with fraction of aligned reads per transcript length and alignment rates for co-transcriptional *mntP* TECprobe-VL experiments are presented in SI Fig. 31 Plots with fraction of aligned reads per transcript length and alignment rates for equilibrium refolded *mntP* TECprobe-VL experiments are presented in SI Fig. 32. To further increase sequencing depth when necessary, multiple independent PCR amplifications were performed with the same cDNA library and sequenced; reads were subsequently pooled together for data processing.

### *In vitro* synthesis of full-length riboswitch RNAs with T7 RNAP and cleanup for SHAPE-MaP and DMS-MaP

RNA for *in vitro* SHAPE/DMS-MaP was synthesized using the HiScribe T7 High Yield RNA Kit (New England Biolabs). 100 µL reactions were assembled according to manufacturer instructions and incubated at 37 °C overnight. RNA was purified and concentrated using the RNA Clean and Concentrator kit (Zymo Research). For the *alx* riboswitch and *alx* aptamer, RNAs were gel-purified to isolate full-length transcripts away from premature transcription termination products using a denaturing PAGE (8% acrylamide, 8 M urea, 0.5x TBE (Fisher Scientific)). For gel-purification, RNAs were diluted 1:5 in buffer and mixed with an equal volume 2x formamide loading dye (95% formamide, 0.5 mM EDTA, 0.025% bromophenol blue, 0.025% xylene cyanol), heated at 95 °C for 2 mins, and loaded on a pre-heated gel. Gel was run at 50 W, stained with 1x SYBR Green II RNA Gel Stain (Invitrogen) in 1X TBE for 10 mins, and visualized using a Edvotek TruBlu 2 Blue Transilluminator. Bands corresponding to full-length RNA were excised and eluted in 10 mL TE buffer (10 mM Tris pH 8.0, 1 mM EDTA pH 8.0) at 4 °C overnight. The eluted solution of RNA was spun to pellet acrylamide, and RNA was concentrated using an Amicon Ultra Centrifugal filter (3 kDa molecular weight cutoff). RNAs were ethanol precipitated, stored at -20 °C, and resuspended in RNase-free H_2_O. For the *mntP* riboswitch, there are no terminated transcripts, eliminating the need for gel purification. After transcription and cleanup, *mntP* RNAs were ethanol precipitated, stored at - 20 °C, and resuspended in RNase free H_2_O.

### SHAPE-MaP and DMS-MaP on the full-length equilibrium refolded *alx* and *mntP* riboswitches

5 pmol of RNA in added RNase-free H_2_O to a volume of 6 µL was heat denatured for 2 mins at 95 °C, cooled at 4 °C for 2 mins, and then 3 µL buffer was added for a final volume of 9 µL. The final buffer concentration was 100 mM HEPES, 100 mM NaCl, and 5 mM MgCl_2_ at either pH 7.2 or pH 8.4 and supplemented with 1 mM MnCl_2_ where indicated. RNA was incubated at 37 °C for 20 mins to allow refolding. For modification with 1M7, 9 µL of RNA was mixed with 1 µL of 100 mM 1M7 (“+” modified sample) or 1 µL DMSO (“-” negative control), incubated for 75 s at 37 °C, and placed on ice. For the denatured control, 5 pmol RNA (3 µL) was mixed with 5 µL formamide and 1 µL 10x denaturing buffer (500 mM HEPES, 40 mM EDTA) at pH 7.2 or 8.4. RNA was denatured for 1 min at 95 °C, mixed with 1 µL of 100 mM 1M7, incubated at 95 °C for 1 min, and placed on ice. All RNAs were purified and eluted in 50 µL of RNase-free H_2_O using MicroSpin G-25 columns (Cytiva) and stored at -20 °C. For modification with DMS, 9 µL of RNA was mixed with 1 µL of 6.5% v/v DMS (“+” modified sample) or 1 µL ethanol (“-” negative control), incubated for 5 minutes at 37 °C, quenched with 2.4 µL of 30% v/v 2-mercaptoethanol solution at 37 °C for 1 min before placing on ice. For the denatured control, 5 pmol RNA (3 µL) was mixed with 5 µL formamide and 1 µL 10x denaturing buffer (500 mM HEPES, 40 mM EDTA) at pH 7.2 or 8.4. Then RNA was denatured for 1 min at 95 °C, mixed with 1 µL of 6.5% v/v DMS, incubated at 95 °C for 1 min, quenched with 2.4 µL of 30% 2-mercaptoethanol solution at 37 °C for 1 min before placing on ice. All RNAs were purified and eluted in 50 µL of RNase-free H_2_O using MicroSpin G-25 columns (Cytiva) and stored at - 20 °C.

Reverse transcription (RT) of 10 µL of RNA was performed by annealing 1 µL RT primer (0277) for 5 min at 65 °C. Then, 8 µL RT buffer was added (50 mM Tris-Cl pH 8, 75 mM KCl, 10 mM DTT, 0.5 mM dNTPs, and MnCl_2_ added to 15 mM before use). Samples were incubated for 2 min at 42 °C before introducing Superscript II reverse transcriptase (Thermo Fisher Scientific). RT reactions were incubated for 3 h at 42 °C. The reverse transcriptase was inactivated for 15 min at 70 °C before storing the reactions at 4 °C. cDNA was purified and eluted in 50 µL of RNase-free H_2_O using MicroSpin G-25 columns (Cytiva) and stored at -20 °C. The first PCR (step 1) amplified cDNA through 20 amplification cycles using primers 0278 and 0279. Excess primer was removed using the Monarch PCR & DNA Cleanup Kit (New England Biolabs). The second PCR (step 2) further amplified the first PCR amplicon through an additional 10 cycles. The step 2 PCR used primers 0149 and an RPI indexing primer (0163-0178 and 0235-0248). Step 2 PCR was cleaned up using Agencourt AMPure XP beads (Beckman Coulter), and final DNA concentration was measured using the Invitrogen Quant-IT dsDNA High-Sensitivity HS Assay kit quantified with a BioTek Synergy H1 Multimode Microplate Reader before pooling libraries for sequencing. Sequencing of these pooled libraries was performed by Novogene Co. on a NovaSeq 6000 system with 100M 150 PE reads. Sequencing data for SHAPE-MaP and DMS-MaP experiments is publicly available in the Sequencing Read Archive (SRA) under project number PRJNA1197522 and BioSamples are tabulated in Supplementary Table 8.

### RNA structure prediction and visualization

RNA structure prediction for co-transcriptionally probed riboswitch folding intermediates was performed using the RNAstructure V6.4(29) Fold command with default settings, adding the DMS outputs as a constraint, and forcing the 3’-most 11 to 14 nt to be single-stranded to account for the *E. coli* RNAP footprint. The set of minimum free-energy structures were visualized in VARNA(30) to assess agreement with the DMS probing data, and structure diagrams were colored in accordance with DMS reactivity. Additionally, existing X-ray crystal structures for Mn-sensing aptamer domains were visualized with UCSF Chimera(31) and used to guide modeling of conserved secondary and tertiary structure motifs within the *alx* and *mntP* aptamers. RNA structure prediction for equilibrium refolded riboswitch folding intermediates was performed as described above; however, the 3’ most nt were not kept single-stranded since the intermediate RNA transcripts were extracted from stalled transcription elongation complexes before DMS probing.

The denatured and equilibrium-refolded full-length *mntP* riboswitch*, alx* riboswitch, and *alx* aptamer RNA structures were predicted using the RNAStructure V6.4(29) Fold command with default settings, adding the SHAPE or DMS outputs as a constraint. The minimum free energy structures were visualized in VARNA(30) and colored in accordance with nucleotide reactivity.

### β-galactosidase assays

Overnight culture of each strain was diluted 1:100 into LBK with appropriate antibiotic at either pH 6.8 or pH 8.4, with or without 1 mM supplemented MnCl_2_. The cultures were grown to mid-log phase and supplemented with 1 mM IPTG where necessary for expression of T7 RNAP.

β-galactosidase assays were performed using the Miller method(32). Briefly, bacterial growth was measured using OD_600_ before cultures were placed on ice. 100 µL cells were lysed via vortexing in 900 µL sodium phosphate buffer (100 mM sodium phosphate pH 7, 10 mM KCl, 2 mM MgSO_4_, 38 mM 2-mercaptoethanol), 20 µL 0.1% SDS, and 20 µL chloroform. Substrate *O*-nitrophenyl-β-D-galactopyranoside (ONPG) was added and the reaction was stopped with 1 M Na_2_CO_3_ when a yellow color appeared, or after 35 min. After centrifuging 5 min at 16, 000 g, absorbance was measured at 420 nm (A_420_) to calculate activity using Miller units. Each culture was tested in duplicate and A_420_ values were averaged. Each final reported value is an average of three replicates with standard deviation.

### Sequence analysis for *yybP-ykoY* riboswitches

The sequences for *yybP-ykoY* riboswitches were downloaded from the Rfam database (Rfam reference number RF00080) and aligned using the Kalign multi sequence alignment program (https://www.ebi.ac.uk/jdispatcher/msa). The sequence alignment and consensus L3 sequence was visualized using Jalview(33). The *yybP-ykoY* riboswitch sequences found within the Enterobacteriales order were accessed from both the Rfam database and from *yybP-ykoY* regulog collection (https://regprecise.lbl.gov/collection_rfam.jsp?riboswitch_id=13). These sequences were aligned with the Muscle multi sequence alignment program (https://www.ebi.ac.uk/jdispatcher/msa). The sequence alignment and consensus L3 sequence was visualized using Jalview(33).

## RESULTS

### The Mn-sensing aptamers of the *alx* and *mntP* riboswitches differ structurally

In Fig. 2A, we present secondary structure models of the *alx* and *mntP* riboswitch aptamer domains based on prior experimental data for the full-length riboswitches(10, 18), which we updated to reflect the overall aptamer architecture and conserved base-pairs identified by X-ray crystal structures(6, 12, 21). The model of the *mntP* aptamer in Fig. 2A is supported by an X-ray crystal structure of the aptamer in the apo state (PDB ID: 4Y1M)(6). Since no detailed structural information exists for the *alx* aptamer, we mapped its secondary structure with DMS/SHAPE-MaP using the traditional approach of *in vitro* synthesis by T7 RNA polymerase (RNAP) followed by denaturation and refolding.

*pH and Mn effect on the alx aptamer structure.* The aptamer RNA was refolded at either neutral (7.2) or alkaline pH (8.5), consistent with pH values used in published work for DMS probing, *in vitro* transcription, and gene reporter fusion experiments(18, 19, 34). The aptamer RNAs were probed with either 1-methyl-7-nitroisatoic anhydride (1M7) or DMS. The resulting secondary structure models of the *alx* aptamer agree with the published 2D predictions(18) and 3D structure of an *L. lactis* chimera with the *alx* L3 sequence(21) (Fig. 2B and SI Fig. 1A-C). Comparison of the nucleotide (nt) reactivities as a function of refolding pH revealed clear flexibility differences in L3 and L1, with multiple nt (e.g., A37 and A11) showing decreased reactivity towards 1M7 or DMS at alkaline pH (Fig. 2C).

We next refolded the *alx* aptamer RNA at either neutral (7.2) or alkaline pH (8.5) in the presence of 1 mM MnCl_2_ and probed with either DMS or 1M7. While the total physiological [Mn] (free and bound) in *E. coli* is estimated to be 15-21 µM(35–37), we chose a saturating concentration of ligand to maximize the ligand-induced shift in the RNA conformational ensemble, consistent with prior chemical probing experiments on other riboswitches(27, 38). MgCl_2_ was present at 5 mM in the RNA refolding buffer, thus Mg remained in excess of Mn, as is the case *in vivo*(39). We observed L3 generally becomes less reactive towards 1M7 in the presence of Mn, consistent with reduced flexibility upon Mn binding (SI Fig. 1D). Interestingly, in the presence of Mn, pH-dependent reactivity changes for multiple L1/L3 nt are abolished (e.g., A37 and A11, Fig. 2C), supporting that these nt are buried in the metal-sensing core with Mn and consistent with published Mn-bound aptamer structures(6, 12, 21) (see Supplementary Note 1 and SI Fig. 1E for additional analysis).

*Structural comparison of the alx and mntP aptamers.* Despite belonging to the same family, the *alx* and *mntP* riboswitch aptamers harbor intriguing structural differences. First, the *alx* aptamer contains an intrinsic termination site in the P4 helix(34), which occurs before the metal-sensing pocket forms (Fig. 2A). The *alx* aptamer lacks the P2 helix, resulting in a three-way junction (3WJ) rather than a 4WJ in the *mntP* aptamer and all other aptamers studied by X-ray crystallography. Lastly, the L3 sequences in *alx* and *mntP* metal-sensing cores differ (Fig. 1C), raising the possibility that L3 tunes Mn affinity of the aptamer.

### Transcription by native RNA polymerase is required for *alx* and *mntP* riboswitch-mediated translational regulation *in vivo*

We next wondered whether *alx* and *mntP* riboregulation is kinetically controlled. To test this, we constructed *alx* and *mntP* translational *lacZ* reporters cloned with either the T7 promoter or native *alx/mntP* promoter into single copy plasmids (Fig. 2D). The effect of extracellular alkaline pH or elevated [Mn] on gene expression was measured by β-galactosidase assays in either an *E. coli* strain (MC4100) lacking *alx* (τι*alx*), to preclude negative autoregulation(19), or the τι*alx* strain with T7 RNAP expressed from the *lac* promoter (chromosomally encoded, τι*alx*T7). The plasmids containing the native promoter translational reporters were transformed into the τι*alx* strain and the plasmids containing the T7 promoter translational reporters were transformed into the τι*alx*T7 strain. The strains were cultivated in i) neutral pH (LBK pH 6.8) or alkaline pH (LBK pH 8.4) media, and ii) neutral pH (LBK pH 6.8) media with supplemented MnCl_2_ .

We observed the expected changes in *alx* and *mntP* translation in response to pH and supplemented MnCl_2_ when reporter fusions were transcribed from their native promoters(19) (Fig. 2E-F). While only *alx* translation was induced by alkaline pH, translation of both genes increased with supplemented MnCl_2_ (Fig. 2E-F). Strikingly, transcription by T7 RNAP abolished the response to alkaline pH and/or Mn (Fig. 2E-F), demonstrating that faster transcription of the *alx* and *mntP* riboswitches (∼220 nt/s; T7 RNAP(40) vs. ∼20-90 nt/s; *E. coli* RNAP(41)) prevents riboswitching in response to alkalinization and/or elevated [Mn] in the cytosol. In sum, these data establish that transcription by *E. coli* RNAP is required for ligand-coupled structural rearrangements in the *alx* and *mntP* expression platforms.

### Analysis of the *mntP* aptamer folding during transcription using co-transcriptional chemical probing

To investigate both pH and Mn-mediated structural changes in the *alx* and *mntP* riboswitch folding intermediates, we performed co-transcriptional chemical probing (<u>v</u>ariable length <u>T</u>ranscription <u>E</u>longation <u>C</u>omplex RNA structure <u>prob</u>ing TECprobe-VL method, see SI Fig. 2 for construct designs)(27). Co-transcriptional chemical probing experiments were performed at both neutral (7.2) and alkaline pH (8.5) ± 1 mM MnCl_2_ (referred to as “+Mn” or “–Mn” condition, respectively) with 10 mM MgCl_2_ present to prevent compromised transcription fidelity from Mn-coordination by RNAP, as observed for DNA polymerases(42) and reverse transcriptases(43). Since published work established that pH≥ 8 renders all 4 nucleobases accessible to DMS modification(44), we first present data for *alx* and *mntP* DMS-probed folding intermediates at pH 8.5.

*The two legs of the Mn aptamer fold differently.* We initially mapped *mntP* aptamer folding transitions and determined the specific transcript lengths where helices in the aptamer become stably folded (Fig. 3A-B and SI Fig. 3). P3.1, P3.2, and P4 in the left leg of the aptamer fold first, as detected with coordinated drops in reactivity of nucleotides that are base-paired in these helices (Fig. 3C). In stark contrast, the right leg of the aptamer folds into transient, low-melting temperature (T_m_) stem-loops that form and resolve until the synthesis of the aptamer is completed (∼nt 124, assuming a 14-nt RNAP footprint, Fig. 3B). While the 5’ RNA stretch for these aptamer elements (L1, P1.1, and P1.2) is transcribed early on, their 3’ pairing partner does not emerge from RNAP until late in transcription. Thus, the weak stem-loop intermediates may prevent RNA from being trapped in stable but not biologically relevant folds, consistent with other riboswitches(16, 27). P2 in the right leg folds towards the end of aptamer transcription, even though the sequence encoding P2 emerges at transcript length ∼50 nt, indicating that folding of the entire right leg occurs at the end of aptamer synthesis (Fig. 3C).

**Figure 3.**
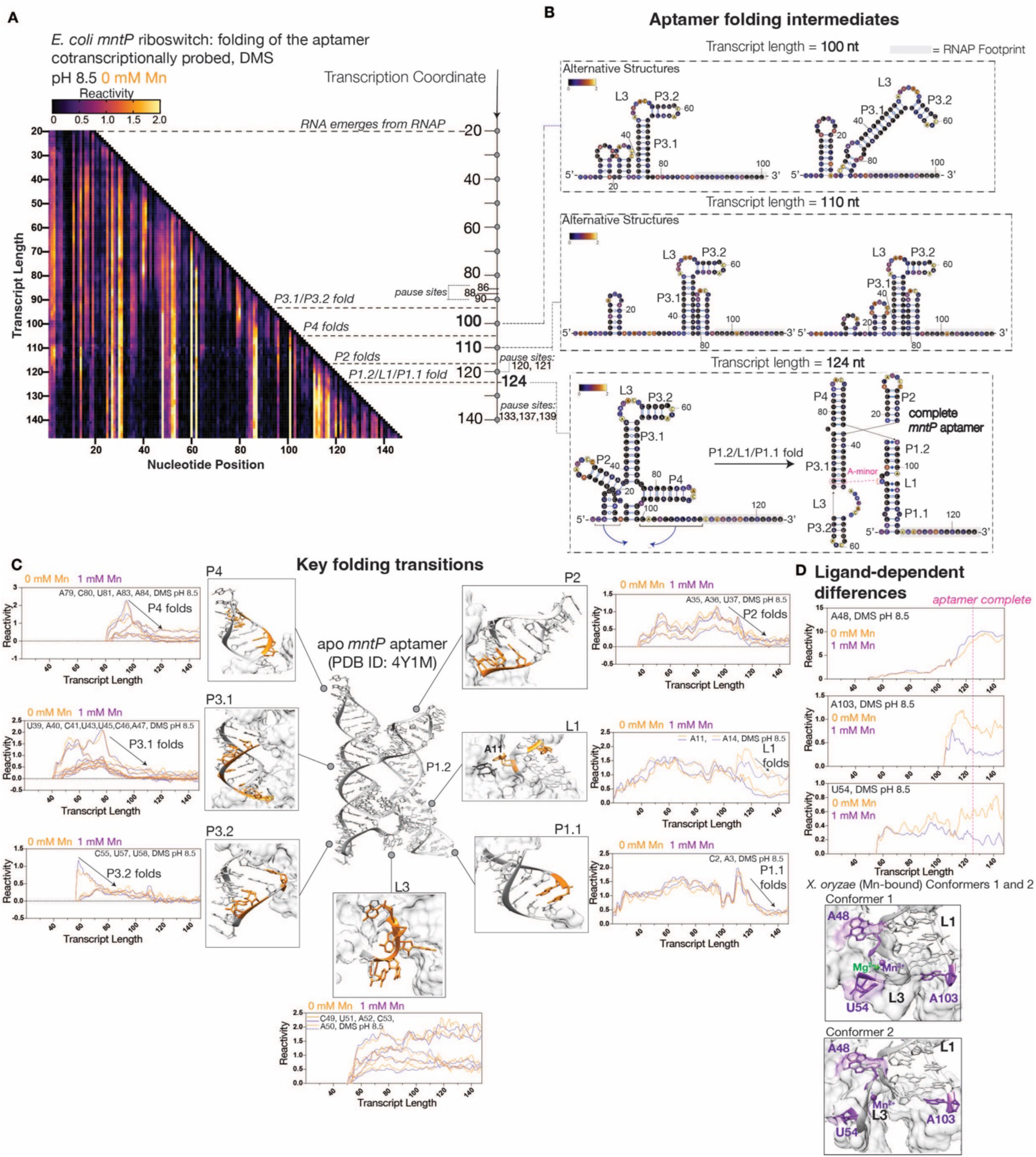
Co-transcriptional DMS probing of the *mntP* riboswitch aptamer at alkaline pH. **A.** TECprobe-VL DMS reactivity matrix for the *mntP* aptamer at pH 8.5. Reactivities shown were normalized with a single, whole-dataset calculated normalization factor. Data are from two independent replicates that were concatenated and analyzed together. **B.** Secondary structures of *mntP* aptamer folding intermediates colored by DMS reactivity of the indicated transcript at pH 8.5. **C.** Transcript length-dependent reactivity changes showing folding transitions of the *mntP* aptamer at pH 8.5 ± 1 mM Mn. DMS data for 0 mM Mn (orange) and 1 mM Mn (purple) conditions from Fig. 3A and SI Fig. 3, respectively. *E. coli mntP* apo aptamer (PDB ID: 4Y1M) shown with labeled loops and helical domains. Nt highlighted in orange correspond to the nt positions with reactivities plotted in the adjacent graph. **D.** Plots show transcript length- and Mn-dependent reactivity changes in *mntP* aptamer folding intermediates at pH 8.5 ± 1 mM Mn. DMS data for 0 mM Mn (orange) and 1 mM Mn (purple) conditions from Fig. 3A and SI Fig. 3, respectively. Vertical dotted line marks completion of aptamer folding. *X. oryzae* aptamer metal-sensing core (PDB ID: 6N2V) with Mg^2+^ (green sphere) and Mn^2+^ (purple sphere) highlighted. Conformer 1 shows Mn^2+^ fully bound and Conformer 2 shows Mn^2+^ partially bound. Nt highlighted in purple correspond to nt reactivity plots above.

*RNA folding intermediates sample for Mn.* Surprisingly, we observed Mn-dependent differences in reactivity of several nucleotides *<u>before</u>*complete *mntP* aptamer has been synthesized (Fig. 3D). For example, A48, the first nucleotide in the L3 loop, demonstrates markedly high reactivity ±Mn, with its reactivity trending higher +Mn starting at transcript length ∼115 nt – before the aptamer synthesis finished ∼124 nt (Fig. 3D). We hypothesize that this is indicative of an RNA folding intermediate sampling divalent metal ions in solution in search for Mn, with its L3 and partially formed L1/P1.1. Such partial binding of Mn was observed in a crystal structure and molecular dynamics (MD) simulations of the full-length *X. oryzae* aptamer, referred to as “conformer 2”, which lacks the ordered stacking pattern in L3 and contains a partially dehydrated Mn in the M_B_ site(12) (Fig. 3D). The partially dehydrated Mn lacks half of the contacts seen in “conformer 1” (coordinated to fully dehydrated Mn); however, the inner sphere contact between the phosphate of the first nucleotide in L3 and the metal is observed (A48 in *E. coli mntP* aptamer vs. A46 in *X. oryzae* aptamer)(12). Therefore, A48 can contact a partially dehydrated Mn before the full aptamer is synthesized, explaining its Mn-dependent reactivity differences in *mntP* aptamer folding intermediates. U54 in L3 and A103 in L1 also show Mn-dependent reactivity differences in aptamer intermediates, with their reactivities trending higher –Mn (Fig. 3D). These reactivity differences can be explained by greater conformational flexibility in the metal-binding site (see Fig. 3D for U54 and A103 positioning in *X. oryzae* conformers 1 and 2). Additionally, we observed multiple RNAP pauses preceding *mntP* aptamer completion that are enhanced by the presence of Mn and may mediate partial Mn binding/ sampling of divalents in the environment by the RNA (see Supplementary Note 2 and SI Figs. 4-5 for further analysis).

**Figure 4.**
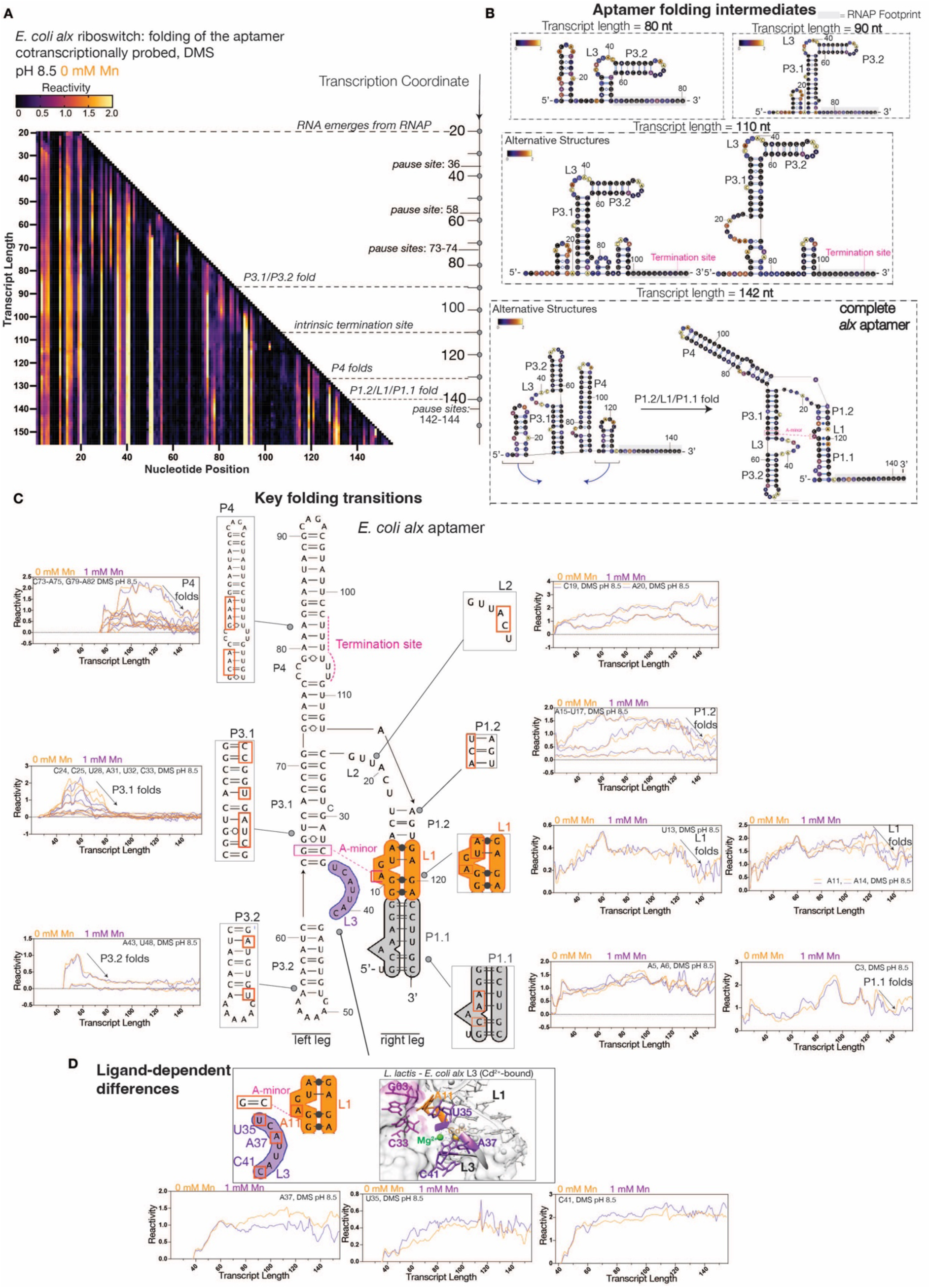
Co-transcriptional DMS probing of the *alx* riboswitch aptamer at alkaline pH. **A.** TECprobe-VL DMS reactivity matrix for the *alx* aptamer at pH 8.5. Reactivities shown were normalized with a single, whole-dataset calculated normalization factor. Data are from seven independent replicates that were concatenated and analyzed together. **B.** Secondary structures of *alx* aptamer folding intermediates, which are colored by the DMS reactivity of the indicated transcript at pH 8.5. **C.** Transcript length-dependent reactivity changes showing folding transitions of the *alx* aptamer at pH 8.5 ± 1 mM Mn. DMS data for 0 mM Mn (orange) and 1 mM Mn (purple) conditions from Fig. 4A and SI Fig. 7, respectively. Secondary structure model of *alx* aptamer with labeled structural domains. L1 (orange), L3 (purple), P1.1 (gray), and intrinsic termination site (pink) are highlighted. Nt boxed in red correspond to adjacent nt reactivity plots. **D.** *L. lactis* Mn-sensing aptamer with *E. coli alx* L3 sequence (PDB ID: 6CC1). L3 nt (purple), cross-helix A-minor nt: A11 (orange) and G63-C33 (magenta), Mg^2+^ (green sphere), and Cd^2+^ (yellow sphere) are highlighted. Plots show transcript length- and Mn-dependent reactivity changes in *alx* aptamer folding intermediates at pH 8.5 ± 1 mM Mn. DMS data for 0 mM (orange) and 1 mM (purple) conditions from Fig. 4A and SI Fig. 7, respectively.

**Figure 5.**
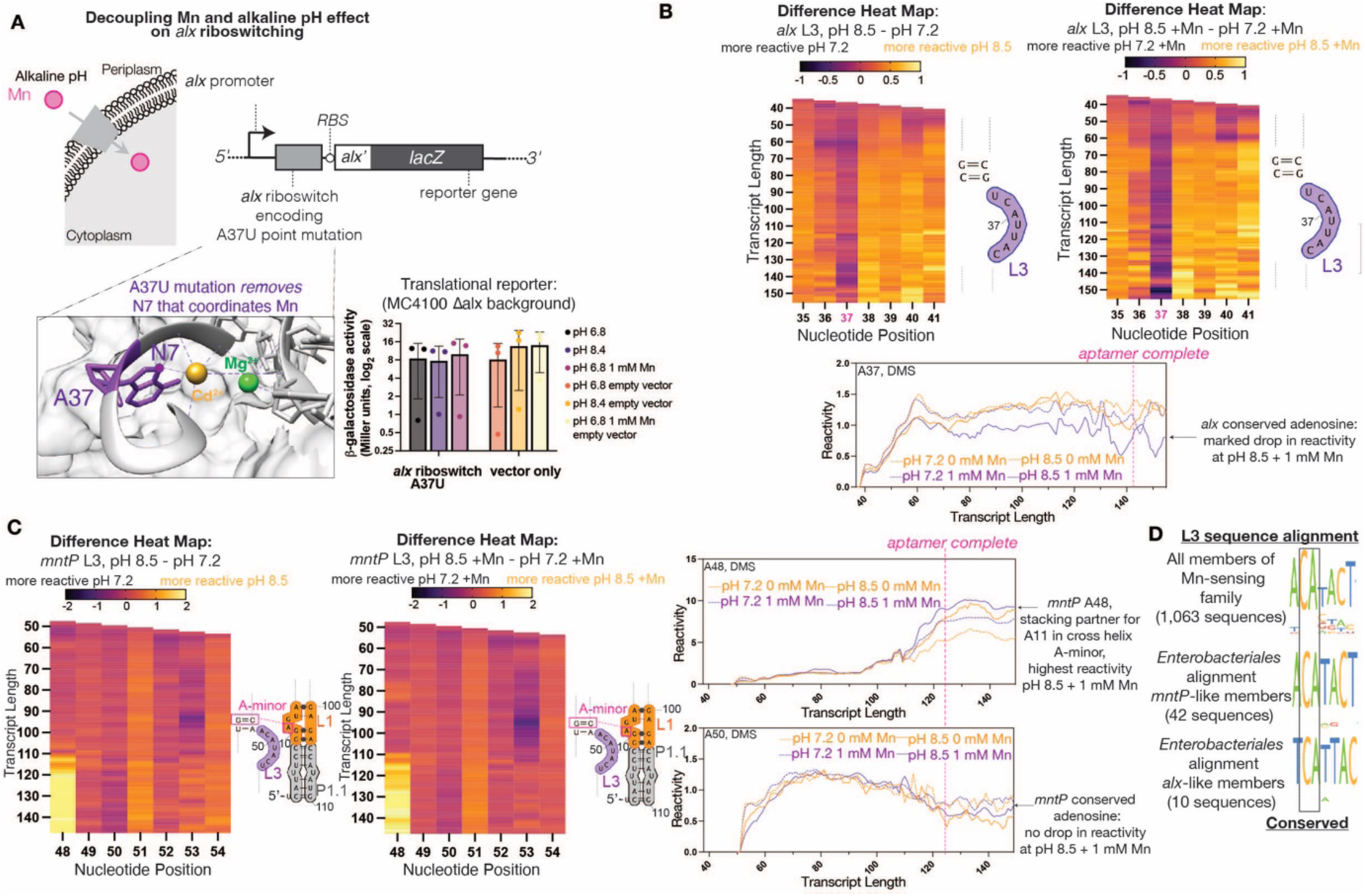
Decoupling the effects of alkaline pH and Mn on *alx* riboswitching *in vivo* and pH-dependent reactivity changes in the divergent *alx* and *mntP* L3 sequences. **A.** (Top panel) *E. coli* cell with increased cytosolic [Mn] resulting from an alkaline pH challenge (left) and construct design for *alx* translational reporter (right). (Bottom panel) *L. lactis* Mn-sensing aptamer with *E. coli alx* L3 sequence (PDB ID: 6CC1, left). Conserved A37 (purple), Mg^2+^ (green sphere), and Cd^2+^ (yellow sphere) are highlighted. (Bar graph) Loss of *alx* translational response to either Mn or pH upon mutation of the Mn-coordinating adenosine. **B-C**. Reactivity difference heatmaps and reactivity changes in aptamer intermediates for *alx* L3 nt (**B**) and *mntP* L3 nt (**C**) during aptamer folding at pH 7.2 vs. 8.5 ± 1 mM Mn. *alx* L3 and *mntP* L3 (purple), L1 (orange), L3 (purple), P1.1 (gray), and cross-helix A-minor (pink) are highlighted. DMS data for 0 mM Mn (orange) and 1 mM Mn (purple) conditions from Fig. 4A, SI Fig. 7, and SI Fig. 9B (**B**, *alx*) and Fig. 3A, SI Fig. 3, and SI Fig. 9A (**C**, *mntP*). Vertical dotted line marks completion of aptamer folding. **D.** L3 consensus sequences from *yybP-ykoY* family members in the Rfam database (top) and *mntP*-like (middle)/ *alx*-like (bottom) members in the *Enterobacteriales* order.

To identify the intermediates whose folding is truly controlled by transcription kinetics vs. thermodynamic stability of the folds, we applied a modified TECprobe-VL workflow wherein intermediate RNA transcripts are extracted from the stalled transcription elongation complexes and equilibrium refolded before DMS probing. These experiments identified clear structural differences between co-transcriptionally folded and equilibrium refolded *mntP* intermediates and provided additional evidence of divalent sampling by aptamer folding intermediates (see Supplementary Note 3 and SI Fig. 6 for further analysis).

**Figure 6.**
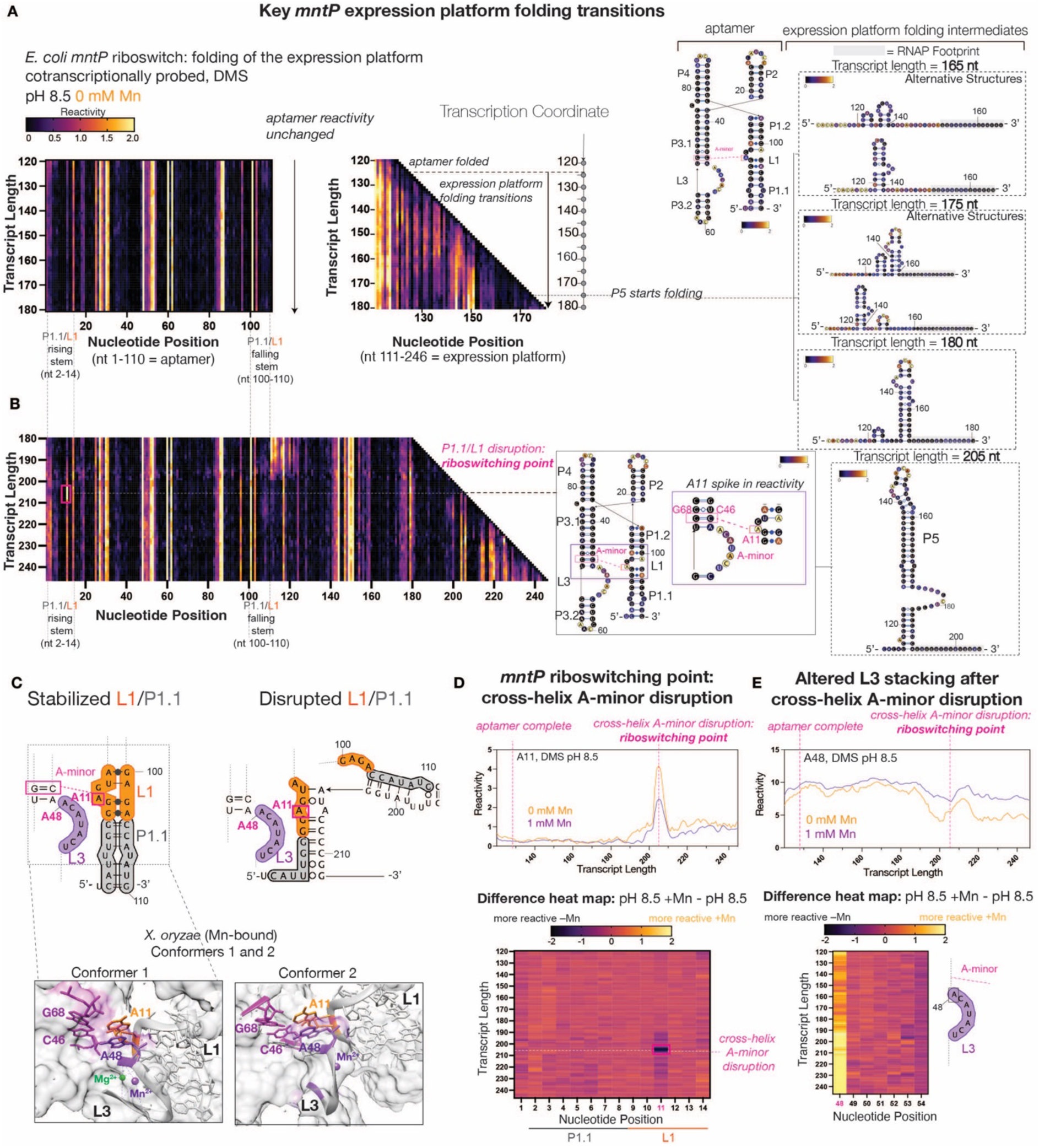
Co-transcriptional DMS probing of the *mntP* riboswitch during expression platform folding. **A-B**. (Left panel) TECprobe-VL DMS reactivity matrices of *mntP* riboswitch at pH 8.5. Reactivities shown were normalized with a single, whole-dataset calculated normalization factor. Data are from two independent replicates that were concatenated and analyzed together. (Right panel) Secondary structures of *mntP* expression platform folding intermediates colored by DMS reactivity of the indicated transcript at pH 8.5. **C.** (Top panel) Truncated secondary structure models of *mntP* riboswitch with a stabilized L1/P1.1 (Mn-bound) or disrupted L1/P1.1 (Mn-unbound). L1(orange), L3 (purple), P1.1 (gray), and cross-helix A-minor interaction (pink) are highlighted. (Bottom panel) *X. oryzae* aptamer metal-sensing core (PDB ID: 6N2V) with Mg^2+^ (green sphere) and Mn^2+^ (purple sphere) highlighted. Conformer 1 shows Mn^2+^ fully bound and Conformer 2 shows Mn^2+^ partially bound. A11 (orange) and A48 (purple) are highlighted. **D-E.** (Top panels) Transcript length- and Mn-dependent reactivity changes in A11 (**D**) and A48 (**E**) showing cross-helix A-minor disruption. (Bottom panels) Reactivity difference heatmaps for P1.1/L1 (**D**) and L3 (**E**) at pH 8.5 ± 1 mM Mn. DMS data for 0 mM and 1 mM Mn conditions from Fig. 6A-B and SI Fig. 12A, respectively. Vertical or horizontal dotted lines mark indicated RNA folding transitions.

### Analysis of the *alx* aptamer folding during transcription using co-transcriptional chemical probing

We next investigated the *alx* aptamer folding transitions at pH 8.5 (Fig. 4A-B and SI Fig. 7). Here, P3.1, P3.2, and P4 in the left leg fold first as the aptamer is extended from ∼80 nt to ∼125 nt (Fig. 4B-4C). Like in the *mntP* aptamer, the 5’ end of the right leg folds into transient, low-T_m_ stem-loops that resolve once the full-length aptamer RNA has emerged from RNAP (Fig. 4B), highlighting a potentially conserved mechanism of the *yybP-ykoY* aptamer folding.

**Figure 7.**
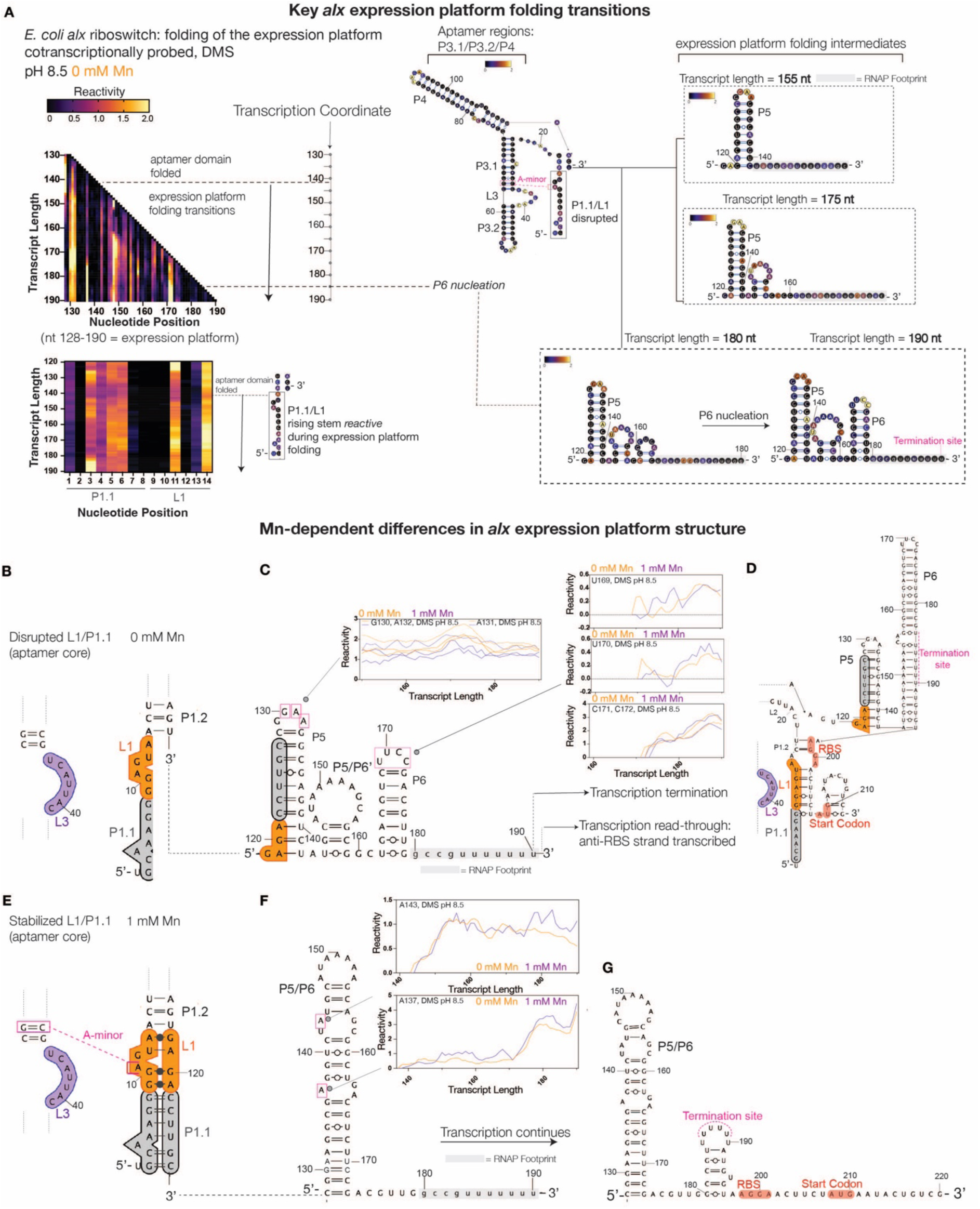
Co-transcriptional DMS probing of the *alx* riboswitch during expression platform folding. **A.** (Left panel) TECprobe-VL DMS reactivity matrix of the *alx* expression platform (top) and L1/P1.1 rising stem (bottom) at pH 8.5. Reactivities shown were normalized with a single, whole-dataset calculated normalization factor. Data are from seven independent replicates that were concatenated and analyzed together. (Right panel) Secondary structures of *alx* folding intermediates colored by DMS reactivity of the indicated transcript at pH 8.5. **B.** Truncated secondary structure model of the *alx* aptamer with a disrupted L1/P1.1 (Mn- unbound). L1 (orange), L3 (purple), and P1.1 (gray) are highlighted. **C.** Secondary structure model of *alx* expression platform at transcript length 190 nt –Mn. L1 falling stem (orange) and P1.1 falling stem (gray) paired in P5 are highlighted. Adjacent to nt boxed in pink are plots with their respective transcript-length and Mn-dependent reactivity changes. DMS data for 0 mM (orange) and 1 mM Mn (purple) conditions are from Fig. 7A and SI Fig. 15A, respectively. **D.** Secondary structure model of full-length, translationally inactive *alx* expression platform with same aptamer color scheme as in panels A-B. RBS/start codon (red) and P6 intrinsic terminator stem (pink) are highlighted. **E.** Truncated secondary structure model of the *alx* aptamer with a stabilized L1/P1.1 (Mn-bound) with same aptamer color scheme as in panel A. Cross-helix A-minor motif (pink) is highlighted. **F.** Secondary structure model of *alx* expression platform at transcript length 190 nt +Mn. Adjacent to nt boxed in pink are plots with their respective transcript-length and Mn-dependent reactivity changes. DMS data for 0 mM (orange) and 1 mM Mn (purple) conditions are from Fig. 7A and SI Fig. 15A, respectively. **G.** Secondary structure model of full-length, translationally active *alx* expression platform with same aptamer color scheme as in panel D. RBS/start codon (red) and intrinsic termination site in inactive conformer (pink) are highlighted.

*L3 flexibility differs between alx and mntP aptamers.* Like with *mntP*, we captured Mn-dependent reactivity differences for L3 in *alx* aptamer folding intermediates (Fig. 4D). For example, the reactivity of U35, the first nt in the *alx* L3, trends higher +Mn starting at transcript lengths as short as ∼50 nt (Fig. 4D). This trend mimics that seen with the first nt in *mntP* L3, A48, with a few key differences. First, the absolute reactivity values ±Mn are greater for A48 in *mntP* L3 (Fig. 3D). Second, the increased reactivity +Mn occurs much earlier for *alx* aptamer intermediate transcript lengths (∼50 nt for *alx* vs. ∼115 nt for *mntP*). The absolutely conserved adenosine in *alx* L3 that directly coordinates Mn (A37) is more reactive –Mn for aptamer folding intermediates starting at transcript length ∼70 nt (Fig. 4D). The higher A37 reactivity – Mn is consistent with greater L3 flexibility in this condition and further supports that the L3 loop can begin sampling for divalents before the aptamer domain is fully folded. Curiously, the absolutely conserved adenosine in *mntP* L3 (A50) does not demonstrate this Mn-dependent reactivity difference (Fig. 3C), pointing to a difference in L3 dynamics between *alx* and *mntP* aptamer folding intermediates. Similar to *mntP*, we observed clear structural differences between co-transcriptionally folded and equilibrium refolded *alx* intermediates (see Supplementary Note 4 and SI Fig. 8 for further analysis).

**Figure 8.**
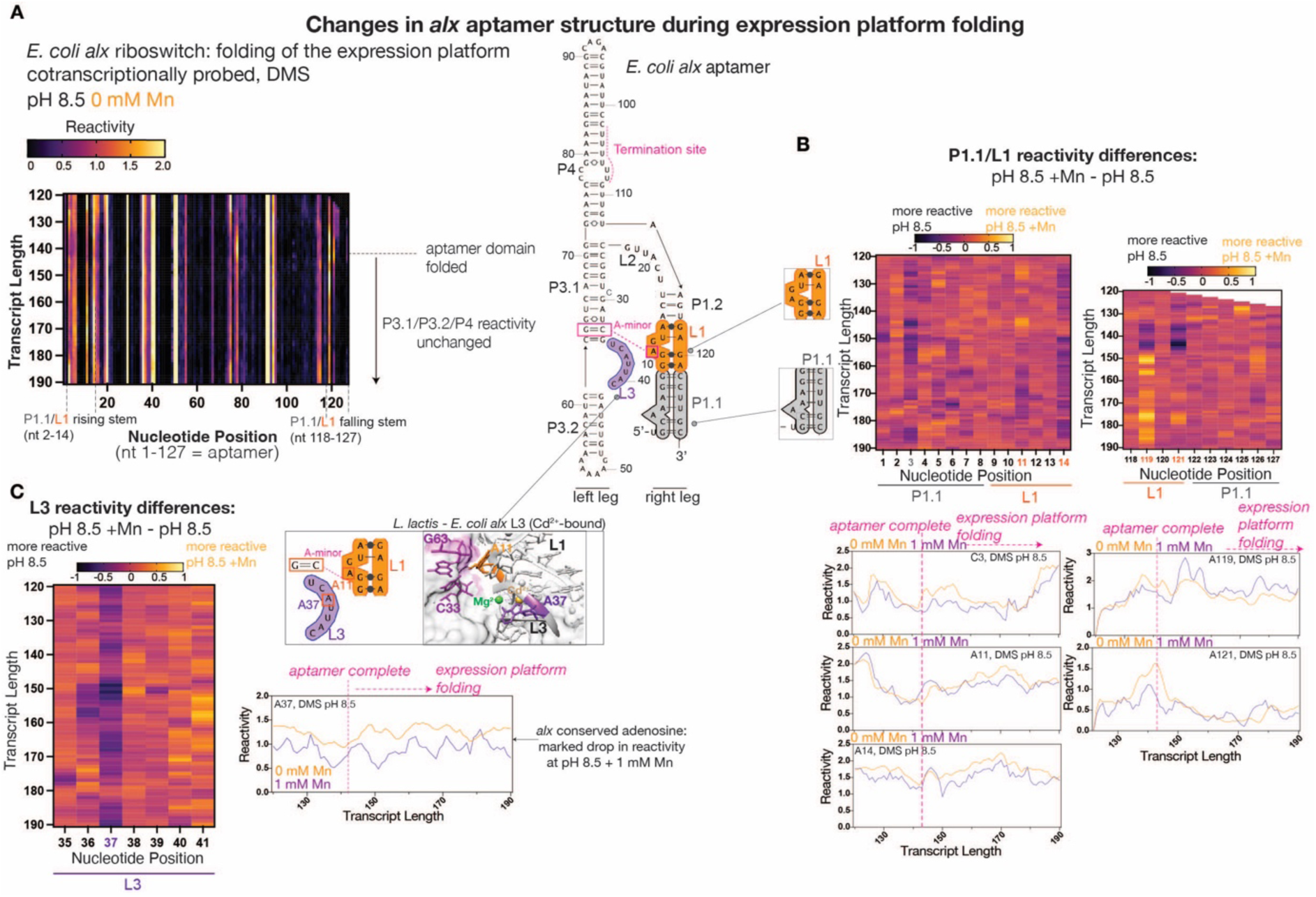
Changes in the *alx* aptamer during expression platform folding. **A.** (Left panel) TECprobe-VL DMS reactivity matrix of the *alx* riboswitch aptamer during expression platform folding at pH 8.5. Reactivities shown were normalized with a single, whole-dataset calculated normalization factor. Data are from seven independent replicates that were concatenated and analyzed together. (Right panel) Secondary structure model of *alx* aptamer with labeled structural domains. L1 (orange), L3 (purple), P1.1 (gray), and intrinsic termination site (pink) are highlighted. **B-C.** Reactivity difference heatmaps for L1/P1.1 (**B**) and L3 (**C**) at pH 8.5 ± 1 mM Mn. Adjacent to heatmaps are transcript length- and Mn-dependent reactivity changes in L1/P1.1 nt (**B**) and L3 nt (**C**). DMS data for 0 mM and 1 mM Mn conditions from Fig. 8A and SI Fig. 15A, respectively. Vertical or horizontal dotted lines/arrows mark indicated RNA folding transitions. *L. lactis* Mn-sensing aptamer with *E. coli alx* L3 sequence (PDB ID: 6CC1) shown in panel B with A37 (purple), cross-helix A-minor nt: A11 (orange) and G63-C33 (magenta), Mg^2+^ (green sphere), and Cd^2+^ (yellow sphere) highlighted.

### Mn binding is a prerequisite for *alx* riboswitch pH response

Early work characterized the *alx* riboswitch as a pH sensor(18); however, recent studies have called this into question(19). Since the intracellular [Mn] increases in *E. coli* at alkaline pH(19), we wondered whether the *alx* riboswitch is actually sensing the increase in [Mn] as a consequence of cytosol alkalinization. To test this, we performed translational *lacZ* reporter assays on the A37U *alx* mutant, where the absolutely conserved adenosine that directly coordinates Mn in L3 is replaced with a uridine. This point mutation was previously shown to ablate Mn-sensing in the *X. oryzae* aptamer(12) making it the best candidate mutation to decouple the effect of Mn and alkaline pH on *alx* riboregulation *in vivo*. Strikingly, the 68-fold increase in *alx* reporter translation observed upon a shift from pH 6.8 to 8.4 for the wild-type riboswitch was abolished by the A37U mutation (Fig. 5A). These data provide the first evidence that the *alx* riboswitch is not a true pH sensor; instead, the riboswitch senses increased cytosolic [Mn], which is an indirect effect of cytosol alkalinization. Published *alx* translational reporter data showed an additive effect of alkaline pH and Mn(19). We thus hypothesize that alkaline pH enhances the affinity of the *alx* aptamer for Mn, rather than alters the *alx* riboswitch folding, unifying observations from early studies on this system(18, 45) with a recently published genetic study(19). A combination of alkaline pH and high [Mn] produced higher *mntP* reporter translation compared to the neutral pH condition with added Mn, suggesting that alkaline pH may also enhance Mn affinity for the *mntP* aptamer, but to a lesser degree than for the *alx* aptamer(19). To explore this intriguing point further, we DMS-probed the *alx* and *mntP* folding intermediates at pH 7.2 ±Mn with the TECprobe-VL method, as described in the following section.

### pH effects on the *alx* and *mntP* aptamer folding: L3

We largely observed the same co-transcriptional folding transitions for the two legs in the *alx* and *mntP* aptamers at pH 7.2 as described for pH 8.5 (SI Fig. 9A-B). Although the *alx* and *mntP* aptamers exhibit the same global folding transitions at pH 7.2, key pH-dependent reactivity differences in their metal-sensing cores emerged. Here, we focused on A and C nucleotide reactivities since G and U nucleotides are protonated at the respective N1 and N3 positions at neutral pH and are thus unreactive towards DMS (pKa ∼ 9.2(46)). First, the absolutely conserved adenosine in the *alx* L3 (A37) exhibits a general Mn-dependent reactivity decrease during aptamer folding at both pH (Fig. 5B). However, the magnitude of this reactivity decrease is clearly pH-dependent, with A37 consistently less reactive at pH 8.5 +Mn compared to other conditions tested (Fig. 5B), suggesting a potentially more stable A37-Mn coordination at alkaline pH. In stark contrast, the absolutely conserved adenosine in *mntP* L3 (A50) shows no drop in reactivity at pH 8.5 +Mn; instead, A50 lacks clear pH-dependent reactivity trends during aptamer folding (Fig. 5C). The first L3 nucleotide in the *mntP* aptamer (A48) is more reactive +Mn vs. –Mn regardless of pH (Fig. 5C), supporting that A48 stacked with A11 in an Mn-bound, ordered metal-sensing core increases its DMS reactivity. Interestingly, the absolute reactivity values for A48 are markedly lower at pH 7.2 ±Mn (Fig. 5C), suggesting that A48 has an upwardly shifted pK_a_ close to physiological pH (∼pH 7.4-7.8 for *E. coli*(23)). We thus hypothesize that deprotonation of A48 at alkaline pH promotes its stacking with A11 in the cross-helix A-minor interaction, favoring docking of the helical legs in the aptamer (see Supplementary Note 5 and SI Figs. 9C-D, 10, and 11 for further analysis of pH-dependent structural changes in *alx/mntP* L1 and L3 in co-transcriptionally folded vs. equilibrium refolded aptamer intermediates).

**Figure 9.**
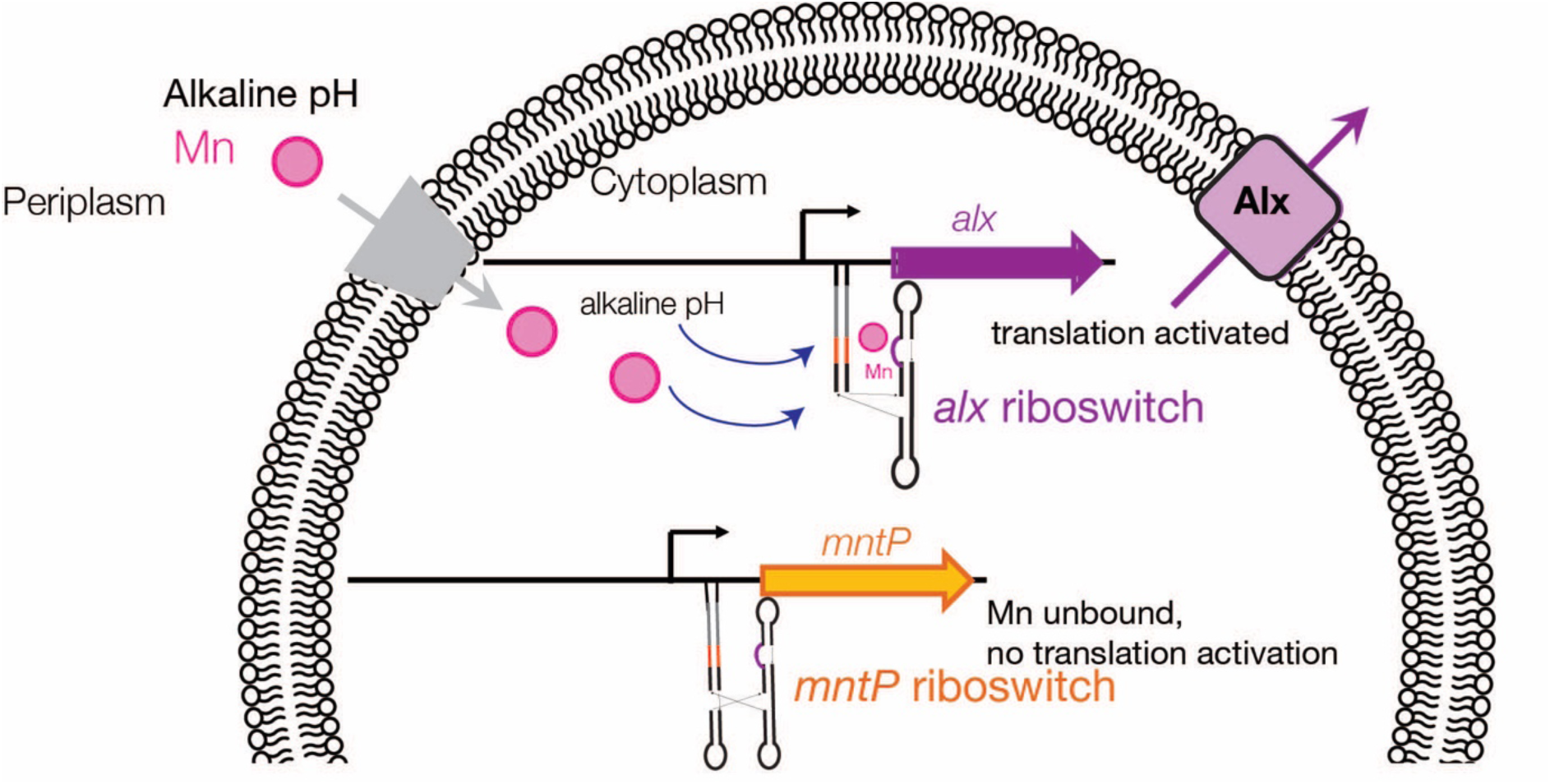
Model for *alx* and *mntP* riboregulation in *E. coli*. Cartoon model of *alx* and *mntP* riboregulation of their respective genes in *E. coli*. At alkaline pH, the cytosolic [Mn] increases, resulting in the *alx* riboswitch turning “on” expression of Alx to prevent toxic accumulation of Mn in the cytosol. The modest increase in cytosolic [Mn] brought about by alkaline pH is insufficient to shift the *mntP* riboswitch towards its translationally active conformation, leaving *mntP* expression “off”.

We wondered whether L3 sequences in *Enterobacteriales yybP-ykoY* family members show a unique trend that could explain the pH-dependent reactivity differences in the *alx* and *mntP* aptamers. We aligned sequences of *yybP-ykoY* riboswitch family members within the *Enterobacteriales* order, revealing two distinctive groups: an *alx*-like group (consensus: UCAUUAC, Fig. 5D) and an *mntP*-like group (consensus: ACAUACU, Fig. 5D). We then compared the above sequences to the *yybP-ykoY* riboswitch family L3 consensus sequence generated from 1, 063 aligned sequences (consensus: ACAUACU, Fig. 5D). Generally, the first nucleotide in L3 is an A – an important position since this nucleotide stacks with the conserved A11 in L1 that forms the cross-helix A-minor interaction. Therefore, this position being strictly a U in *alx*-like members suggests a functional role. Since the N3 of U has a close-to-physiological pKa of ∼9.2(46), it is possible that a shift from neutral to alkaline pH alters its stacking with the conserved A11 and consequently Mn-coupled structural rearrangements in the expression platform.

### Analysis of *mntP* expression platform folding during transcription

*Cross-helix A-minor disruption at the riboswitching point.* The expression platform for the *mntP* riboswitch contains a single helix that begins folding at transcript length ∼175 nt and was detected by coordinated drops in reactivity for its constituent nucleotides (Fig. 6A and Fig. S12A). In the absence of Mn, the aptamer remains stably folded for most of expression platform synthesis until the transcript is ∼205 nt long (Fig. 6B). At this critical decision point – the riboswitching point – the L1 and P1.1 elements of the aptamer are disrupted if no Mn is available, allowing for alternative pairing of P1.1 switch helix in the expression platform (Fig. 6C). The disruption of P1.1 in the absence of Mn is consistent with previously published single-molecule fluorescence experiments with an isolated aptamer that show that this helix is less stable without Mn(12). Analysis of single-nucleotide trajectories revealed that A11 reactivity transiently spikes at ∼205 nt, whether or not Mn is included in the transcription reaction, but the amplitude of the spike is Mn-dependent (Fig. 6D). The spike in A11 reactivity coincides with P1.1 strand invasion and its reduced amplitude +Mn indicates A11 remains trapped in the long-range A-minor interaction in Mn-bound aptamer. The L1/P1.1 reactivity difference heatmap ±Mn highlights A11 as the nucleotide most sensitive to the presence of the ligand, serving as a marker for the riboswitching point (Fig. 6D). We also assessed reactivity changes for A48 in L3 during expression platform folding, since A48 stacks with A11 and helps anchor the cross-helix A-minor interaction. A48 reactivity drops ±Mn starting at transcript length ∼185 nt, with a larger decrease –Mn (Fig. 6E). A48 reactivity is consistently lower –Mn for the remainder of expression platform folding (Fig. 6E), likely due to broken A48-A11 stacking.

We observed similar reactivity trends in *mntP* expression platform folding at pH 7.2 as pH 8.5 (e.g., A11 reactivity spike, SI Fig. 12B-C) and uncovered additional evidence of P1.1/L1 disruption in the vicinity of the riboswitching point (see Supplementary Note 6 and SI Fig. 12D). Taken together, our observations highlight that a global structural change occurs in the *mntP* expression platform at transcript lengths ∼200-205 nt, which is likely prompted by the start of strand invasion at P1.1 in the aptamer (see Supplementary Note 6 and SI Fig. 12D for additional analysis of pH- and Mn-dependent reactivity changes in *mntP* L1, L3, and P1.1 during expression platform folding).

#### Co-transcriptional binding of Mn is required for riboswitching

We next evaluated the differences between co-transcriptionally folded and equilibrium refolded *mntP* intermediate transcripts that contain both the aptamer domain and expression platform, to assess whether co-transcriptional folding is required to observe Mn-coupled structural rearrangements in the expression platform. While we observed similar P5 folding transitions (SI Fig. 13A-B), the Mn-dependent spike in A11 reactivity is abolished when the RNA is equilibrium refolded ±Mn at either pH 7.2 or 8.5 (SI Fig. 13C). These data support that co-transcriptional Mn-binding is critical for stabilization of the A-minor motif that docks the two legs of the aptamer together and for prevention of the P1.1/L1 invader strand from disrupting the aptamer (see Supplementary Note 7 and SI Fig. 13D for reactivity analysis of the A11 stacking partner, A48, in equilibrium refolded intermediates). Further supporting the above observations, we found that T7-synthesized, equilibrium refolded *mntP* riboswitch adopts a translationally inactive conformation ±Mn (DMS/1M7-probed, described further in Supplementary Note 8 and SI Fig. 14).

### Analysis of *alx* expression platform folding during transcription

#### Distinct features of the alx expression platform folding

The expression platform for the *alx* riboswitch contains multiple helices that begin folding at transcript length ∼155 nt, as detected by coordinated drops in reactivity for the constituent nucleotides in each helix (Fig. 7A and SI Fig. 15A). The *alx* expression platform contains two helices (termed here P5 and P6) in its translationally inactive conformation that are predicted to fold –Mn, resulting in the ribosome binding sequence (RBS) base-paring with a disrupted P1.1/L1 (Fig. 7B-D). On the other hand, when P1.1/L1 remain base-paired in the Mn-bound *alx* aptamer, an alternative helix in the expression platform (termed here P5/P6) is predicted to fold, and the downstream RBS-containing strand is single-stranded (Fig. 7E-G).

Unlike the *mntP* riboswitch, the *alx* P1.1/L1 rising stem is highly dynamic during the entirety of *alx* expression platform folding both ±Mn, which was detected as consistent reactivity towards DMS for multiple nt (Fig. 7A). The strand predicted to invade P1.1/L1 in the *alx* aptamer emerges at transcript length 155 nt, resulting in formation of the P5 helix (Fig. 7A). Once the expression platform is extended to transcript length 190 nt, the P6 intrinsic terminator helix folds (Fig. 7A). Interestingly, the DMS reactivities for nucleotides in the *alx* expression platform were largely similar ±Mn, resulting in similar secondary structure predictions for expression platform folding intermediates (SI Fig. 15B-D).

Despite the global structural similarities in the *alx* expression platform secondary structural models ±Mn, we detected multiple Mn-dependent reactivity differences for specific nucleotides that indicate a modest ligand-dependent shift in the conformational ensemble. P5 stem-loop nucleotides are more reactive –Mn (Fig. 7C), consistent with a population bias towards the inactive conformer. A137 (bulged in active conformer vs. base-paired in inactive conformer) demonstrates markedly high reactivity in general and exhibits higher reactivity +Mn (Fig. 7F), indicating a shift in the conformational ensemble towards the *alx* active conformer.

#### pH effects during folding of the alx expression platform

We observed similar DMS reactivity patterns in the *alx* expression platform at neutral vs. alkaline pH, such as folding of P5 and P6 (SI Fig. 16A-B), indicating that a shift to alkaline pH impacts metal-sensing rather than directly affecting folding of the expression platform and thus riboswitching. However, we observed interesting pH-dependent reactivity trends for A137, which was more reactive +Mn at both pH and most reactive overall at pH 8.5 +Mn (SI Fig. 16B), supporting that the active conformer is sampled more frequently under this condition.

We next assessed pH- and Mn-dependent reactivity trends in the *alx* aptamer during expression platform folding (Fig. 8A-C). Contrasting *mntP*, A11 remains accessible for DMS methylation across all conditions tested with no significant pH and Mn-dependent differences (Fig 8B and SI Fig. 16C). Since A11 is conserved in *alx* and *mntP* L1, it is curious that A11 exhibits distinctive reactivity trends between the two riboswitches. Unlike in *mntP*, the absolutely conserved A37 in *alx* aptamer remains least reactive at pH 8.5 +Mn for the duration of expression platform folding (supportive of more stable Mn binding at alkaline pH), with multiple transient spikes in reactivity indicating a change in L3 dynamics at pH 8.5 +Mn during expression platform folding (Fig. 8C and SI Fig. 16C). Additionally, we observed interesting pH- and Mn-dependent reactivity trends for *alx* expression platform nt in equilibrium refolded intermediates (see Supplementary Note 9 and SI Fig. 17).

### The *mntP* and *alx* RBS regions are dynamic in co-transcriptionally displayed RNA

The RNA surrounding the *mntP* RBS did not demonstrate a clear Mn-dependent reactivity trend, with DMS-constrained structure models predicting low-T_m_ hairpins interspersed with single-stranded regions (Fig. 18A). We thus conclude that the RNA around the RBS is dynamic, and any Mn-dependent structural differences are likely averaged out. Prior smFRET study observed such RBS dynamics for the preQ_1_ translational riboswitch with ligand-dependent accessibility bursts in the RBS region(47). Therefore, characterization of the *mntP* RBS dynamics requires complementary, e.g., single-molecule methods. Likewise, heatmap visualization of the *mntP* RBS nucleotides at pH 7.2 or 8.5 ±Mn or in equilibrium refolded intermediates at both pH showed no clear reactivity difference that would indicate increased RBS accessibility +Mn (SI Fig. 18B-C).

For *alx*, the presence of the P6 intrinsic terminator stem resulted in a stark decrease in representation for longer transcripts containing the RBS (SI Fig. 19A), precluding accurate modeling of the RNA structure in this region. To boost representation of full-length *alx*, we performed a modified co-transcriptional chemical probing experiment with a terminally biotinylated DNA template and the fast-acting SHAPE reagent benzoyl cyanide (BzCN, reaction time ∼1s) to better capture the short-lived co-transcriptional, Mn-dependent structural changes in the RBS. Consistent with *mntP*, the single-length BzCN probing experiment revealed no clear Mn-dependent differences in the reactivities of RBS nucleotides (Fig. 19B), supporting that the *alx* RBS is dynamic. These dynamics may serve a functional role; for instance, the RBS may be rapidly docking/undocking with another tertiary structural element in *alx*, with the docking rate fine-tuning RBS accessibility. Underscoring the importance of co-transcriptional chemical probing, we found that T7-synthesized, equilibrium refolded *alx* riboswitch adopts a translationally inactive conformation ±Mn (DMS/1M7-probed, described further in Supplementary Note 10 and SI Fig. 20). Taken together, while *alx* and *mntP* differ structurally, they may share common dynamic features that gate RBS accessibility.

## DISCUSSION

In this work, we performed a suite of chemical probing experiments on the Mn-sensing translational riboswitches in *E. coli* that precede the *mntP* and *alx* genes encoding Mn exporters. While static structural snapshots are available for Mn-sensing aptamers, these structures do not answer the question of how binding of Mn to the aptamer is “communicated” to the expression platform – the question at the heart of riboswitch function. Because the *mntP* and *alx* aptamers display unique architectures, they are an interesting test case of how structurally distinct aptamers in the same riboswitch family “talk” to the expression platform. We demonstrated that co-transcriptional folding with the native *E. coli* RNAP is essential for Mn-coupled structural rearrangements in the *mntP* and *alx* riboswitches *in vivo*, which prompted us to perform co-transcriptional chemical probing experiments.

The co-transcriptional chemical probing revealed that 5’-proximal halves in both *alx* and *mntP* aptamers traverse low-T_m_ RNA hairpin intermediates during folding, presumably as a safeguard against premature folding of RNA into stable but non-functional folds. Surprisingly, we found that both riboswitches show Mn-dependent reactivity changes *before* the full aptamer domain has folded. The Mn-dependent structural changes in aptamer folding intermediates map to nucleotides in the *alx* and *mntP* metal-sensing cores, indicating that a partially formed metal-sensing core can interrogate the environment for Mn. Curiously, these Mn-dependent differences emerge at much earlier transcript lengths for *alx* aptamer folding, which may permit a rapid response to the subtle increase in [Mn] prompted by cytosol alkalinization(19).

Notably, other riboswitches that have been investigated with co-transcriptional SHAPE-seq or SHAPE-MaP (ZTP, ppGpp, and fluoride) did not show ligand-dependent structural changes for folding intermediates(27). Ligands such as ZTP and ppGpp are larger than an Mn ion and require a network of interactions to bind their cognate aptamers(48, 49). The resulting structural complexity of these aptamer-ligand interactions likely prevents their stable interaction with an aptamer folding intermediate. In the case of fluoride riboswitch, binding of fluoride anion to the aptamer is mediated by a network of Mg^2+^ ions to overcome steric repulsion by the phosphate backbone(50), making fluoride less likely to bind an aptamer folding intermediate. Additionally, it will be interesting to see whether other divalent-sensing riboswitches (e.g., Mg^2+^- and Ni^2+^/Co^2+^-sensing riboswitches) can begin sampling for ligand before the complete aptamer has folded.

To get at the molecular basis of the divergent pH response of the *alx* and *mntP* riboswitches, we performed co-transcriptional DMS probing of RNA at both neutral and alkaline pH ±Mn. We found that pH uniquely affects the structure of the *alx* and *mntP* metal-sensing cores, providing the first clues to how pH differentially affects Mn sensing. The *alx* and *mntP* aptamers differ in their global architecture, forming a 3WJ and a 4WJ, respectively. Future study should investigate how nucleobase protonation states in the *alx* and *mntP* metal-sensing cores are coupled to junction dynamics, which controls the docking of the aptamer’s helical legs. Prior smFRET study on the *X. oryzae* aptamer, a 4WJ aptamer like *mntP*, illustrated how Mn binding promotes docking of the two helical legs that in turn stabilizes the P1.1 switch helix(12). The dynamics of *alx*-like aptamers has not yet been studied but may play a role in the differential pH effect on *alx* vs. *mntP* Mn sensing and consequent riboswitching.

The stark differences in the flexibility, architecture, and pH/Mn response of the *mntP* vs *alx* riboswitches may be coupled to the cellular function of their respective Mn exporters. We showed that alkaline pH alone does not rearrange the *alx* riboswitch into a translationally active conformer; rather, the *alx* riboswitch senses the increased cytosolic [Mn] brought about by cytosol alkalinization. We hypothesize that alkaline pH lowers the K_D_ of the *alx* aptamer for Mn, explaining why an alkaline pH shift and consequent increase of [Mn] from 28 to 42 µM (free and bound) activates expression of *alx* but not *mntP*(19) (Fig. 9). The physiological functions of such pH-enhanced expression of Mn exporters could be multi-fold. For example, Mn(II) is oxidized at alkaline pH(51, 52), deeming it unusable by Mn-dependent enzymes. Further, oxidized Mn(IV) is an extremely reactive agent able to oxidize carbon-carbon double bonds in nucleobases, the capability exploited in permanganate footprinting of protein-nucleic acid complexes(53, 54). Removal of Mn by exporters to minimize the pool of oxidized Mn could thus be a protective measure for the cell experiencing alkaline stress.

### Limitations of this study

There are multiple limitations to the experimental approaches used here. First, co-transcriptional chemical probing experiments are an endpoint measurement and do not capture the true structural intermediates traversed by an RNA during transcription. Further, the DMS probing reaction is 5 min., meaning that the co-transcriptionally displayed RNA can equilibrate to some degree. As a control, we performed DMS probing experiments on equilibrium refolded intermediates, which showed structural differences. However, we cannot rule out the possibility that the RNA folds observed with co-transcriptional DMS probing are not always representative of a true folding intermediate. Lastly, since the DMS probing reactions were performed *in vitro*, the absence of ribosomal proteins may have altered the observed structural ensemble, which should be addressed in future work.

## DATA AVAILABILITY

The raw sequencing data generated in this study will be publicly available in the Sequencing Read Archive (https://www.ncbi.nlm.nih.gov/sra) with the BioProject accession code PRJNA1197522. Information for individual BioSamples is provided in Supplementary Tables 7-8. All processed reactivity data will be deposited in the RNA Mapping Database (https://rmdb.stanford.edu/) and the individual accession codes will be provided in a Supplementary Table.

## SUPPLEMENTARY DATA

Supplementary Data are available at NAR online.

## AUTHOR CONTRIBUTIONS

C.S. and T.V.M, conceptualization; C.S. and D.P., methodology; C.S., D.P., and C.B., investigation; C.S. and D.P., formal analysis; C.S. and D.P., visualization; C.S. and D.P., writing-initial draft; C.S., D.P., and T.V.M, writing-review and editing; T.V.M, supervision; T.V.M., funding acquisition.

## Supporting information

SI_Tabes_1-8

## ACKNOWLEDGEMENTS

We thank Dr. Eric Strobel and Dr. Courtney Szyjka for protocols and helpful discussions regarding TECprobe-VL experiments. We also thank members of our research group for their valuable feedback during the course of this study.

