## Supporting_Information for "Structurally distinct manganese-sensing riboswitch aptamers regulate different expression platform architectures"

Christine N. Stephen[1], Danea E. Palmer[1], Clarisa Bautista, and Tatiana V. Mishanina[1*]
[1]Department of Chemistry and Biochemistry, University of California San Diego, 9500 Gilman Dr, La Jolla, CA 92093

This pdf file includes: Supplementary Figures 1-32, Supplementary Notes 1-10, and Supplemental Methods.

### Supplementary Figures

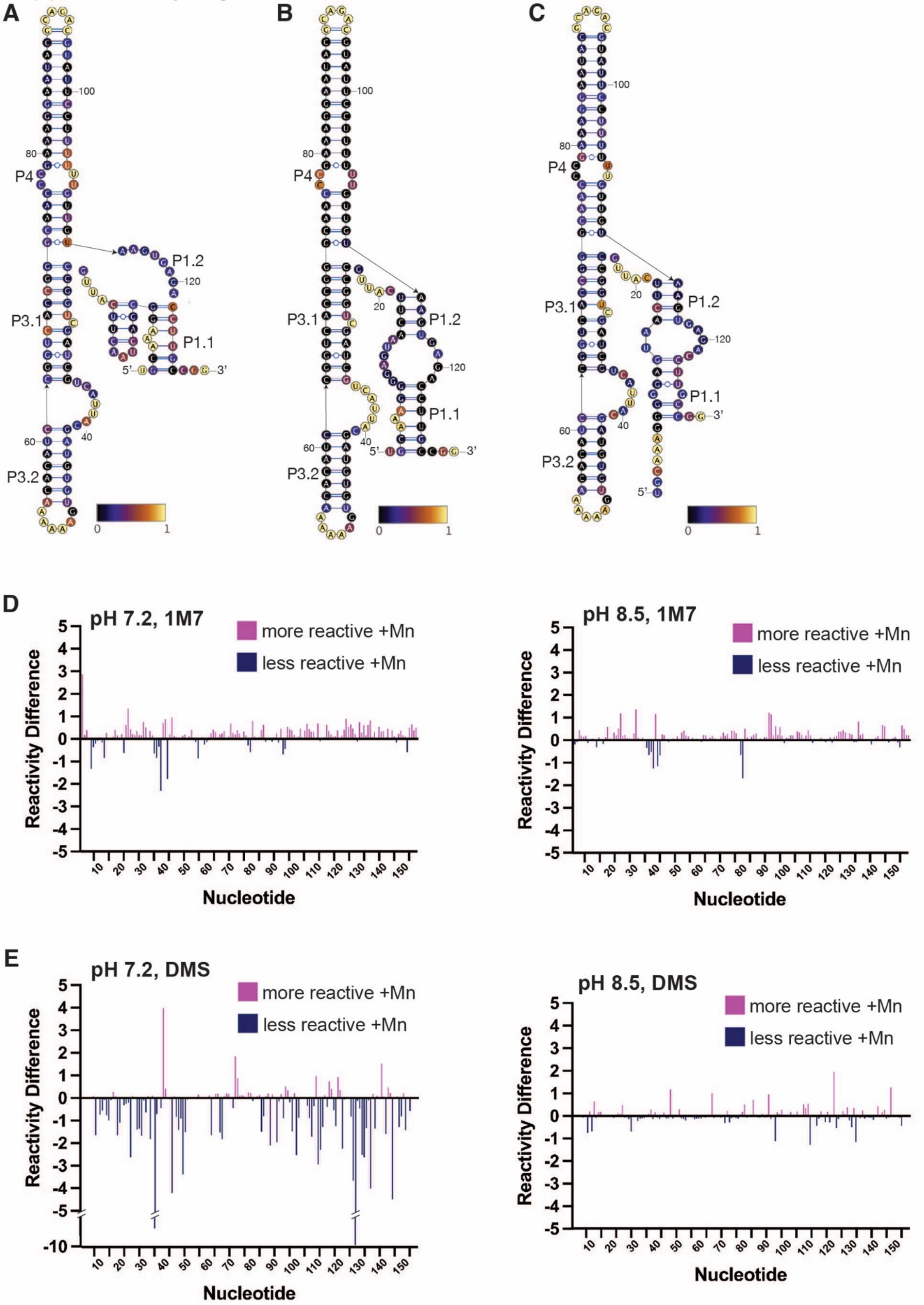

**Supplementary Figure 1. Structure probing of equilibrium-refolded *alx* aptamer RNA.** Secondary structure of equilibrium-refolded *alx* aptamer RNA modeled based on reactivity with 1M7 at **A.** pH 7.2 +Mn. **B.** pH 8.5 -Mn. **C.** pH 8.5 +Mn. **D.** Individual nucleotide reactivity differences when modified with 1M7 to compare +Mn vs. -Mn, measured at both pH 7.2 and 8.5. **E.** Individual nucleotide reactivity differences when modified with DMS to compare +Mn vs. -Mn, measured at both pH 7.2 and 8.5.

**TECprobe-VL construct designs, *alx* and *mntP* riboswitches**

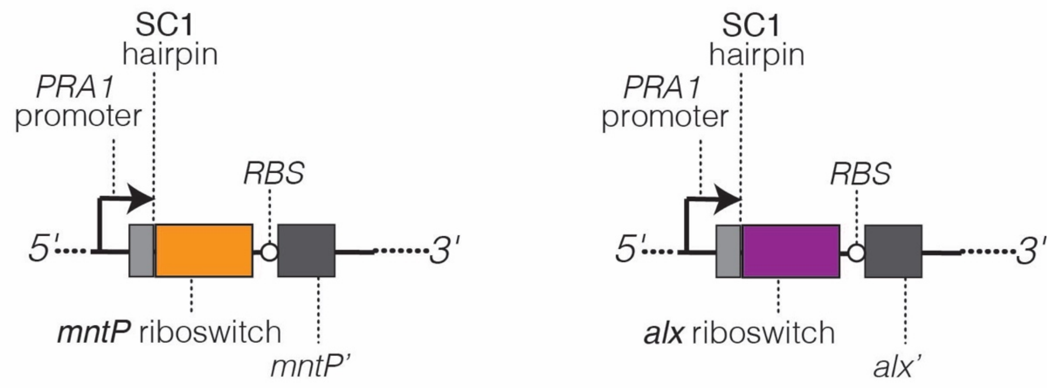


**Supplementary Figure 2. TECprobe-VL construct designs for the *alx* and *mntP* riboswitches.** Transcription template schematics annotated with key sequence elements. All TECprobe-VL transcription templates contained the PRA1 promoter, 5' SC1 hairpin, full-length *alx* or *mntP* riboswitch sequence, the RBS, and a portion of the downstream gene sequence (termed *alx'* - nt 1-100 *alx* coding sequence, and *mntP*''-nt 1-45 *mntP* coding sequence).

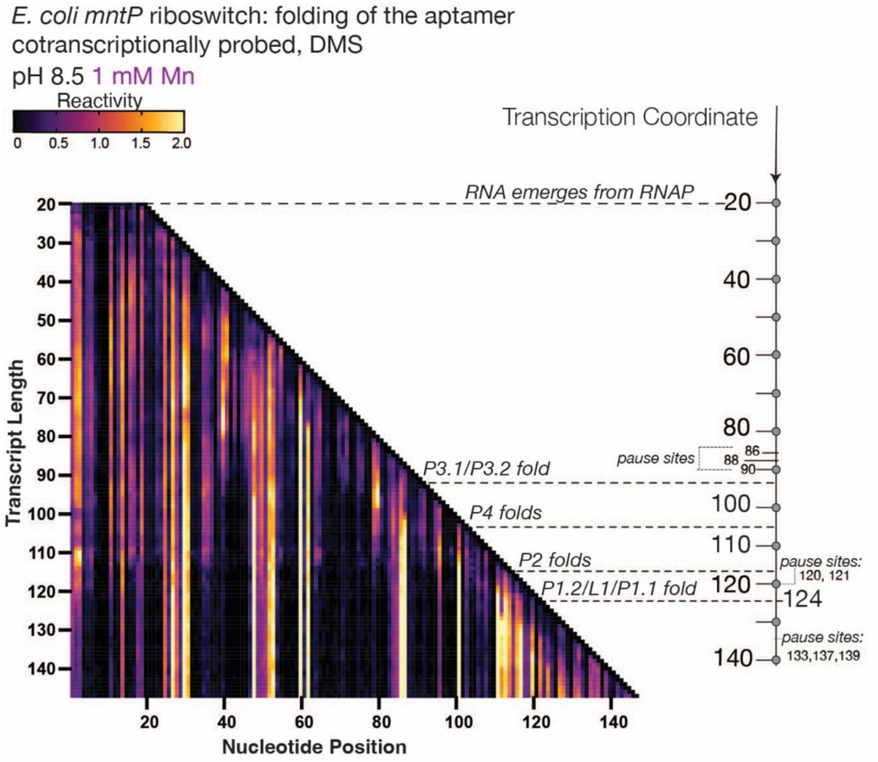


**Supplementary Figure 3. Co-transcriptional DMS probing of the *mntP* riboswitch aptamer at alkaline pH +Mn.** TECprobe-VL DMS reactivity matrix for the *mntP* aptamer at pH 8.5 +1 mM Mn. Reactivities shown were normalized with a single, whole-dataset calculated normalization factor. Data are from two independent replicates that were concatenated and analyzed together.

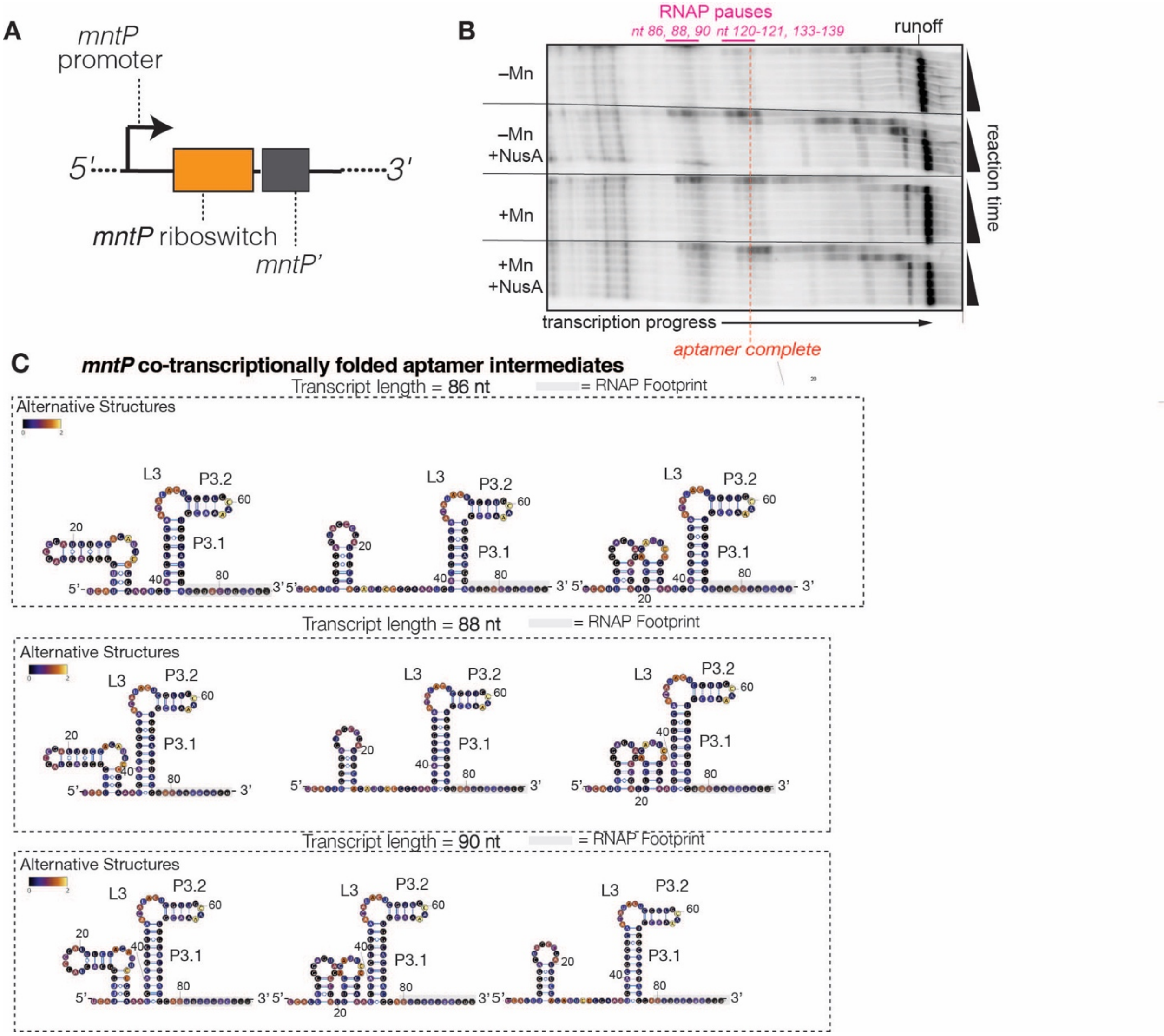


**Supplementary Figure 4. Synchronized, single-round *in vitro* transcription initiated from the *mntP* promoter and RNA structure models for pause-adjacent aptamer folding intermediates. A.** Transcription template schematic with key sequence features highlighted, including the native *mntP* promoter, full-length *mntP* riboswitch, and a portion of the downstream *mntP* gene (termed *mntP*'). **B.** *mntP* riboswitch transcripts were radiolabeled via incorporation of [$\alpha$-$^{32}$P]ATP and visualized on a 6% denaturing polyacrylamide gel. RNA species in the vicinity of pause sites are labeled with transcript lengths. Transcription elongation assays were performed ±Mn and ±NusA. **C.** Secondary structures of *mntP* aptamer folding intermediates, which were colored by the DMS reactivity of the indicated transcript at pH 8.5 -Mn.

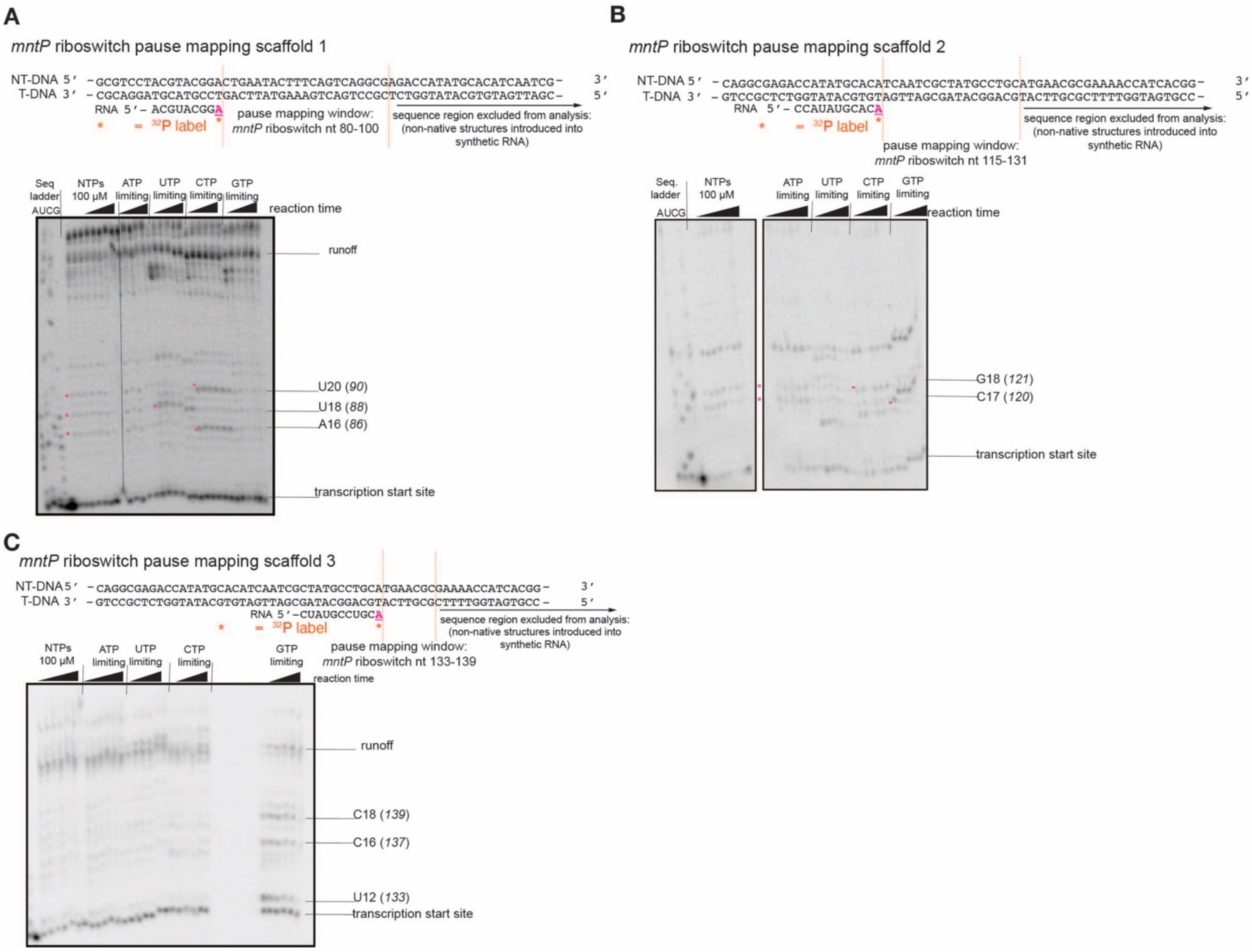


**Supplementary Figure 5. Mapping *mntP* pause locations during aptamer and early expression platform folding. A-C.** Experiments performed on synthetic nucleic acid scaffolds to map pause sites spanning transcript lengths 80-139 nt encoded by the *mntP* riboswitch sequence. Elongation complexes were positioned upstream of the designated pause mapping window and radiolabeled by incorporation of [$\alpha$-$^{32}$P]ATP. Transcription was initiated by addition of all four NTPs with one NTP present at a limiting concentration to increase pause duration. Radiolabeled RNAs were visualized on a 15% denaturing polyacrylamide gel. All pause sites are labeled next to gel images by the 3' RNA nucleotide in the RNAP active site and the length of radiolabeled RNA, with the corresponding transcript length in full-length *mntP* riboswitch denoted in parentheses. Pause sites were located based on the sequencing ladder and by comparing results from limiting NTP time courses.

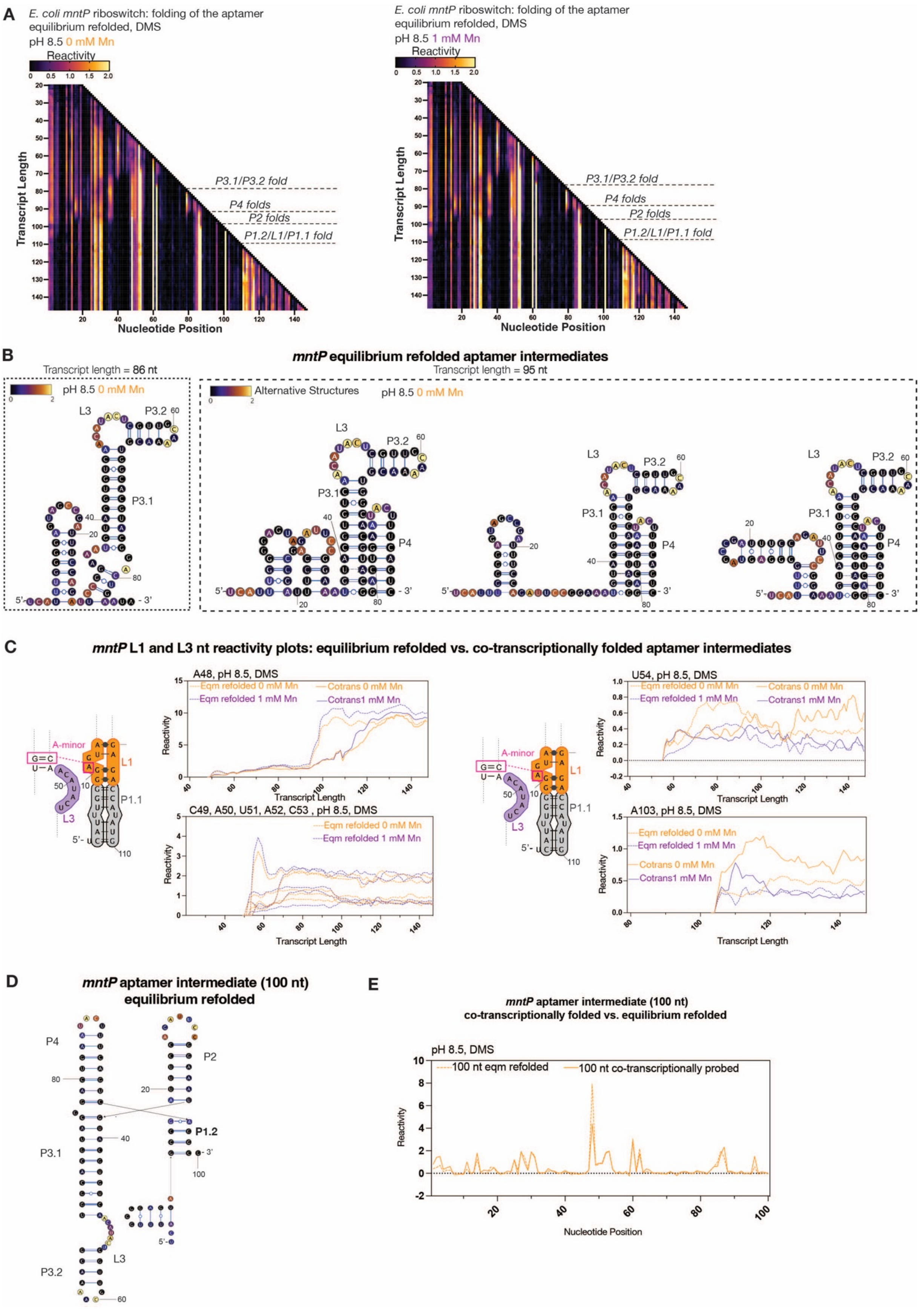
A
E. coli mntP riboswitch: folding of the aptamer equilibrium refolded, DMS
pH 8.5 0 mM Mn
pH 8.5 1 mM Mn
Reactivity
Transcript Length
Nucleotide Position
P3.1/P3.2 fold
P4 folds
P2 folds
P1.2/L1/P1.1 fold
B
mntP equilibrium refolded aptamer intermediates
Transcript length = 86 nt
Transcript length = 95 nt
Alternative Structures
C
mntP L1 and L3 nt reactivity plots: equilibrium refolded vs. co-transcriptionally folded aptamer intermediates
A48, pH 8.5, DMS
U54, pH 8.5, DMS
C49, A50, U51, A52, C53 , pH 8.5, DMS
A103, pH 8.5, DMS
Eqm refolded 0 mM Mn
Eqm refolded 1 mM Mn
Cotrans 0 mM Mn
Cotrans1 mM Mn
A-minor
D
mntP aptamer intermediate (100 nt) equilibrium refolded
E
mntP aptamer intermediate (100 nt) co-transcriptionally folded vs. equilibrium refolded
pH 8.5, DMS
100 nt eqm refolded
100 nt co-transcriptionally probed

**Supplementary Figure 6. Equilibrium DMS probing of *mntP* aptamer folding intermediates at alkaline pH ±Mn. A.** TECprobe-VL DMS reactivity matrices for *mntP* aptamer folding intermediates, which were purified and equilibrium refolded at pH 8.5 ± 1 mM Mn. Reactivities shown were normalized with a single, whole-dataset calculated normalization factor. Data are from two independent replicates that were concatenated and analyzed together. **B.** Secondary structures of *mntP* aptamer folding intermediates, which were purified and equilibrium refolded prior to DMS probing. Secondary structure models are colored by the DMS reactivity of the indicated transcript at pH 8.5 -Mn. **C.** Plots showing *mntP* L3 and L1 nt transcript length- and Mn- dependent reactivity changes in equilibrium refolded vs. co-transcriptionally folded aptamer intermediates. DMS data from 0 mM Mn (orange) and 1 mM Mn (purple) conditions from Fig. 3A, SI Fig. 3, and SI Fig. 6A. **D.** Secondary structure model for the 100 nt *mntP* aptamer folding intermediate, which was purified and equilibrium refolded prior to DMS probing. The secondary structure model is colored by the DMS reactivity at pH 8.5 -Mn. **E.** Plot of nt reactivities in the equilibrium refolded vs. co-transcriptionally folded 100 nt *mntP* aptamer intermediate, probed with DMS at pH 8.5 -Mn. The DMS reactivities for the co-transcriptionally folded RNA are from transcript length 115nt, which has 100 nt of structured RNA exposed for chemical probing due to the RNAP footprint. Data from SI Fig. 6A and Fig. 3A.

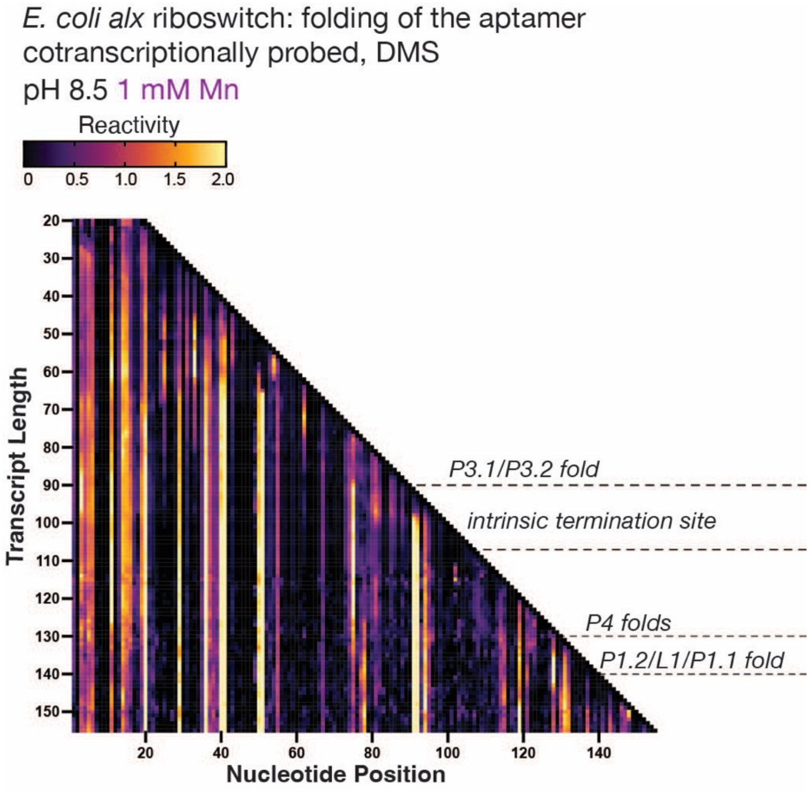


**Supplementary Figure 7. Co-transcriptional DMS probing of the *alx* riboswitch aptamer at alkaline pH +Mn.** TECprobe-VL reactivity matrix for the DMS-probed *alx* aptamer at pH 8.5 + 1 mM Mn. Reactivities shown were normalized with a single, whole-dataset calculated normalization factor. Data are from five independent replicates that were concatenated and analyzed together.

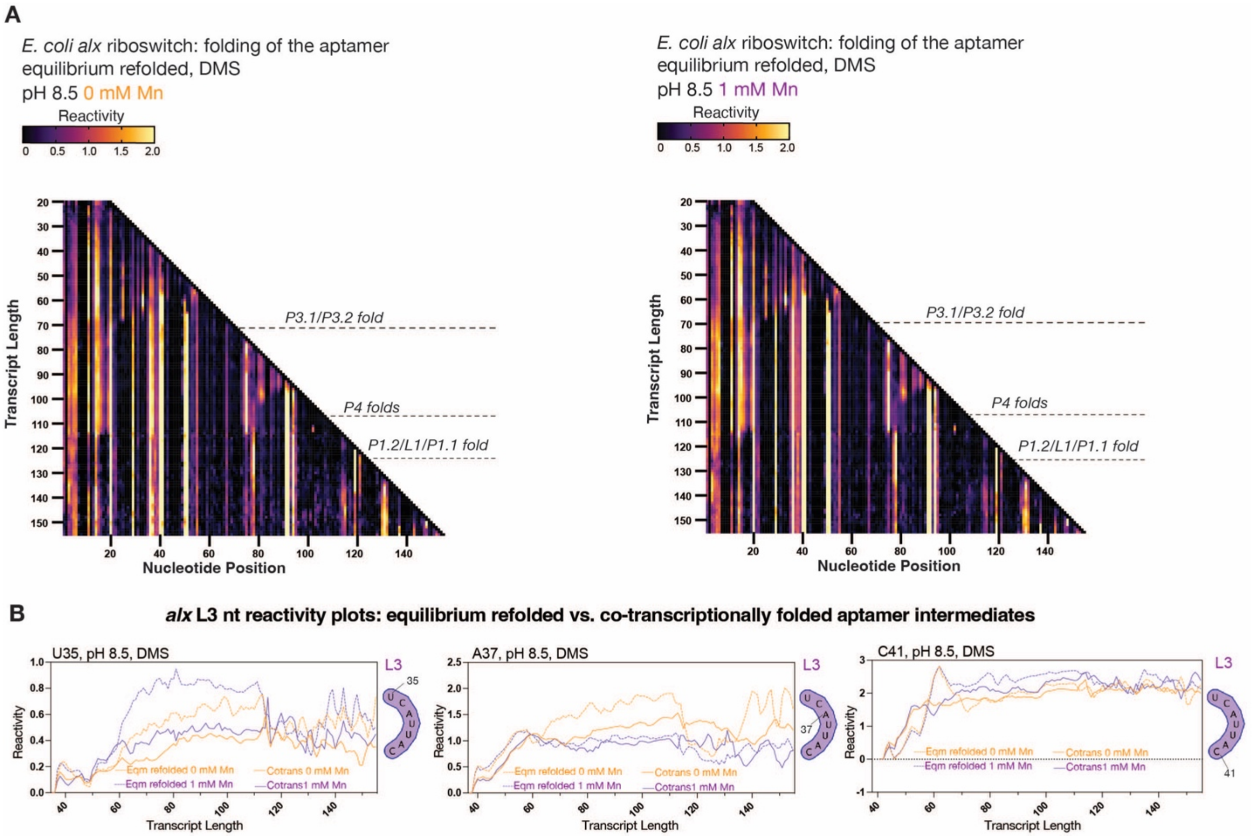


**Supplementary Figure 8. Equilibrium DMS probing of *alx* aptamer folding intermediates at alkaline pH ±Mn. A.** TECprobe-VL DMS reactivity matrices for *alx* aptamer folding intermediates, which were purified and equilibrium refolded prior to DMS probing at pH 8.5 ± 1 mM Mn. Reactivities shown were normalized with a single, whole-dataset calculated normalization factor. Data are from two independent replicates that were concatenated and analyzed together. **B.** Plots showing *alx* L3 nt transcript length- and Mn-dependent reactivity changes in equilibrium refolded vs. co-transcriptionally folded aptamer intermediates at pH 8.5 ± 1 mM Mn. DMS data from 0 mM Mn (orange) and 1 mM Mn (purple) conditions are from Fig. 4A, Fig. 7, and SI Fig. 8A.

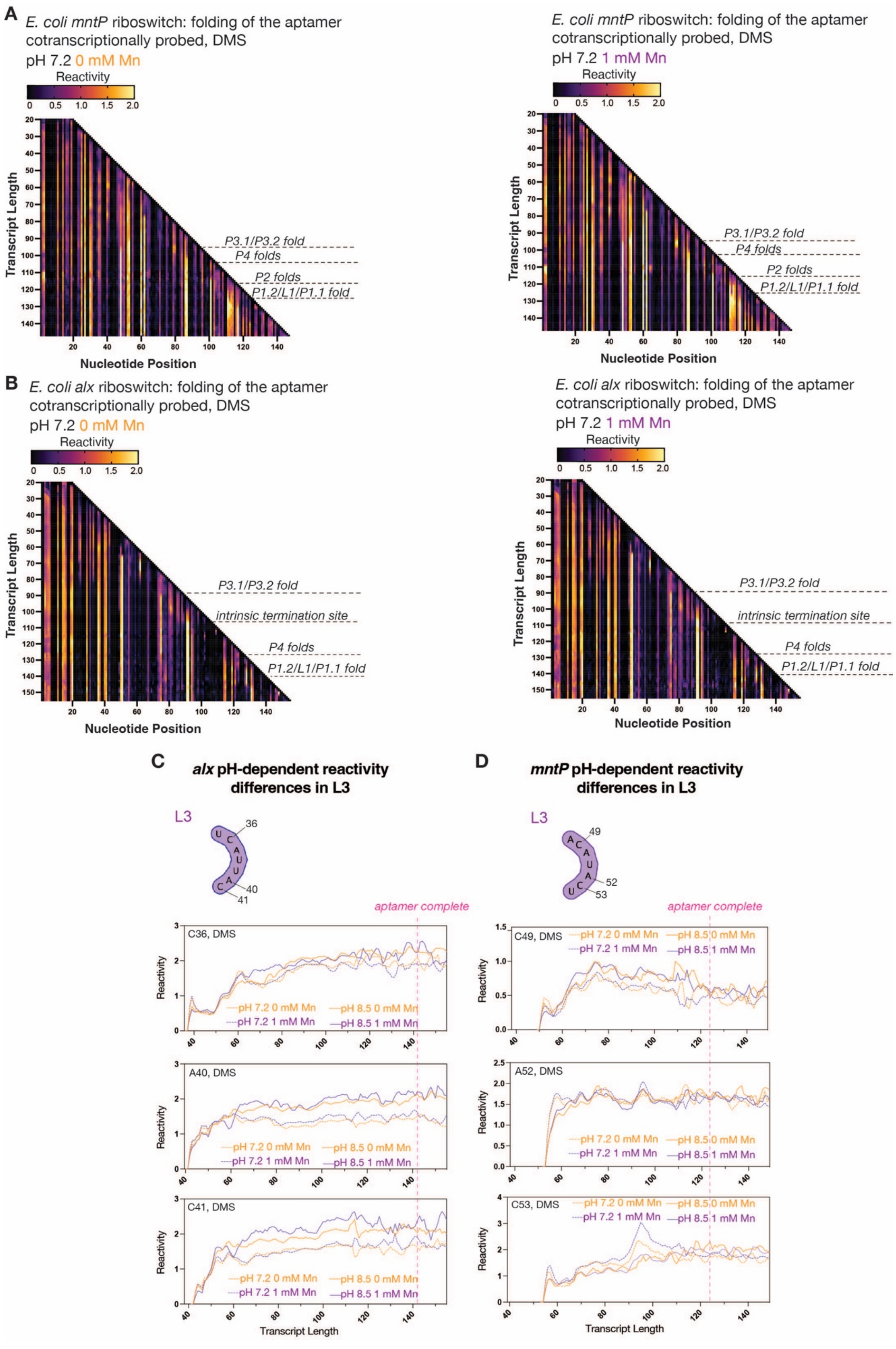

A
E. coli mntP riboswitch: folding of the aptamer cotranscriptionally probed, DMS
pH 7.2 0 mM Mn
Reactivity
P3.1/P3.2 fold
P4 folds
P2 folds
P1.2/L1/P1.1 fold
Transcript Length
Nucleotide Position
E. coli mntP riboswitch: folding of the aptamer cotranscriptionally probed, DMS
pH 7.2 1 mM Mn
B
E. coli alx riboswitch: folding of the aptamer cotranscriptionally probed, DMS
pH 7.2 0 mM Mn
intrinsic termination site
E. coli alx riboswitch: folding of the aptamer cotranscriptionally probed, DMS
pH 7.2 1 mM Mn
C
alx pH-dependent reactivity differences in L3
D
mntP pH-dependent reactivity differences in L3
L3
aptamer complete
C36, DMS
A40, DMS
C41, DMS
C49, DMS
A52, DMS
C53, DMS
pH 7.2 0 mM Mn
pH 7.2 1 mM Mn
pH 8.5 0 mM Mn
pH 8.5 1 mM Mn
Reactivity
Transcript Length

**Supplementary Figure 9. Co-transcriptional DMS probing of the *alx* and mntP aptamers at neutral pH ±Mn and pH-dependent reactivity changes in L3. A-B.** TECprobe-VL DMS reactivity matrices for the mntP (**A**) and alx (**B**) aptamers at pH 7.2 ± 1 mM Mn. Reactivities shown were normalized with a single, whole-dataset calculated normalization factor. Data are from two (*mntP*, **A**) or five (*alx*, **B**) independent replicates that were concatenated and analyzed together. **C-D.** Plots showing *alx* (**C**) and *mntP* (**D**) L3 nt transcript length-, pH-, and Mn-dependent reactivity changes in co-transcriptional aptamer folding intermediates at pH 7.2 or 8.5 ± 1 mM Mn. DMS data from 0 mM Mn (orange) and 1 mM Mn (purple) are from Fig. 4A, Fig. 7, and SI Fig. 9B for *alx* and Fig. 3A, SI Fig. 3, and SI Fig. 9A for *mntP*. Vertical dotted line marks completion of aptamer folding.

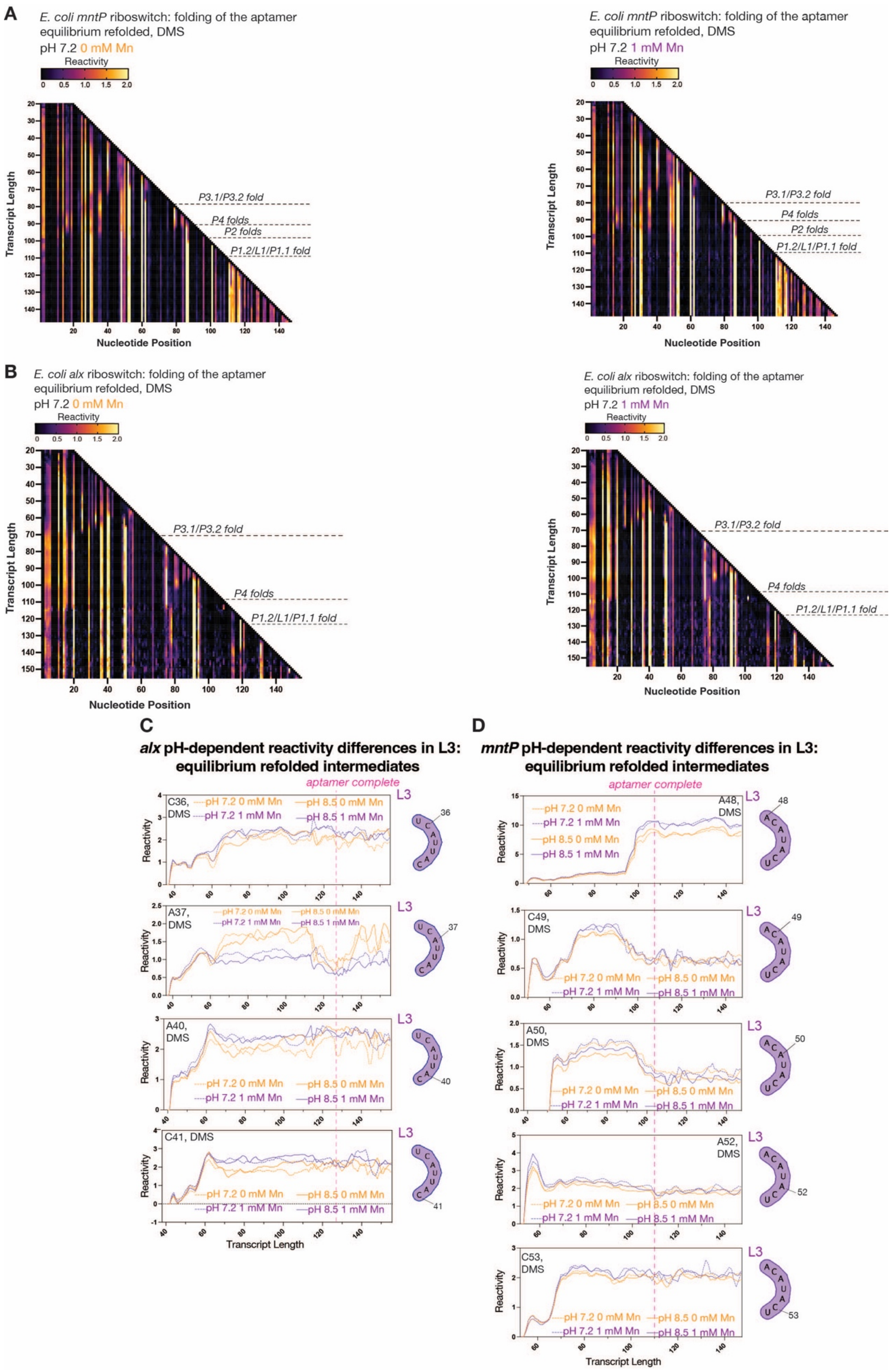
A
E. coli mntP riboswitch: folding of the aptamer
equilibrium refolded, DMS
pH 7.2 0 mM Mn
E. coli mntP riboswitch: folding of the aptamer
equilibrium refolded, DMS
pH 7.2 1 mM Mn
P3.1/P3.2 fold
P4 folds
P2 folds
P1.2/L1/P1.1 fold
Transcript Length
Nucleotide Position
B
E. coli alx riboswitch: folding of the aptamer
equilibrium refolded, DMS
pH 7.2 0 mM Mn
E. coli alx riboswitch: folding of the aptamer
equilibrium refolded, DMS
pH 7.2 1 mM Mn
C
alx pH-dependent reactivity differences in L3: equilibrium refolded intermediates
D
mntP pH-dependent reactivity differences in L3: equilibrium refolded intermediates
aptamer complete
pH 7.2 0 mM Mn
pH 7.2 1 mM Mn
pH 8.5 0 mM Mn
pH 8.5 1 mM Mn
Reactivity
Transcript Length

**Supplementary Figure 10. Equilibrium DMS probing of *alx* and *mntP* aptamer folding intermediates at neutral pH ±Mn and pH-dependent reactivity changes in L3. A-B.** TECprobe-VL DMS reactivity matrices for the *mntP* (**A**) and *alx* (**B**) aptamer intermediates, which were purified and equilibrium refolded at pH 7.2 ± 1 mM Mn. Reactivities shown were normalized with a single, whole-dataset calculated normalization factor. Data are from two independent replicates that were concatenated and analyzed together**. C-D.** Plots showing *alx* (**C**) and *mntP* (**D**) L3 transcript length-, pH-, and Mn-dependent reactivity changes in equilibrium refolded aptamer intermediates at pH 7.2 or 8.5 ± 1 mM Mn. DMS data from 0 mM Mn (orange) and 1 mM Mn (purple) are from SI Fig. 8A and SI Fig. 10B for *alx* and SI Fig. 6A and SI Fig. 10A for *mntP*. Vertical dotted line marks completion of aptamer folding.

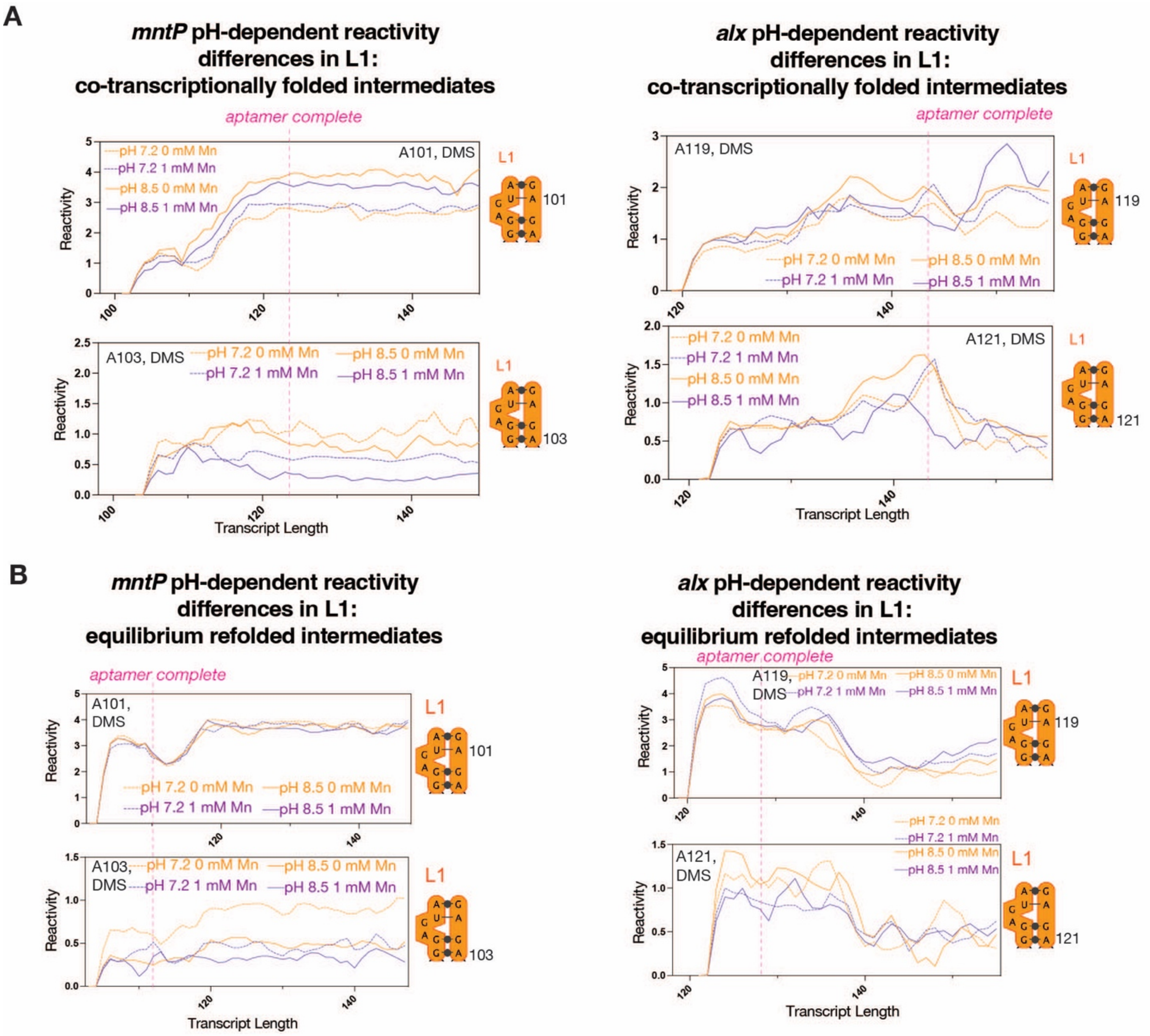


**Supplementary Figure 11. pH-dependent reactivity changes in *alx* and *mntP* L1 in co-transcriptionally folded and equilibrium refolded aptamer intermediates**

**A.** Plots showing *alx* and *mntP* L1 nt transcript length-, pH-, and Mn-dependent reactivity changes in co-transcriptionally folded aptamer intermediates at pH 7.2 or 8.5 ± 1 mM Mn. DMS data from 0 mM (orange) and 1 mM Mn (purple) conditions are from Fig. 4A, SI Fig. 7, and SI Fig. 9B for *alx* and Fig. 3A, SI Fig. 3, and SI Fig. 9A for *mntP*. Vertical dotted line marks completion of aptamer folding. **B.** Plots showing *alx* and *mntP* L1 nt transcript length-, pH-, and Mn-dependent reactivity changes in equilibrium refolded aptamer intermediates at pH 7.2 or 8.5 ± 1 mM Mn. DMS data from 0 mM (orange) and 1 mM Mn (purple) conditions from SI Fig. 8A and SI Fig. 10B for *alx* and SI Fig. 6A and SI Fig. 10A for *mntP*. Vertical dotted line marks completion of aptamer folding.

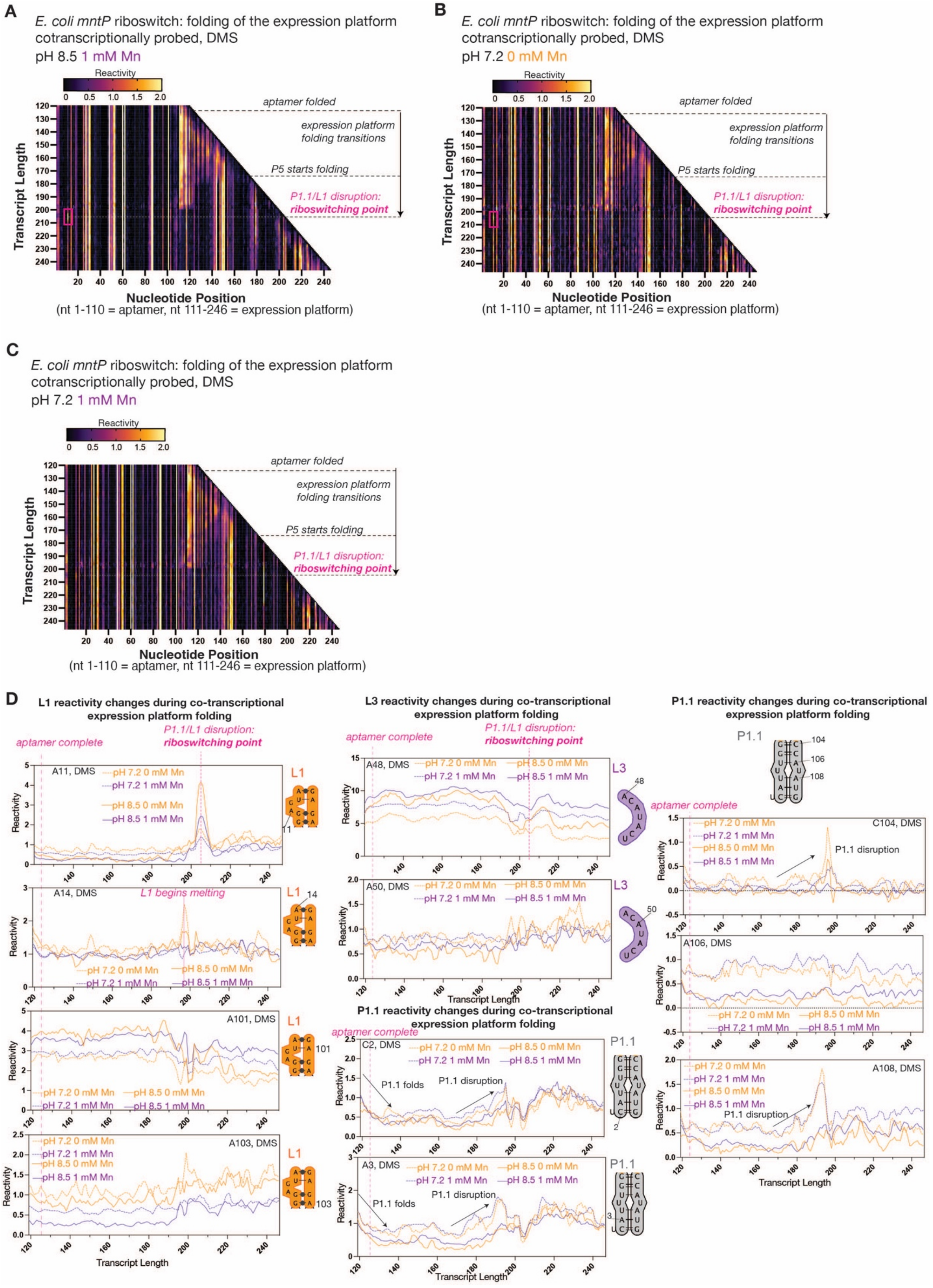

A
E. coli mntP riboswitch: folding of the expression platform cotranscriptionally probed, DMS
pH 8.5 1 mM Mn
Reactivity
0 0.5 1.0 1.5 2.0
aptamer folded
expression platform folding transitions
P5 starts folding
P1.1/L1 disruption: riboswitching point
Transcript Length
Nucleotide Position
(nt 1-110 = aptamer, nt 111-246 = expression platform)
B
E. coli mntP riboswitch: folding of the expression platform cotranscriptionally probed, DMS
pH 7.2 0 mM Mn
C
E. coli mntP riboswitch: folding of the expression platform cotranscriptionally probed, DMS
pH 7.2 1 mM Mn
D
L1 reactivity changes during co-transcriptional expression platform folding
aptamer complete
P1.1/L1 disruption: riboswitching point
A11, DMS
pH 7.2 0 mM Mn
pH 7.2 1 mM Mn
pH 8.5 0 mM Mn
pH 8.5 1 mM Mn
L1
A14, DMS
L1 begins melting
A101, DMS
A103, DMS
Reactivity
Transcript Length
L3 reactivity changes during co-transcriptional expression platform folding
A48, DMS
A50, DMS
L3
P1.1 reactivity changes during co-transcriptional expression platform folding
C2, DMS
A3, DMS
P1.1 folds
P1.1 disruption
P1.1
C104, DMS
A106, DMS
A108, DMS

**Supplementary Figure 12. Co-transcriptional DMS probing of *mntP* riboswitch during expression platform synthesis at neutral and alkaline pH. A-C.** TECprobe-VL DMS reactivity matrices for *mntP* folding intermediates at pH 8.5 +1 mM Mn (**A**), pH 7.2 (**B**), and pH 7.2 + 1 mM Mn (**C**). Reactivities shown were normalized with a single, whole-dataset calculated normalization factor. Data are from two independent replicates that were concatenated and analyzed together. **D.** Plots showing *mntP* L1, L3, and P1.1 nt transcript length-, pH-, and Mn-dependent reactivity changes in co-transcriptionally folded intermediates at pH 7.2 or 8.5 ± 1 mM Mn. DMS data from 0 mM Mn (orange) and 1 mM Mn (purple) conditions from Fig. 6A-B, and SI Fig. 12A-C. Vertical pink dotted lines mark key co-transcriptional RNA folding transitions.

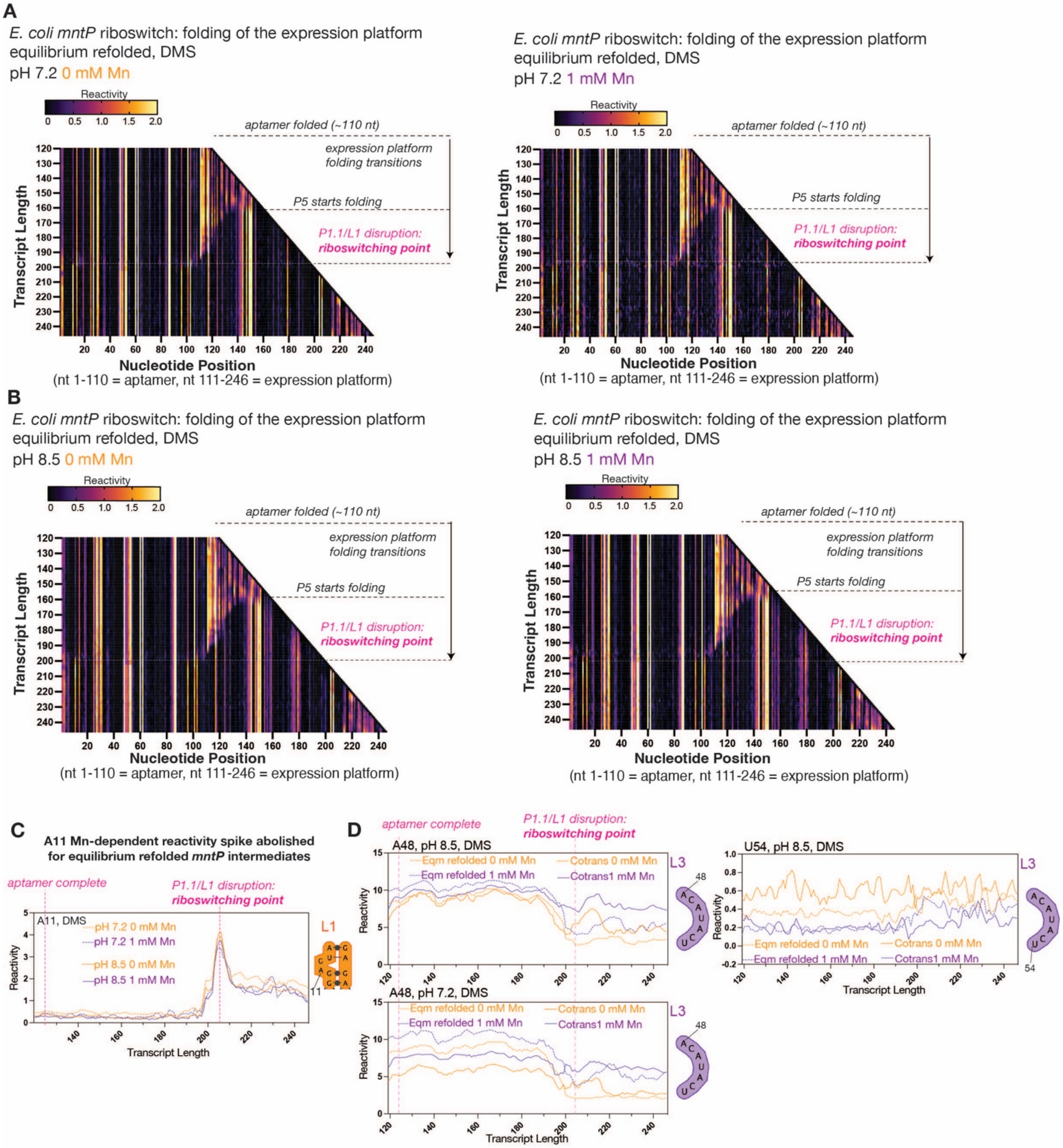


**Supplementary Figure 13. Equilibrium DMS probing on *mntP* folding intermediates during expression platform synthesis. A-C.** TECprobe-VL DMS reactivity matrices for *mntP* folding intermediates, which were purified and equilibrium refolded at pH 7.2 ± 1 mM Mn (**A**) or pH 8.5 ± 1 mM Mn (**B**). Reactivities shown were normalized with a single, whole-dataset calculated normalization factor. Data are from two independent replicates that were concatenated and analyzed together. **C.** Plot showing *mntP* A11 transcript length-, pH-, and Mn-dependent reactivity changes in equilibrium refolded intermediates at pH 7.2 or 8.5 ± 1 mM Mn. Data for 0 mM Mn (orange) and 1 mM Mn (purple) from SI Figs. 13A-B. Vertical dotted lines mark indicated RNA structural transitions. **D.** Plots showing *mntP* L3 nt transcript length- and Mn-dependent reactivity changes in co-transcriptionally folded vs. equilibrium refolded intermediates, which were DMS-probed at the designated pH value. DMS data for 0 mM Mn (orange) and 1 mM Mn (purple) conditions from Fig. 6A-B, SI Fig. 12A-C, and SI Figs. 13A-B. Vertical dotted lines mark indicated co-transcriptional RNA folding transitions.

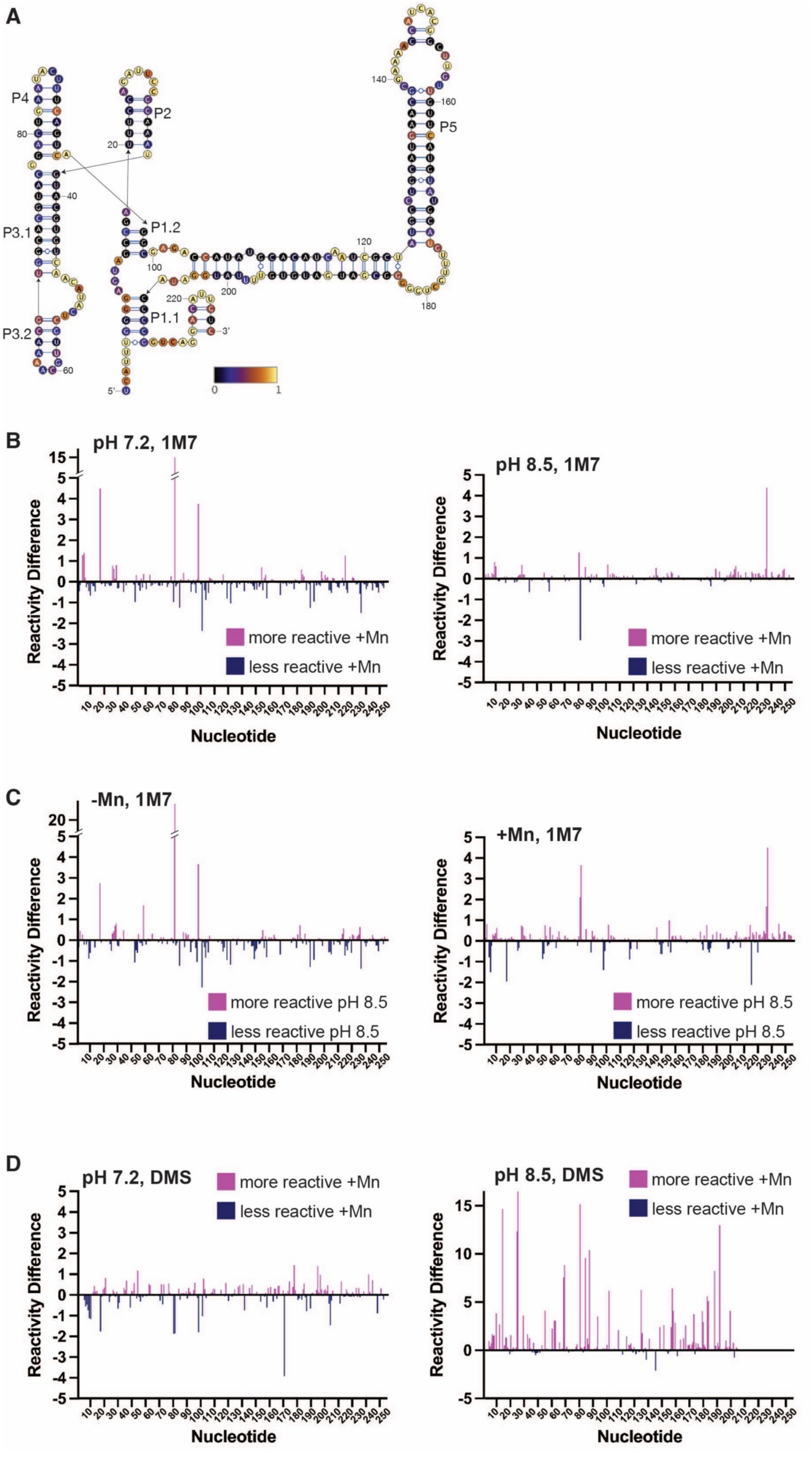
A
P4
P2
P5
P3.1
P1.2
P1.1
P3.2
5'
3'
0
1
B
pH 7.2, 1M7
pH 8.5, 1M7
Reactivity Difference
Nucleotide
more reactive +Mn
less reactive +Mn
C
-Mn, 1M7
+Mn, 1M7
more reactive pH 8.5
less reactive pH 8.5
D
pH 7.2, DMS
pH 8.5, DMS

**Supplementary Figure 14. Structure probing of equilibrium-refolded *mntP* full-length riboswitch RNA. A.** Secondary structure of equilibrium-refolded *mntP* full-length RNA modeled based on reactivity with 1M7 at pH 7.2 -Mn. **B.** Individual nucleotide reactivity differences when modified with 1M7 to compare +Mn vs. -Mn, measured at both pH 7.2 and 8.5. **C.** Individual nucleotide reactivity differences at neutral pH (7.2) vs. alkaline pH (8.5), measured both in the presence and absence of Mn. **D.** Individual nucleotide reactivity differences when modified with DMS to compare +Mn vs. -Mn, measured at both pH 7.2 and 8.5**.**

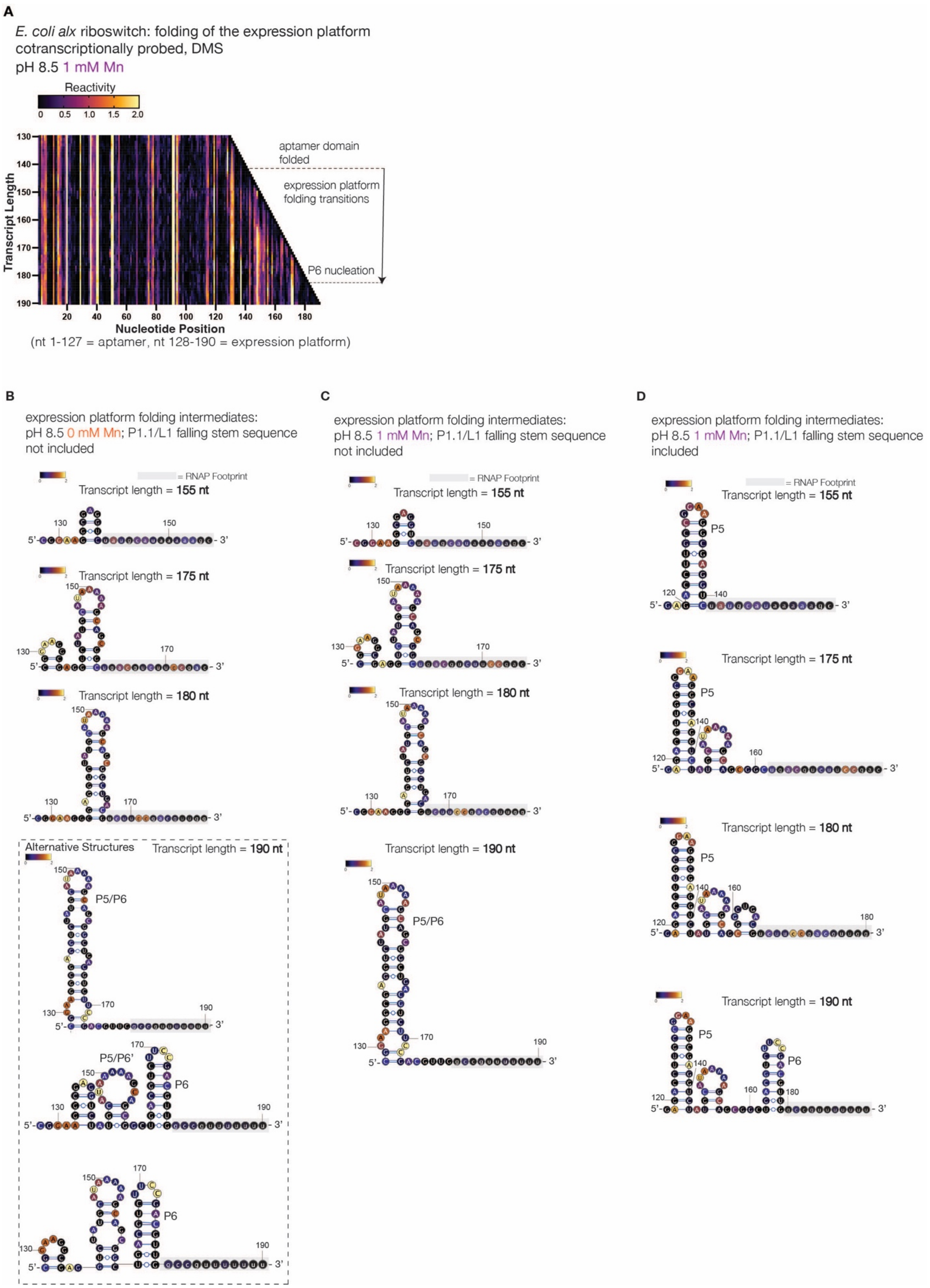
A
E. coli alx riboswitch: folding of the expression platform
cotranscriptionally probed, DMS
pH 8.5 1 mM Mn
Reactivity
0 0.5 1.0 1.5 2.0
Transcript Length
130
140
150
160
170
180
190
aptamer domain folded
expression platform folding transitions
P6 nucleation
20 40 60 80 100 120 140 160 180
Nucleotide Position
(nt 1-127 = aptamer, nt 128-190 = expression platform)
B
expression platform folding intermediates:
pH 8.5 0 mM Mn; P1.1/L1 falling stem sequence not included
= RNAP Footprint
Transcript length = 155 nt
Transcript length = 175 nt
Transcript length = 180 nt
Alternative Structures
Transcript length = 190 nt
P5/P6
P5/P6'
P6
C
expression platform folding intermediates:
pH 8.5 1 mM Mn; P1.1/L1 falling stem sequence not included
= RNAP Footprint
Transcript length = 155 nt
Transcript length = 175 nt
Transcript length = 180 nt
Transcript length = 190 nt
P5/P6
D
expression platform folding intermediates:
pH 8.5 1 mM Mn; P1.1/L1 falling stem sequence included
= RNAP Footprint
Transcript length = 155 nt
Transcript length = 175 nt
Transcript length = 180 nt
Transcript length = 190 nt
P5
P6
5'-
- 3'

**Supplementary Figure 15. Co-transcriptional DMS probing of the *alx* riboswitch during expression platform synthesis at alkaline pH. A.** TECprobe-VL DMS reactivity matrix for *alx* folding intermediates at pH 8.5 +1 mM Mn. Reactivities shown were normalized with a single, whole-dataset calculated normalization factor. Data are from five independent replicates that were concatenated and analyzed together. **B-D.** Secondary structure models for *alx* expression platform folding intermediates, which were co-transcriptionally folded prior to DMS probing. The secondary structure models are colored by DMS reactivity at pH 8.5 at the designated Mn concentration. To simplify the structure modeling, we executed the RNAstructure "Fold" command constrained by DMS reactivities for the expression platform sequence only, with the L1/P1.1 falling stem either included or excluded.

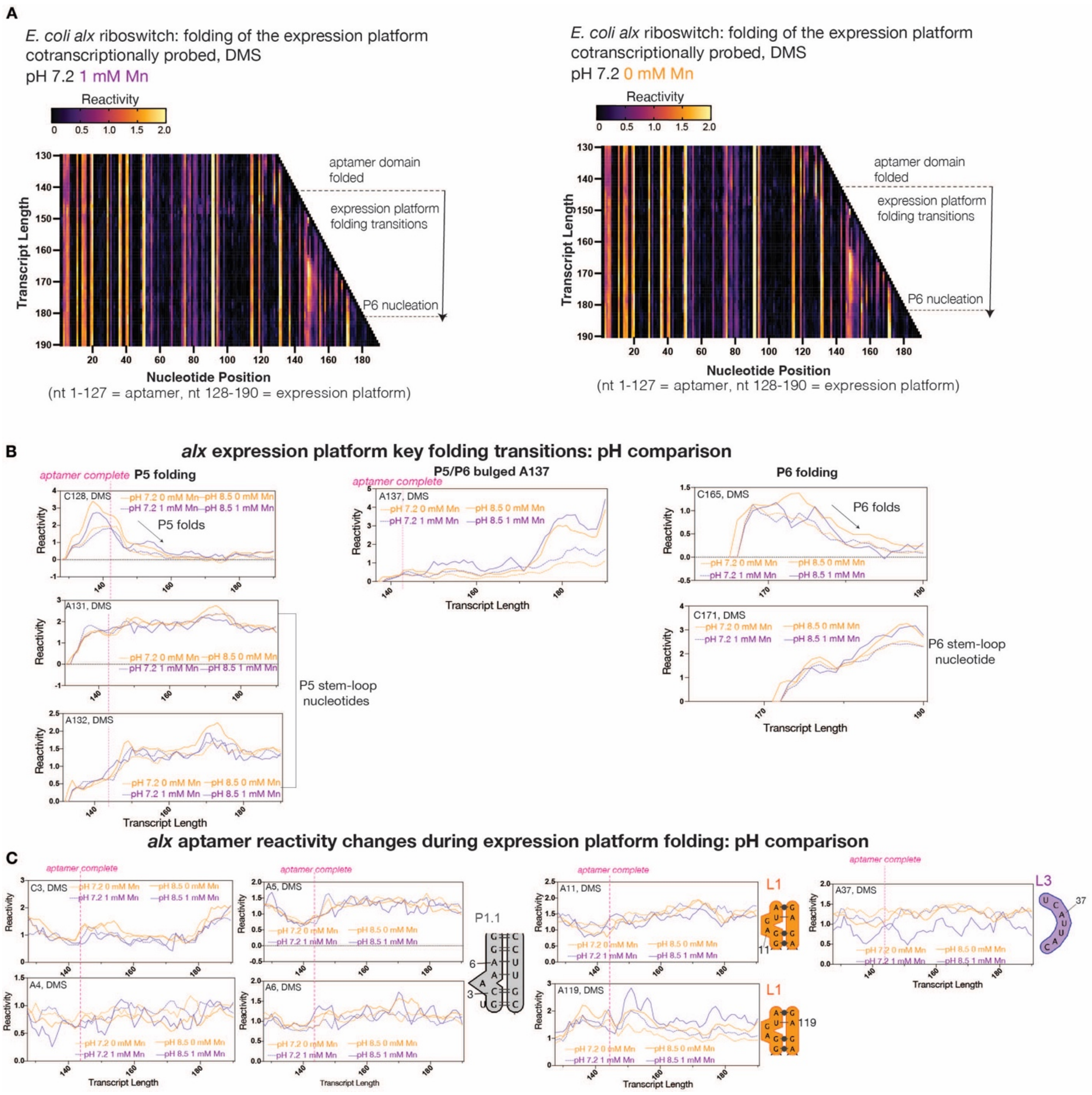

A
E. coli alx riboswitch: folding of the expression platform cotranscriptionally probed, DMS
pH 7.2 1 mM Mn
Reactivity
aptamer domain folded
expression platform folding transitions
P6 nucleation
Transcript Length
Nucleotide Position
(nt 1-127 = aptamer, nt 128-190 = expression platform)
E. coli alx riboswitch: folding of the expression platform cotranscriptionally probed, DMS
pH 7.2 0 mM Mn
B
alx expression platform key folding transitions: pH comparison
aptamer complete
P5 folding
P5/P6 bulged A137
P6 folding
P5 folds
P6 folds
P5 stem-loop nucleotides
P6 stem-loop nucleotide
C
alx aptamer reactivity changes during expression platform folding: pH comparison
P1.1
L1
L3

Mn (orange) and 1 mM Mn (purple) from Fig. 7A SI Fig. 15A, and SI Fig. 16A. Vertical dotted lines mark indicated RNA folding transitions. **C.** Plots showing *alx* aptamer nt transcript length-, pH-, and Mn-dependent reactivity changes in co-transcriptionally folded intermediates at pH 7.2 or 8.5 ± 1 mM Mn. DMS Data for 0 mM Mn (orange) and 1 mM Mn (purple) from Fig. 8A, SI Fig. 15A, and SI Fig. 16A. Vertical dotted lines mark indicated RNA folding transitions.

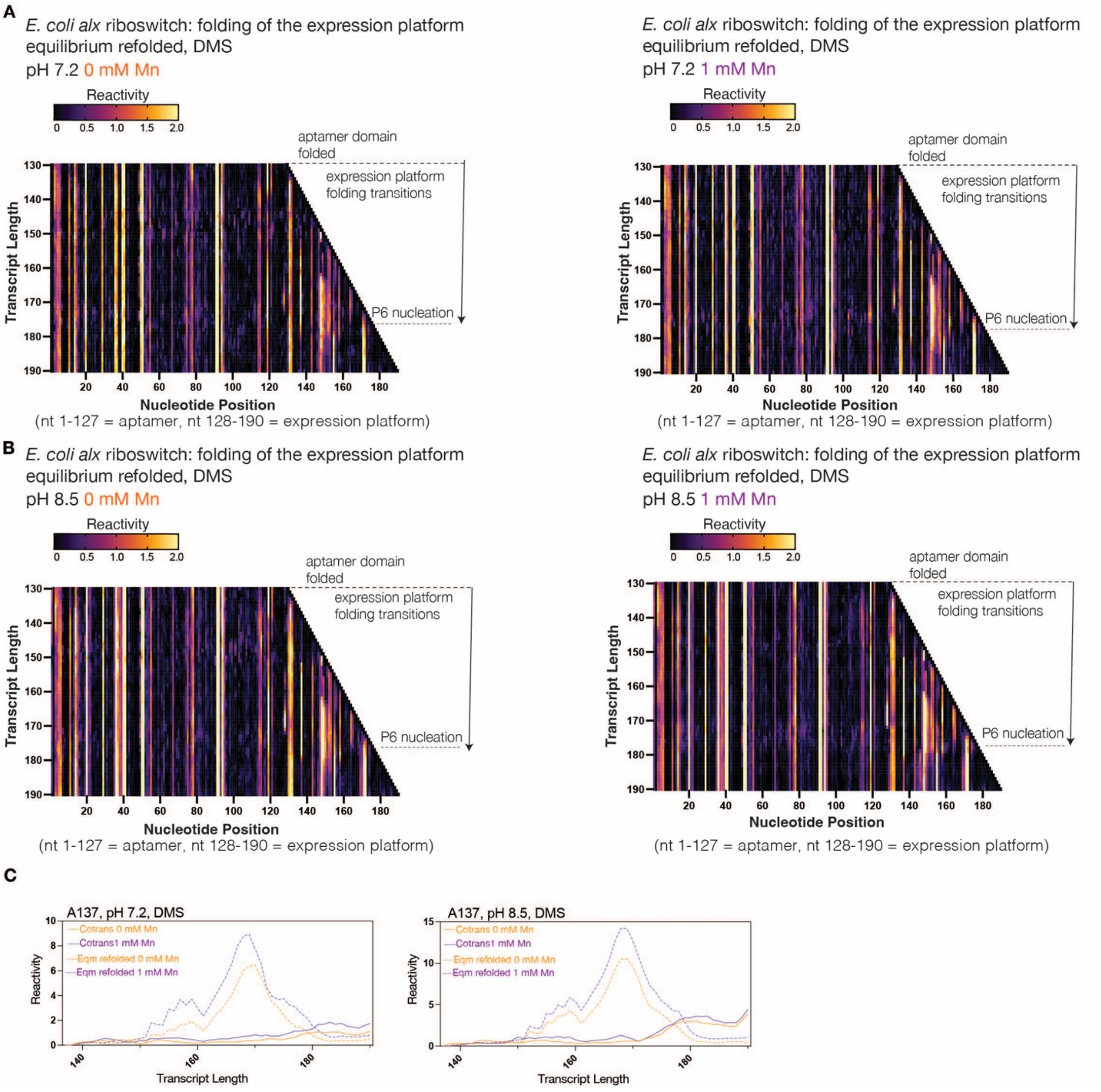


**Supplementary Figure 17. Equilibrium DMS probing of the *alx* riboswitch during expression platform synthesis at neutral vs. alkaline pH and Mn-dependent reactivity differences in the expression platform. A-B.** TECprobe-VL DMS reactivity matrices for *alx* folding intermediates, which were purified and equilibrium refolded at pH 7.2 ± 1 mM Mn (**A**) or pH 8.5 ± 1 mM Mn (**B**). Reactivities shown were normalized with a single, whole-dataset calculated normalization factor. Data are from two independent replicates that were concatenated and analyzed together. **C.** Plots showing comparison of A137 transcript length- and Mn-dependent reactivity changes in co-transcriptionally folded vs. equilibrium refolded intermediates at the designated pH ± 1 mM Mn. DMS data for 0 mM Mn (orange) and 1 mM Mn (purple) from Fig. 7A, SI Fig. 15A, SI Fig. 16A, and SI Fig. 17A-B.

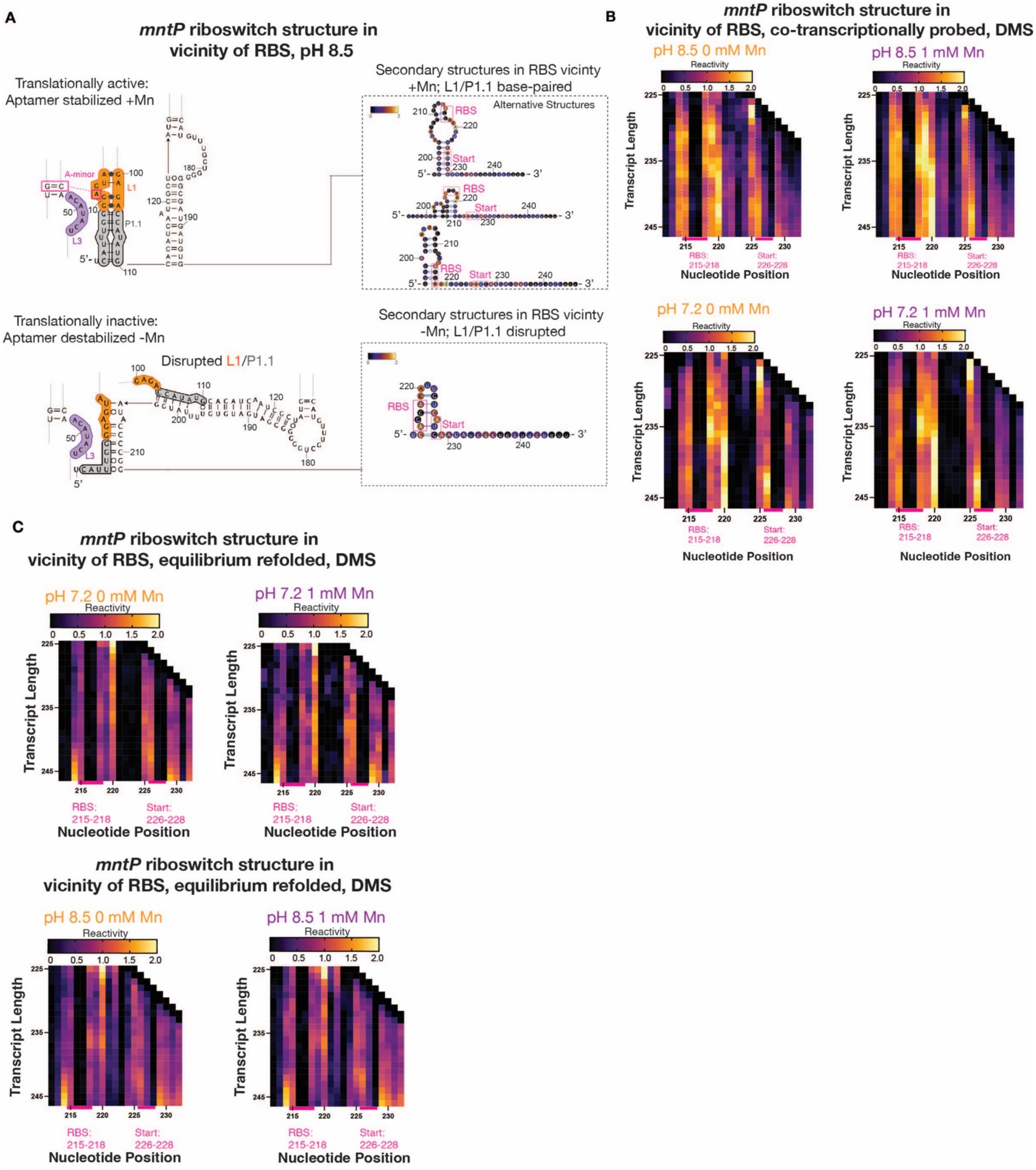


**Supplementary Figure 18. DMS reactivities of *mntP* RBS nt in co-transcriptionally folded and equilibrium refolded intermediates. A.** (Top panel) Truncated secondary structure prediction of full-length, Mn-bound *mntP* riboswitch, with L1 and P1.1 stabilized. Adjacent are secondary structure models for *mntP* expression platform structure at transcript length 246 nt, which are colored by DMS reactivity at pH 8.5 +Mn. (Bottom panel) Truncated secondary structure prediction of full-length, Mn-unbound *mntP* riboswitch, with L1 and P1.1 disrupted. Adjacent is a secondary structure model for *mntP* expression platform structure at transcript length 246 nt, which is colored by DMS reactivity at pH 8.5 -Mn. **B.** TECprobe-VL DMS reactivity matrix for co-transcriptionally folded *mntP* intermediates at pH 7.2 and 8.5 ± 1 mM Mn., windowing over transcript lengths containing the RBS and start codon. Reactivities shown were normalized with a single, whole-dataset calculated normalization factor. Data are from two independent replicates that were concatenated and analyzed together. **C.** TECprobe-VL DMS reactivity matrix for equilibrium refolded *mntP* intermediates at pH 7.2 and 8.5 ± 1 mM Mn., windowing over transcripts lengths containing the RBS and start codon nt. Reactivities shown were normalized with a single, whole-dataset calculated normalization factor. Data are from two independent replicates that were concatenated and analyzed together.

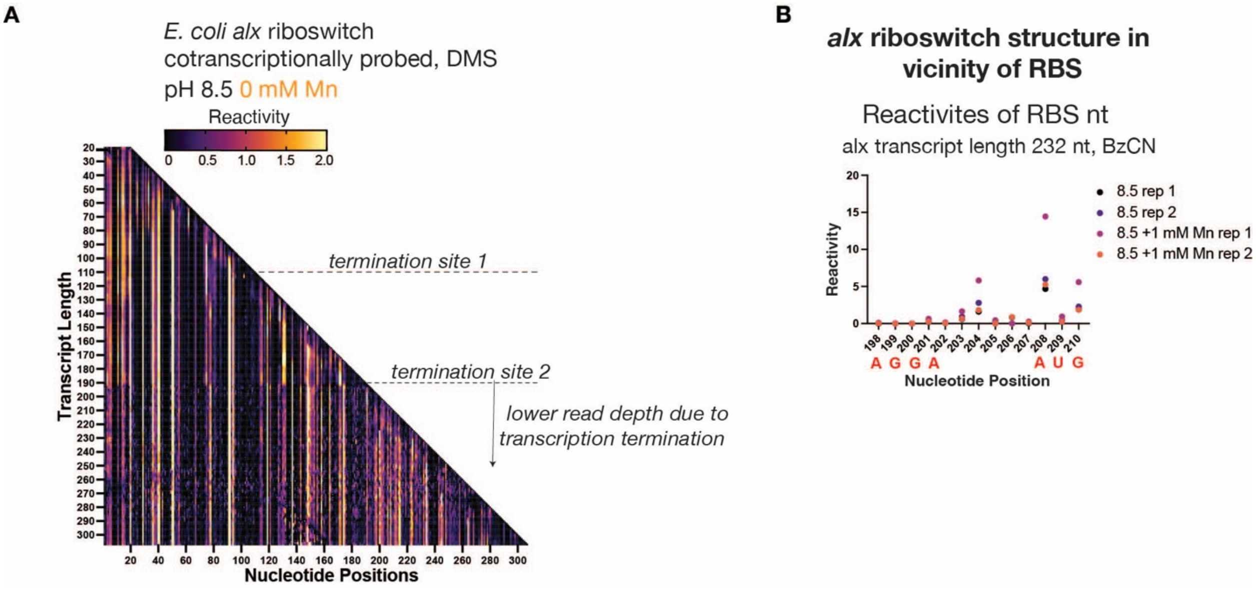


**Supplementary Figure 19. Dropoff in sequencing depth following second *alx* intrinsic termination site and co-transcriptional SHAPE probing of full-length *alx* to capture RBS reactivities A.** TECprobe-VL DMS reactivity matrix for *alx* co-transcriptionally folded intermediates at pH 8.5. Intrinsic transcription termination sites are annotated adjacent to the heatmap, showing a clear drop in reactivity data quality following the second intrinsic termination site. Reactivities shown were normalized with a single, whole-dataset calculated normalization factor. Data are from seven independent replicates that were concatenated and analyzed together. **B.** Plot showing BzCN reactivities of *alx* RBS and start codon nt in the full-length, co-transcriptionally folded riboswitch at pH 8.5 ± 1 mM Mn, corresponding to transcript length 232 nt. Two independent replicates were performed for the ±Mn conditions.

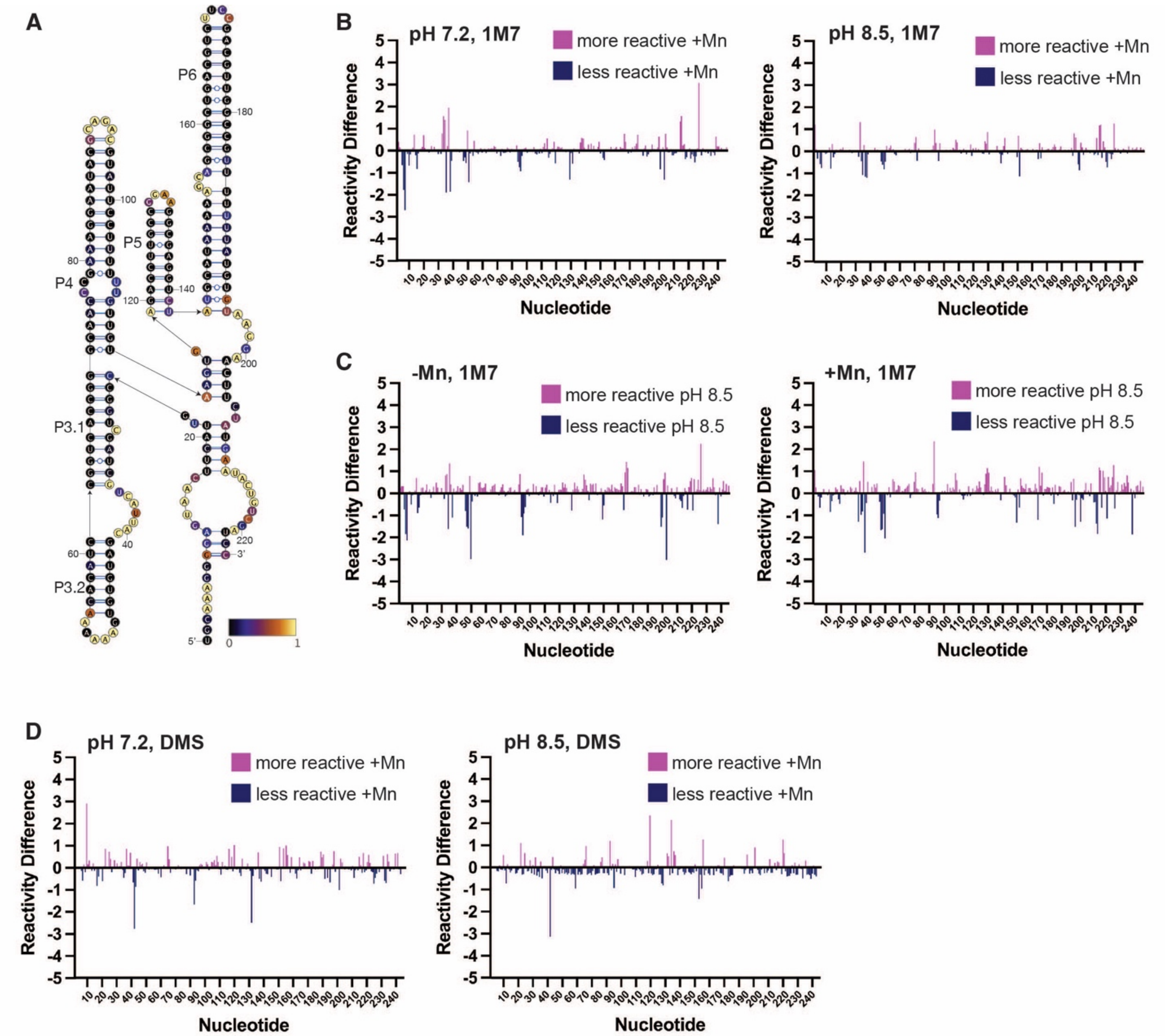


**Supplementary Figure 20. Structure probing of equilibrium-refolded *alx* full-length riboswitch RNA. A.** Secondary structure of equilibrium-refolded *alx* full-length RNA modeled based on reactivity with 1M7 at pH 7.2 -Mn. **B.** Individual nucleotide reactivity differences when modified with 1M7 to compare +Mn vs. -Mn, measured at both pH 7.2 and 8.5. **C.** Individual nucleotide reactivity differences at neutral pH (7.2) vs. alkaline pH (8.5), measured both in the presence and absence of Mn. **D.** Individual nucleotide reactivity differences when modified with DMS to compare +Mn vs. -Mn, measured at both pH 7.2 and 8.5.

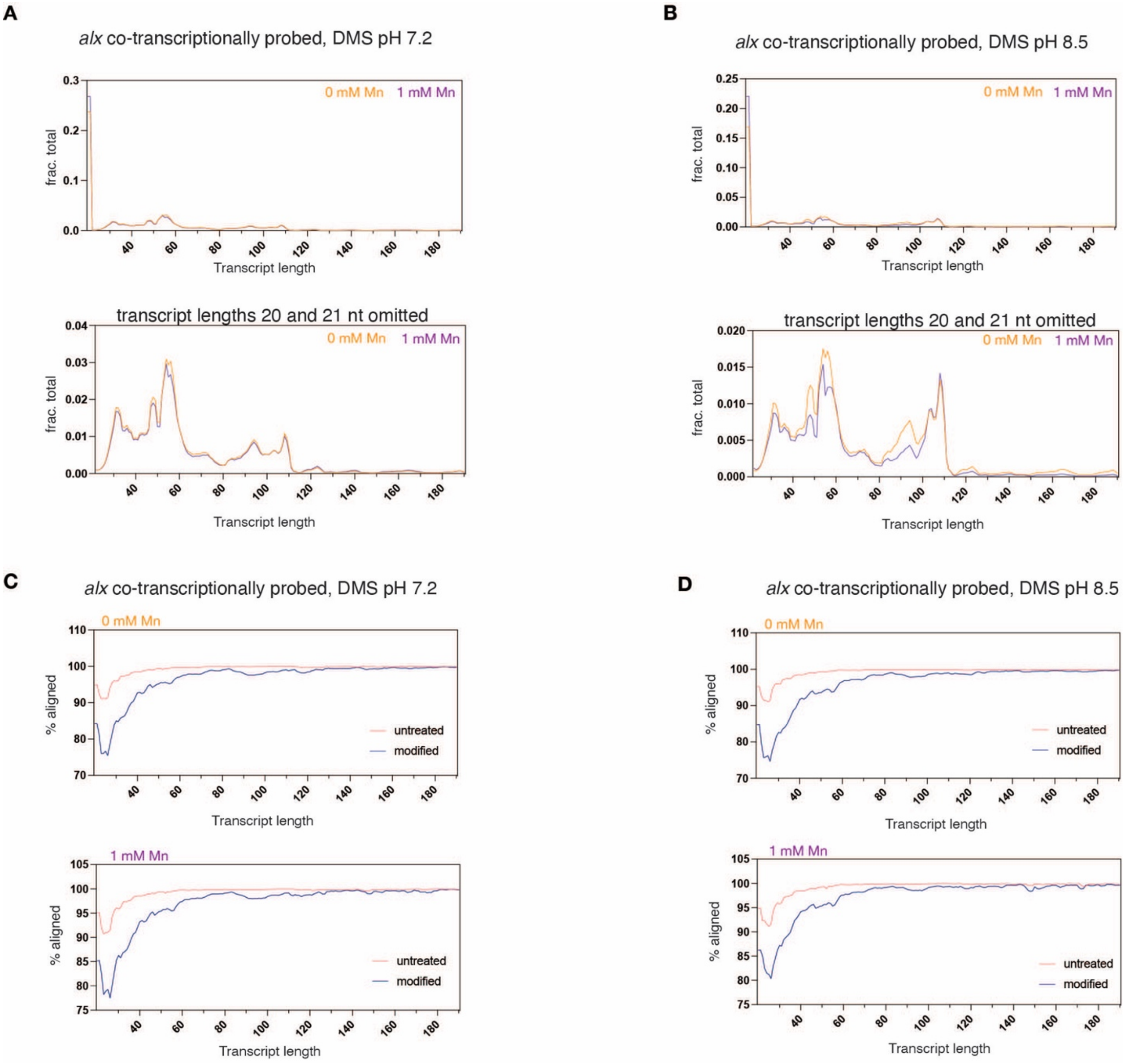


**Supplementary Figure 21. Aligned reads and alignment rates for co-transcriptional DMS probing of the *alx* riboswitch. A-B.** (Top panel) Plot showing fraction of aligned reads mapped to each transcript length. (Bottom panel) Plot showing fraction of aligned reads mapped to each transcript length, with transcript lengths 20 and 21 nt omitted to better visualize the fraction of aligned reads for transcript lengths 22-190 nt. **C-D.** Percentage of reads retained after splitting those that aligned to each intermediate transcript length.

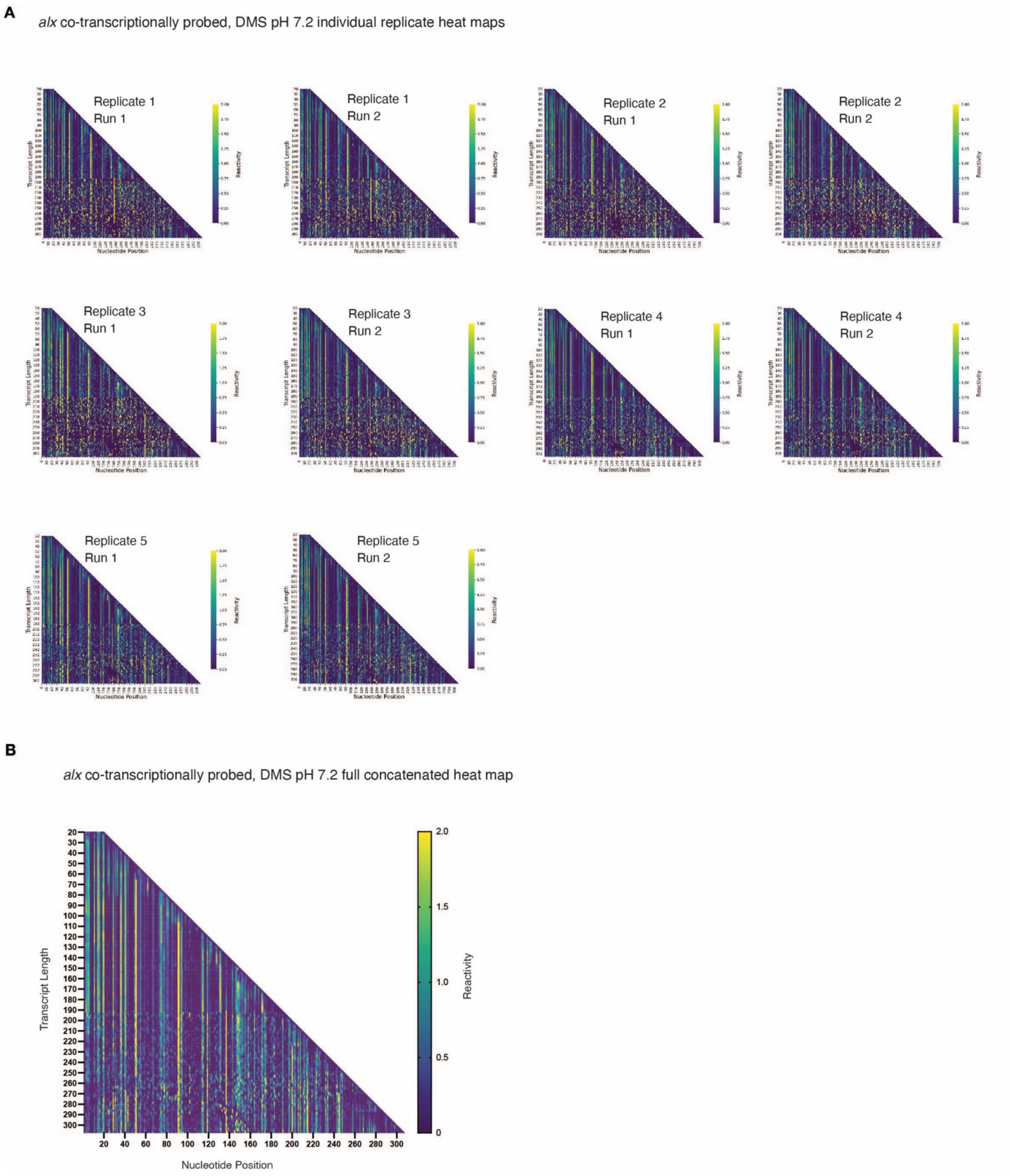


**Supplementary Figure 22. Reactivity matrices for individual and concatenated replicates for co-transcriptional DMS probing of the *alx* riboswitch at pH 7.2. A.** TECprobe-VL DMS reactivity matrices for *alx* folding intermediates from five independent replicates at pH 7.2. “Run 1” and “Run 2” denote sequencing runs on independent PCR amplifications of the same cDNA library. Reactivities shown were normalized with a single, whole-dataset calculated normalization factor. **B.** TECprobe-VL DMS reactivity matrix for *alx* co-transcriptionally folded intermediates at pH 7.2. Reads from the five independent replicates shown in panel A, which were concatenated and analyzed together.

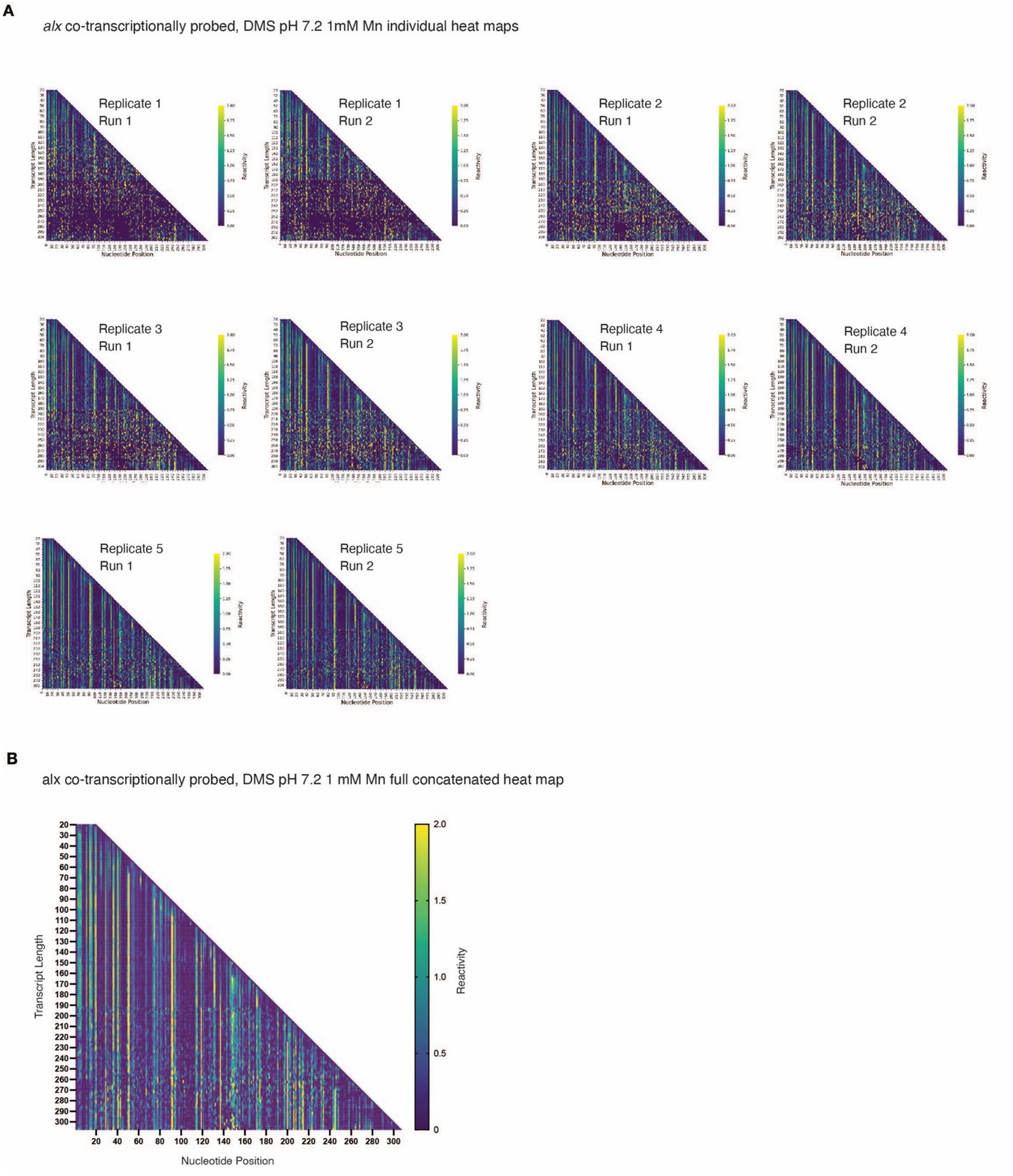


**Supplementary Figure 23. Reactivity matrices for individual and concatenated replicates for co-transcriptional DMS probing of the *alx* riboswitch at pH 7.2 +Mn. A.** TECprobe-VL DMS reactivity matrices for *alx* folding intermediates from five independent replicates at pH 7.2 + 1 mM Mn. “Run 1” and “Run 2” denote sequencing runs on independent PCR amplifications of the same cDNA library. Reactivities shown were normalized with a single, whole-dataset calculated normalization factor. **B.** TECprobe-VL DMS reactivity matrix for *alx* co-transcriptionally folded intermediates at pH 7.2 + 1 mM Mn. Reads from the five independent replicates shown in panel A, which were concatenated and analyzed together.

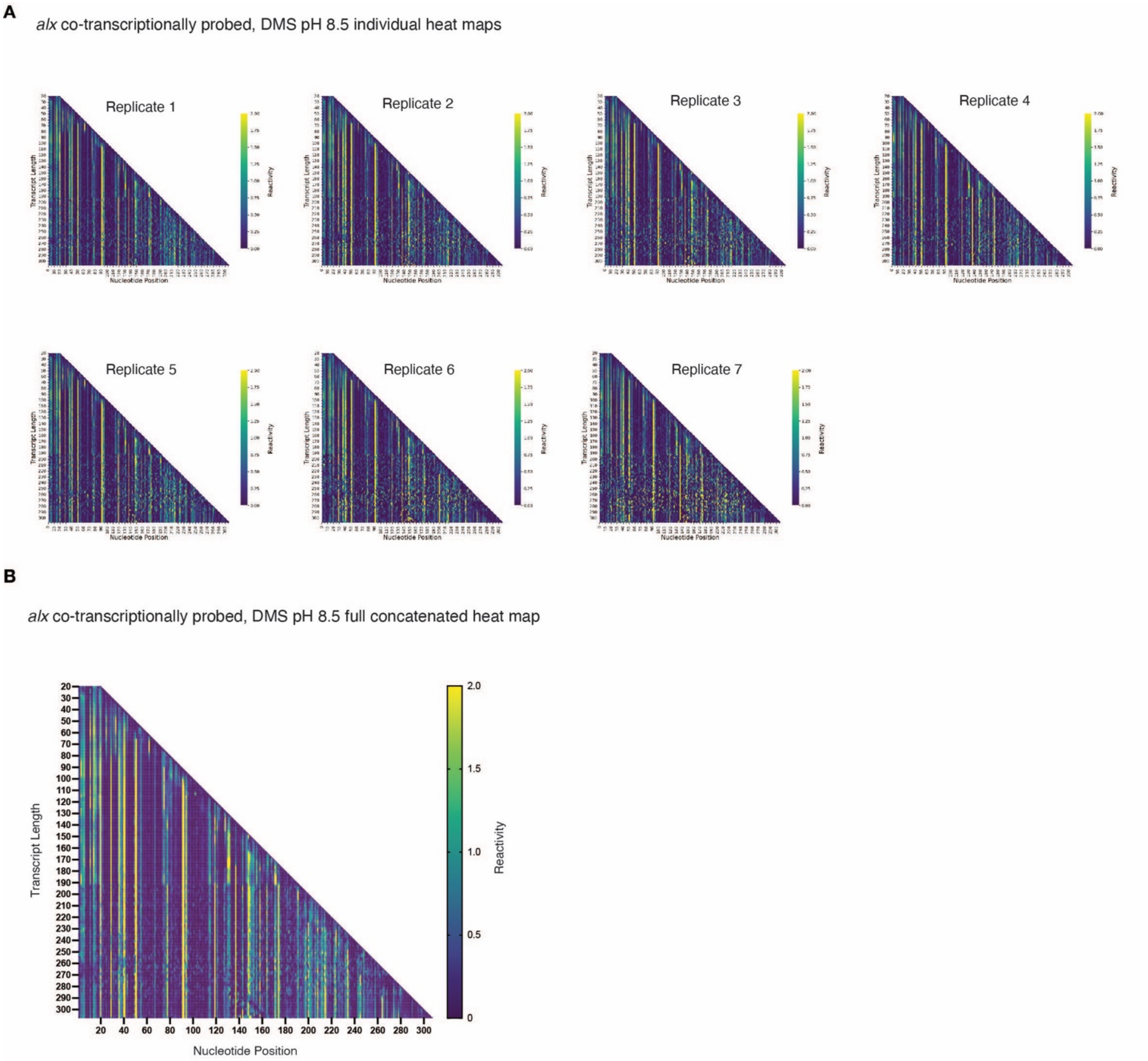


**Supplementary Figure 24. Reactivity matrices for individual and concatenated replicates for co-transcriptional DMS probing of the *alx* riboswitch at pH 8.5. A.** TECprobe-VL DMS reactivity matrices for *alx* folding intermediates from seven independent replicates at pH 8.5. Reactivities shown were normalized with a single, whole-dataset calculated normalization factor. **B.** TECprobe-VL DMS reactivity matrix for *alx* co-transcriptionally folded intermediates at pH 8.5. Reads from the seven independent replicates shown in panel A, which were concatenated and analyzed together.

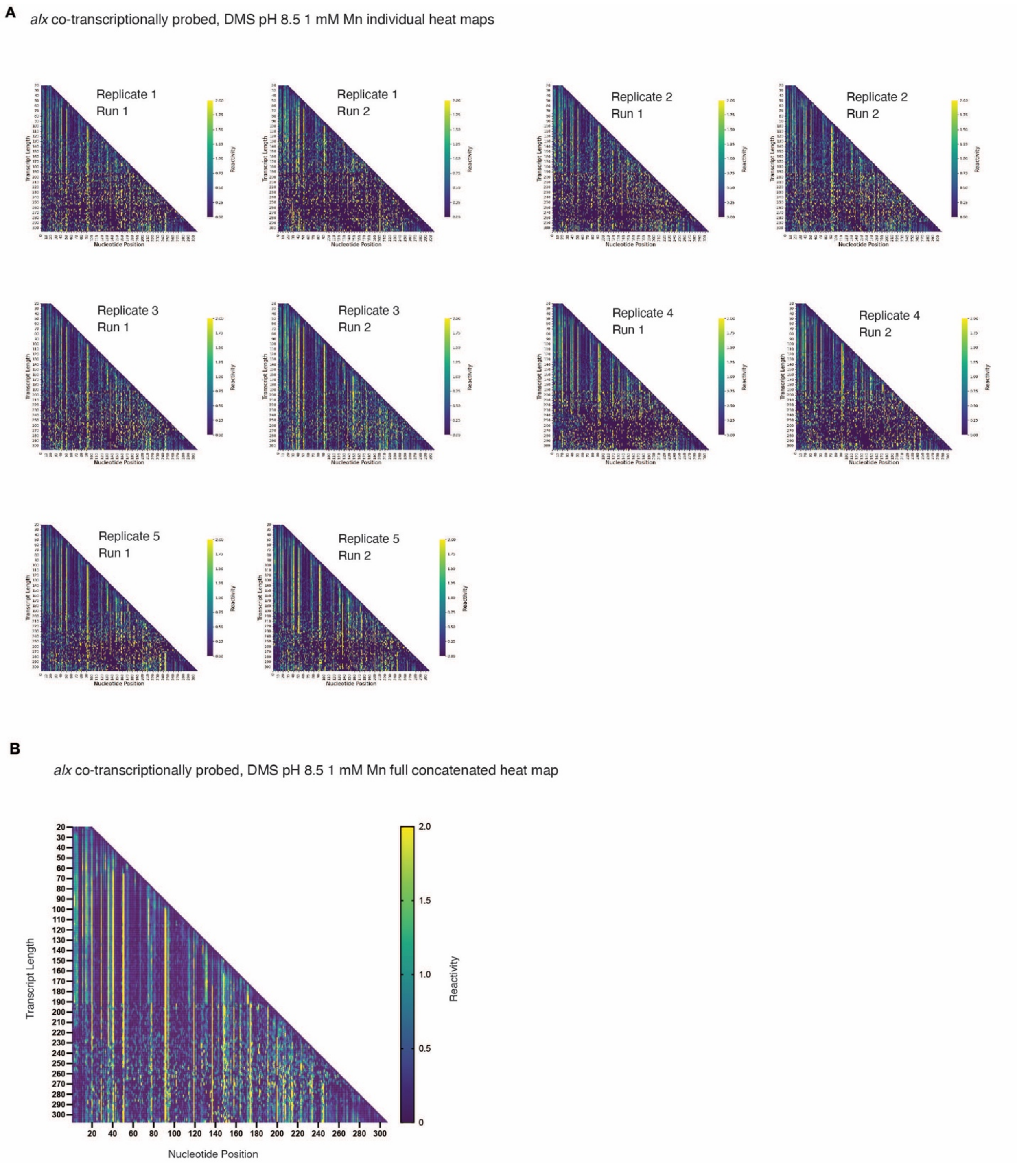


**Supplementary Figure 25. Reactivity matrices for individual and concatenated replicates for co-transcriptional DMS probing of the *alx* riboswitch at pH 8.5 +Mn. A.** TECprobe-VL DMS reactivity matrices for *alx* folding intermediates from five independent replicates at pH 8.5 + 1 mM Mn. “Run 1” and “Run 2” denote sequencing runs on independent PCR amplifications of the same cDNA library. Reactivities shown were normalized with a single, whole-dataset calculated normalization factor. **B.** TECprobe-VL DMS reactivity matrix for *alx* co-transcriptionally folded intermediates at pH 8.5 + 1 mM Mn. Reads from the five independent replicates shown in panel A, which were concatenated and analyzed together.

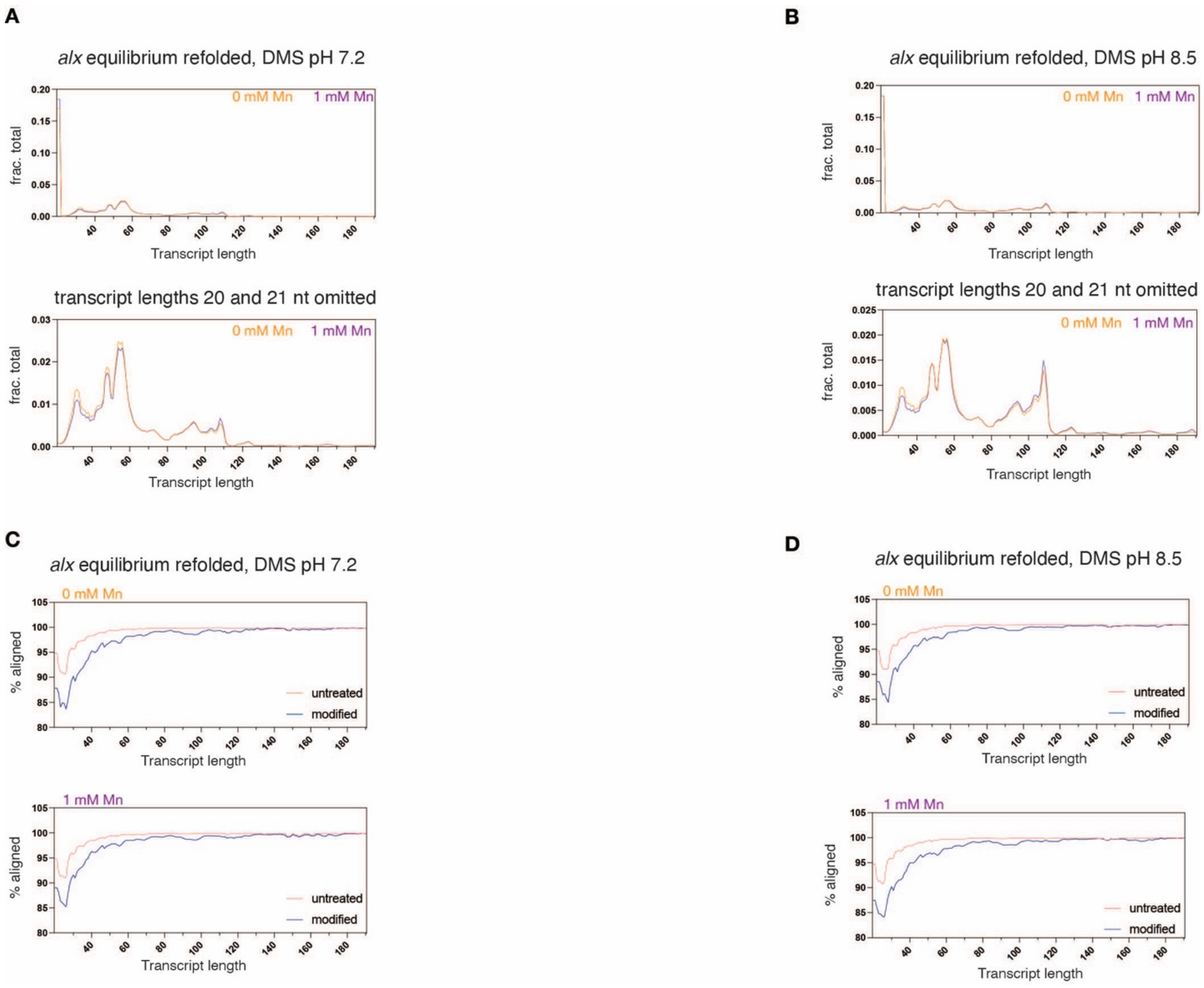


**Supplementary Figure 26. Aligned reads and alignment rates for DMS probing of *alx* equilibrium refolded intermediates. A-B.** (Top panel) Plot showing fraction of aligned reads mapped to each transcript length. (Bottom panel) Plot showing fraction of aligned reads mapped to each transcript length, with transcript lengths 20 and 21 nt omitted to better visualize the fraction of aligned reads for transcript lengths 22-190 nt. **C-D.** Percentage of reads retained after splitting those that aligned to each intermediate transcript length.

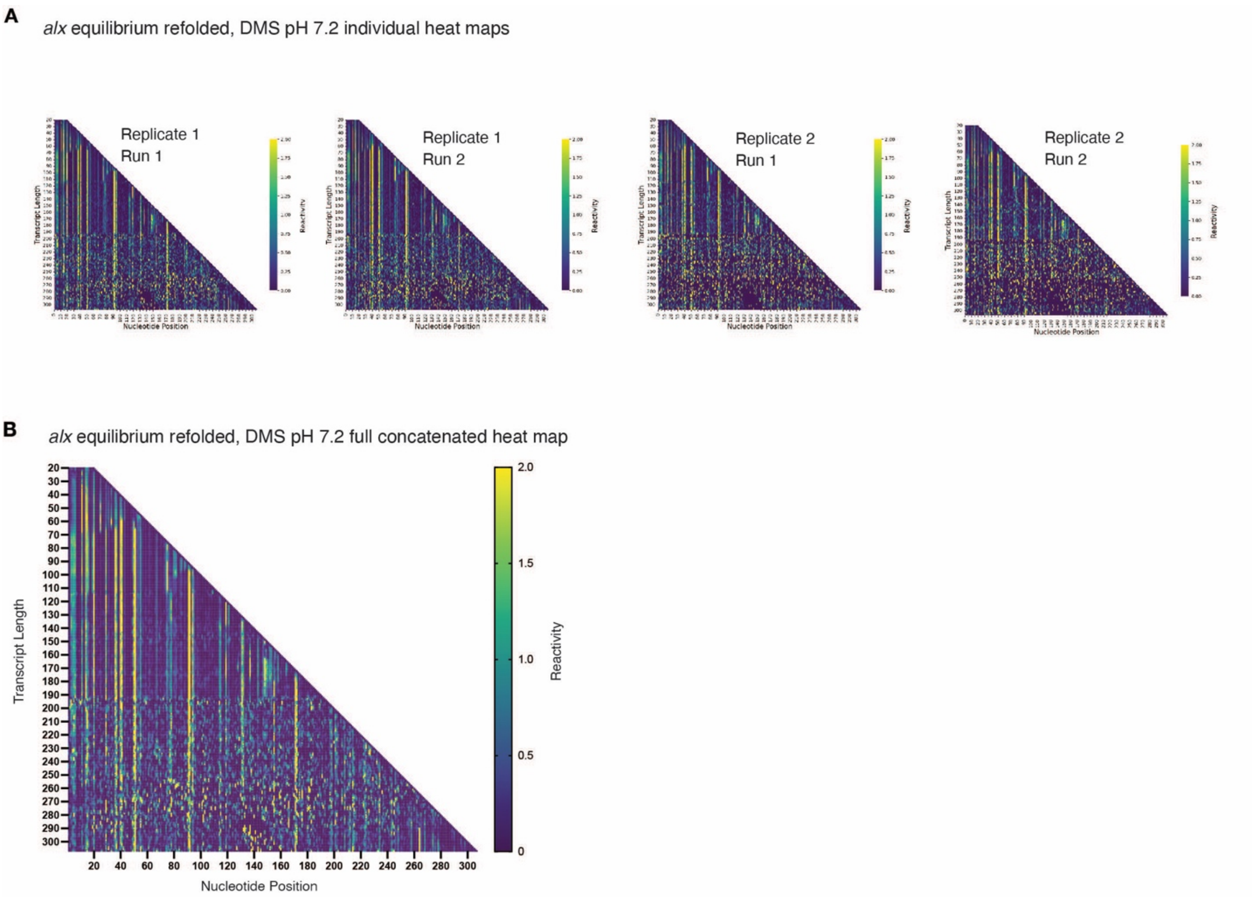


**Supplementary Figure 27. Reactivity matrices for individual and concatenated replicates for equilibrium DMS probing of the *alx* riboswitch at pH 7.2. A.** TECprobe-VL DMS reactivity matrices for *alx* folding intermediates from two independent replicates at pH 7.2. "Run 1" and "Run 2" denote sequencing runs on independent PCR amplifications of the same cDNA library. Reactivities shown were normalized with a single, whole-dataset calculated normalization factor. **B.** TECprobe-VL DMS reactivity matrix for *alx* equilibrium refolded intermediates at pH 7.2. Reads from the two independent replicates shown in panel A, which were concatenated and analyzed together.

**A**

*alx* equilibrium refolded, DMS pH 7.2 1mM Mn individual heat maps

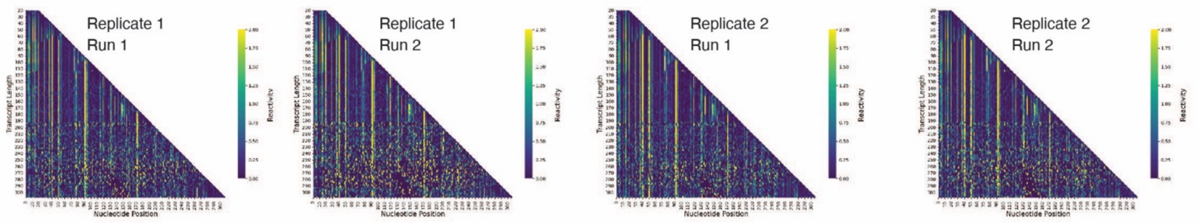


**B**

*alx* equilibrium refolded, DMS pH 7.2 1mM Mn full concatenated heat map

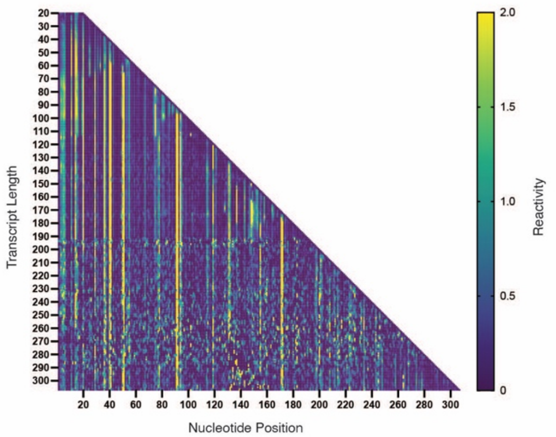


**Supplementary Figure 28. Reactivity matrices for individual and concatenated replicates for equilibrium DMS probing of the *alx* riboswitch at pH 7.2 +Mn. A.** TECprobe-VL DMS reactivity matrices for *alx* folding intermediates from two independent replicates at pH 7.2 + 1 mM Mn. "Run 1" and "Run 2" denote sequencing runs on independent PCR amplifications of the same cDNA library. Reactivities shown were normalized with a single, whole-dataset calculated normalization factor. **B.** TECprobe-VL DMS reactivity matrix for *alx* equilibrium refolded intermediates at pH 7.2 + 1 mM Mn. Reads from the two independent replicates shown in panel A, which were concatenated and analyzed together.

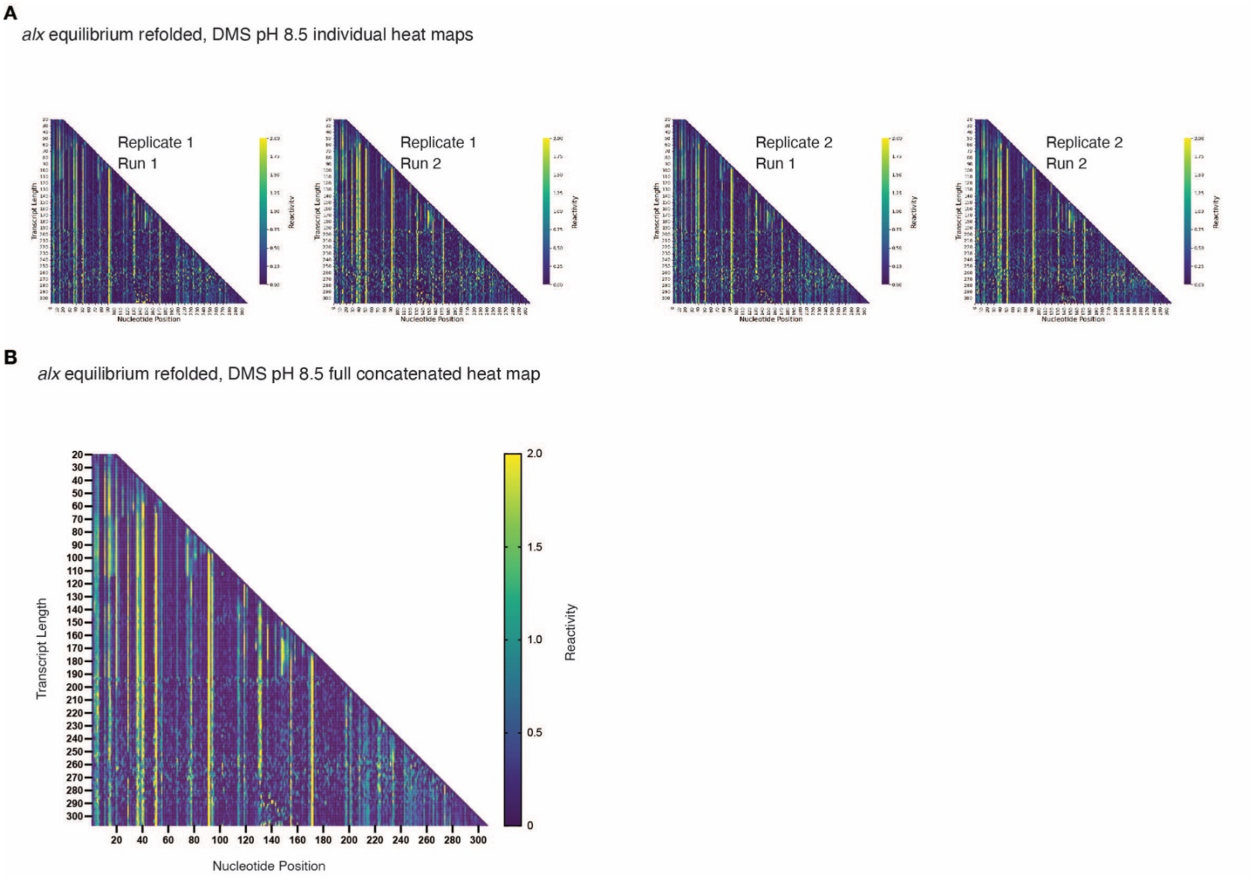


**Supplementary Figure 29. Reactivity matrices for individual and concatenated replicates for equilibrium DMS probing of the *alx* riboswitch at pH 8.5. A.** TECprobe-VL DMS reactivity matrices for *alx* folding intermediates from two independent replicates at pH 8.5. “Run 1” and “Run 2” denote sequencing runs on independent PCR amplifications of the same cDNA library. Reactivities shown were normalized with a single, whole-dataset calculated normalization factor. **B.** TECprobe-VL DMS reactivity matrix for *alx* equilibrium refolded intermediates at pH 8.5. Reads from the two independent replicates shown in panel A, which were concatenated and analyzed together.

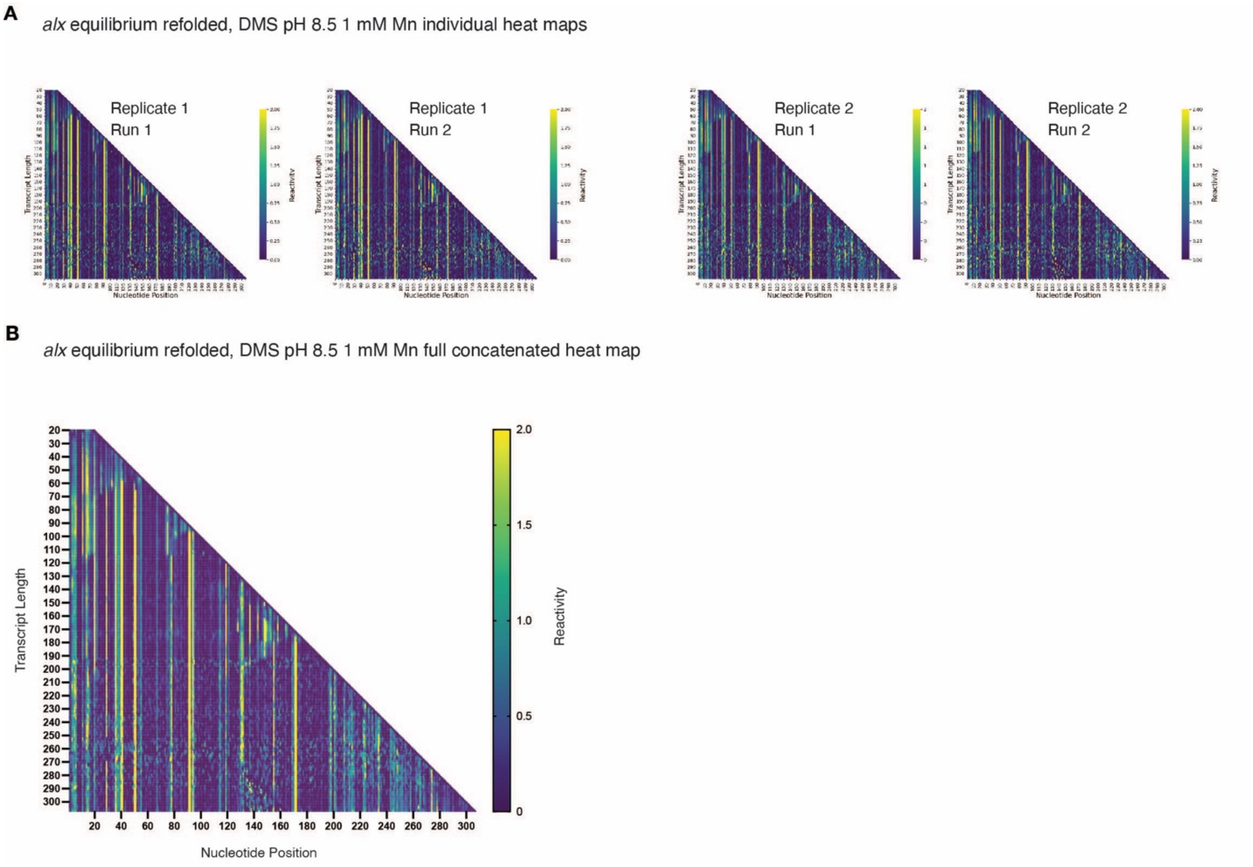


**Supplementary Figure 30. Reactivity matrices for individual and concatenated replicates for equilibrium DMS probing of the *alx* riboswitch at pH 8.5 +Mn. A.** TECprobe-VL DMS reactivity matrices for *alx* folding intermediates from two independent replicates at pH 8.5 + 1 mM Mn. “Run 1” and “Run 2” denote sequencing runs on independent PCR amplifications of the same cDNA library. Reactivities shown were normalized with a single, whole-dataset calculated normalization factor. **B.** TECprobe-VL DMS reactivity matrix for *alx* equilibrium refolded intermediates at pH 8.5 + 1 mM Mn. Reads from the two independent replicates shown in panel A, which were concatenated and analyzed together.

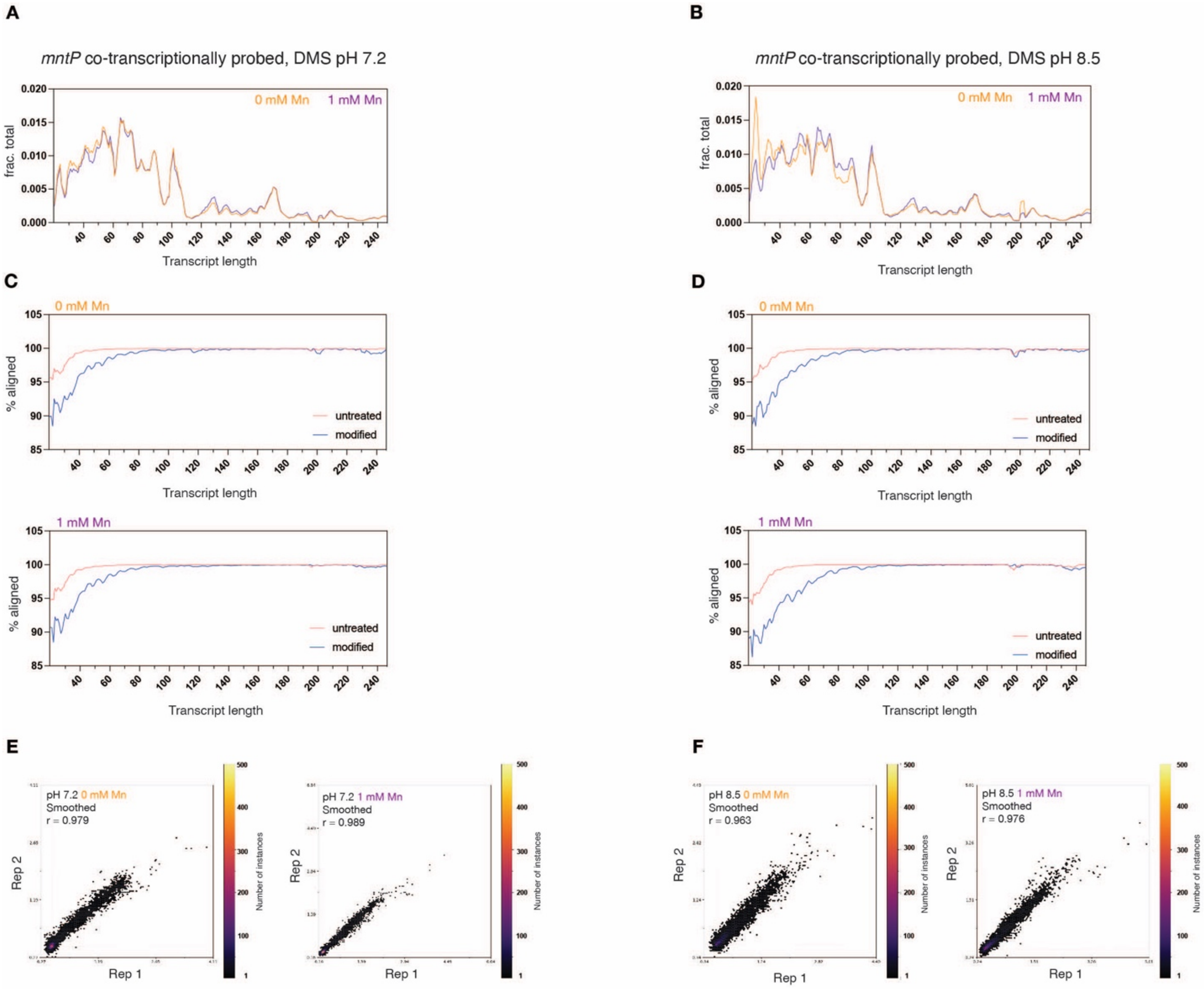


**Supplementary Figure 31. Aligned reads and alignment rates for co-transcriptional DMS probing of the *mntP* riboswitch. A-B.** Plot showing fraction of aligned reads mapped to each transcript length. **C-D.** Percentage of reads retained after splitting those that aligned to each intermediate transcript length. **E-F.** Hexbin plots comparing the DMS reactivity of replicates.

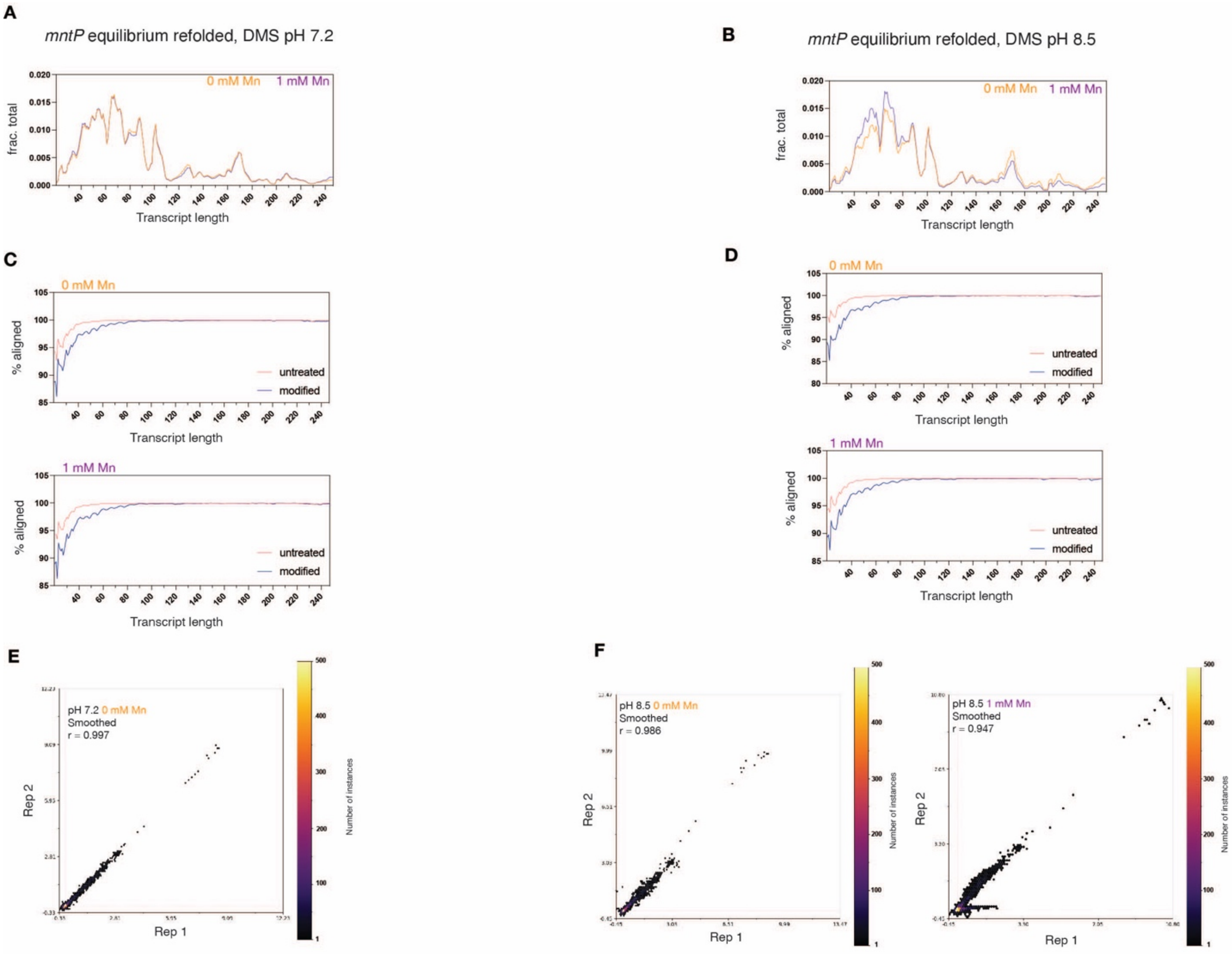


**Supplementary Figure 32. Aligned reads and alignment rates for equilibrium DMS probing of the *mntP* riboswitch. A-B.** Plot showing fraction of aligned reads mapped to each transcript length. **C-D.** Percentage of reads retained after splitting those that aligned to each intermediate transcript length. **E-F.** Hexbin plots comparing the DMS reactivity of replicates. No hexbin plot is shown for the pH 7.2 + 1 mM Mn condition because only one replicate was analyzed for this condition.

### Supplemental Notes

#### Supplementary Note 1

*pH effect on the refolded alx aptamer structure.* Comparison of the nucleotide (nt) reactivities as a function of refolding pH revealed clear flexibility differences in L3 (Fig. 2C). The strictly conserved adenosine (A37) that coordinates Mn is less accessible for modification by both DMS and 1M7 at alkaline pH, despite the absence of Mn (Fig. 2C and SI Fig. 1E). Similarly, most L3 nt exhibit lower reactivities at alkaline vs neutral pH when modified with 1M7; however, the C36 and U38 that neighbor A37 are both more reactive at alkaline pH (Fig. 2C). Also, the L1 conserved adenosine (A11) that forms the cross-helix A-minor interaction becomes less reactive towards 1M7 at alkaline pH -Mn (Fig. 2C). Interestingly, this A11 pH-dependent reactivity difference is abolished when RNA is modified with DMS (SI Fig. 1E). Together, these data indicate that alkaline pH alters the reactivity of key nucleotides in the *alx* metal-sensing core, which could be due to a local pH effect on L1/L3 structure or a global pH-dependent trend in aptamer dynamics.

*Mn effect on the refolded alx aptamer structure.* Reactivity analysis of 1M7-modified RNA revealed that L3 generally becomes less flexible +Mn, consistent with Mn binding, except for U38, which remains more flexible +Mn regardless of pH (SI Fig. 1D). U39 is also more reactive towards 1M7 +Mn at neutral pH (SI Fig. 1D). For DMS-modified aptamer RNA, most L3 nt do not show significant Mn-dependent reactivity differences; however, consistent with 1M7 reactivity analysis, U38 and U39 are both more flexible at neutral pH +Mn (Fig. 1E). The crystal structure of a $Cd^{2+}$-bound *L. lactis* Mn-sensing aptamer mutated to have the *alx* L3 sequence shows that metal ion binding flips U38 and U39 out of the nucleobase stack formed by the A-minor motif, explaining the observed reactivity increase in reactivity when Mn is present(1).

Consistent with the role of A37 in directly coordinating Mn, no pH-dependent change in reactivity is observed +Mn when RNA is modified with either 1M7 or DMS (SI Fig. 1D-

E). Further, in the presence of Mn, the pH-dependent change in reactivity towards 1M7 for A11, which forms the cross-helix A-minor interaction, is abolished (Fig. 2C). This finding agrees with published smFRET studies on the related *X. oryzae* Mn-sensing aptamer, which demonstrated that Mn increases docking of the two helical legs(2). Since A11 is a key player in this docking via the A-minor interaction, it is expected to be more frequently buried in the metal-sensing core, and thus less reactive towards a chemical probe +Mn.

Besides A11, L1 nt in 1M7-modified RNA are either more reactive or show minor reactivity differences +Mn at neutral pH; however, upon shifting to alkaline pH, L1 nt become less reactive +Mn (SI Fig. 1D). When aptamer RNA was DMS-probed at neutral pH, most L1 nucleotides are less reactive +Mn; however, upon a shift to alkaline pH, Mn-dependent reactivity differences are lessened, with G10 and G120 showing increased reactivity (SI Fig. 1E). Taken together, these data show that Mn alone reduces the flexibility of critical Mn-sensing nucleotides in L1 and L3 of the *alx* aptamer and are consistent with prior structural study of Mn-bound aptamers(1–3).

**Supplementary Note 2**

*Mn-enhanced RNA polymerase (RNAP) pause longevity during mntP aptamer synthesis*. As a further support for Mn interacting with RNA folding intermediates, we mapped multiple RNAP pauses during *mntP* aptamer synthesis (nt 86-90) that are enhanced by Mn (SI Fig. 4A-B). Similar enhancement of pauses within the *mntP* aptamer was achieved by addition of NusA in the absence of Mn (SI Fig. 4B), pointing to RNA hairpin stabilization of paused RNAP conformation at these sites. Thus, these pauses are likely prolonged via allosteric effect of Mn-RNA on the RNAP conformation akin to the allosteric effects of nascent RNA hairpins(4, 5). The presence of these hairpin-stabilized pauses is further supported by RNA structure models constrained by DMS reactivities (stabilized by the P3.1 hairpin based on pause locations, SI Fig. 4C). All pause sites described above were mapped by scaffold-based transcription assays, which provided single-nt precision for mapping pause locations (SI Fig. 5A-C). Taken together, these data suggest that the hairpin-stabilized pauses during aptamer synthesis mediate sampling of divalents in the environment by the *mntP* RNA folding intermediates. Recent smFRET study on the *L. lactis* Mn-sensing riboswitch also identified a pause site enhanced either in the presence of Mn or NusA, with the pause in this case positioned at the end of aptamer folding to stabilize P1.1 switch helix(6).

**Supplementary Note 3**

*Some but not all co-transcriptional folding features are preserved in unfolded-refolded RNA intermediates.* To identify the intermediates whose folding is truly controlled by the transcription kinetics vs. thermodynamic stability of the folds, we applied a modified TECprobe-VL workflow wherein intermediate RNA transcripts are extracted from the stalled transcription elongation complexes and equilibrium refolded before DMS probing (SI Fig. 6A). As with co-transcriptionally folded intermediates, we observed transient, low-$T_m$ stem-loops that form and resolve once the full aptamer sequence is present (SI Fig. 6B). We observed similar Mn-dependent differences in reactivity in equilibrium refolded as the co-transcriptionally folded intermediates for A48, U54, and A103 (SI Fig. 6C). Despite the L3 being single-stranded throughout the aptamer folding in both the equilibrium refolded and co-transcriptional intermediates, the reactivity of A48 shows a marked increase ~10 nt before the aptamer folds (SI Fig. 6C). This suggests that once the first 100 nt of the aptamer are available for base-pairing, a structural rearrangement occurs to reposition the L3, resulting in increased reactivity of A48 overall. A key folding transition that likely prompts this structural rearrangement is the pairing of P1.2 in the right leg, where nt 15-18 at the 5' end pair with nt 96-99. Once P1.2 folds, the DMS reactivity profile for transcript length 100 nt largely resembles that of the folded aptamer, with only L1 and P1.1 absent due to lack of their pairing partners (SI Fig. 6D-E). Thus, formation of P1.2 initiates folding of the right leg, which finishes folding once the 5' end of the aptamer transcript (nt 2-18) pairs with the nt that emerge later in transcription (nt 96-110, Fig. 3B). We hypothesize that folding of P1.2 positions nt 2-14 in closer proximity to the partially folded right leg to facilitate pairing of nt 2-14 with nt 100-110 once they emerge from RNAP, permitting efficient completion of the aptamer and metal-sensing core folding, and thus a rapid response to ligand. Further, repositioning of nt 1-14 likely brings these nucleotides into an even closer proximity to L3, resulting in nucleation of the cross-helix A-minor with A11 stacked atop A48 and explaining the Mn-dependent changes in A48 reactivity at this point.

For U54, Mn-dependent differences in reactivity emerge before the transcript is 100 nt, indicating that some positions in the L3 loop can begin sampling for metal even before the right leg in the aptamer begins folding (SI Fig. 6C). Interestingly, the magnitude of the U54 reactivity difference ±Mn is higher for the early equilibrium refolded intermediates (transcript lengths ~60-100 nt) compared to the co-transcriptionally folded intermediates (SI Fig. 6C). However, past transcript length ~100 nt, the magnitude of the U54 reactivity difference ±Mn is higher for the co-transcriptionally folded intermediates, with U54 being more reactive in the absence of ligand (SI Fig. 6C). This indicates that the L3 is more flexible in co-transcriptionally folded intermediates once the metal-sensing core begins folding, resulting in higher reactivity at U54. Additionally, past transcript lengths ~110-120 nt, A103 demonstrates higher reactivity –Mn, with the magnitude of the reactivity difference larger for the co-transcriptionally folded intermediates (SI Fig. 6C).

**Supplementary Note 4**

*alx L3 nt show distinct Mn-dependent reactivity trends in equilibrium refolded intermediates*. Similar to *mntP*, we observed Mn-dependent reactivity differences for equilibrium refolded *alx* intermediates (SI Fig. 8A). First, U35 reactivity trends higher +Mn for equilibrium refolded intermediates starting at transcript length ~50 nt. Curiously, the magnitude of the reactivity difference ±Mn is greater for the early equilibrium refolded intermediates (transcript lengths ~60-115 nt, SI Fig. 8B) than for later intermediates, indicating a potential structural change (e.g., a loss of a tertiary RNA contact) induced by equilibrium refolding that affects L3 dynamics in these early intermediates. Additionally, A37 in the *alx* aptamer demonstrates higher reactivity –Mn for equilibrium refolded intermediates, mimicking the trend observed for co-transcriptionally folded intermediates.

However, the magnitude of the Mn-dependent reactivity difference for A37 is greater for equilibrium refolded intermediates across transcript lengths ~60-115 nt and ~135-155 nt (SI Fig. 8B). Lastly, C41 demonstrates higher reactivity +Mn for equilibrium refolded intermediates, with a larger Mn-dependent reactivity difference for equilibrium refolded intermediates spanning transcript lengths ~70 nt-110 nt (SI Fig. 8B). Taken together, these data indicate a potential structural change induced by equilibrium refolding that affects L3 dynamics, resulting in differing reactivity patterns compared to co-transcriptionally folded intermediates. Further, these data provide additional pieces of evidence that aptamer intermediates undergo Mn-dependent structural changes in L3.

**Supplementary Note 5**

*pH-dependent reactivity trends in alx and mntP L3 nt in equilibrium refolded intermediates*. We next assessed pH differences in L3 nucleotide reactivities for *alx* and *mntP* equilibrium refolded aptamer intermediates, after verifying that the global folding transitions in the aptamers are unaffected by pH (SI Fig. 10A-B). First, *alx* A37 shows a clear Mn-dependent difference in reactivity at both pH from transcript lengths ~60-115 nt, with lower reactivity in general +Mn (SI Fig. 10C). This trend is different from that seen for A37 in co-transcriptionally folded intermediates, where the lowest reactivity overall was observed for the pH 8.5 +Mn condition throughout aptamer folding (Fig. 5B). Additionally, positions C36, A40 and C41 in *alx* equilibrium refolded intermediates do not show the same increased reactivity at pH 8.5 ±Mn as described for co-transcriptionally folded intermediates (SI Fig.9C and SI Fig. 10C). These data support that an alkaline pH shift uniquely affects the structure of co-transcriptionally folded *alx* aptamer intermediates.

The *mntP* L3 also demonstrates different reactivity trends in equilibrium refolded aptamer intermediates. A48 shows no clear pH difference in reactivity (SI Fig. 10D); this is in stark contrast to co-transcriptionally folded aptamer intermediates, where A48 is more reactive at alkaline pH (Fig. 5C). This result indicates that equilibrium refolding of *mntP* aptamer intermediates alters the orientation of A48, the A11 stacking partner, such that pH no longer alters the accessibility of its Watson-Crick face for DMS methylation.

The absolutely conserved adenosine in *mntP* L3 (A50) is more reactive at neutral pH ±Mn for the duration of aptamer folding (SI Fig. 10D), which was not observed for co-transcriptionally folded intermediates (Fig. 5C). These data support that equilibrium refolding of *mntP* aptamer intermediates repositions the L3 loop and changes how pH affects L3 structure.

*pH-dependent reactivity trends in alx and mntP L1 nt in co-transcriptionally folded vs. equilibrium refolded intermediates*. Beyond L3, both the *alx* and *mntP* aptamers also exhibit curious pH-dependent reactivity differences in the absolutely conserved L1. A101 in the *mntP* aptamer is more reactive at alkaline pH whether or not Mn is present (SI Fig. 11A). A103 in *mntP* is generally more reactive –Mn at both pH, with the absolute reactivity values greater at pH 7.2 ±Mn (SI Fig. 11A). At the analogous positions in the *alx* aptamer (A119 and A121) we observed different reactivity trends. In the absence of Mn, A119 in *alx* is more reactive at alkaline pH during aptamer folding. In the presence of Mn, however, there is a clear pH-dependent divergence in the *alx* A119 reactivity at the end of aptamer folding, with a peak in reactivity observed at pH 7.2 and a drop in reactivity at pH 8.5 (SI Fig. 11A). A121 exhibited similar trends to those described above for A119, with a stark divergence in reactivity between the pH 7.2 +Mn and pH 8.5 +Mn conditions towards the end of aptamer folding (SI Fig. 11A). These data solidify that a shift to alkaline pH exerts distinct structural effects on the *alx* and *mntP* aptamers.

Lastly, we assessed single nucleotide reactivity trajectories for *alx* and *mntP* L1 nucleotides in equilibrium refolded intermediates and compared them to their co-transcriptionally folded counterparts. For *mntP* A101, there is no clear pH-dependent reactivity trend (SI Fig. 11B), which differs from co-transcriptionally folded intermediates where A101 was more reactive at pH 8.5 in general (SI Fig. 11A). A119 and A121 in the *alx* L1 showed no clear pH trends for equilibrium refolded aptamer intermediates. A119 reactivity in general was higher towards the end of aptamer folding in equilibrium refolded transcripts for all conditions tested compared to co-transcriptionally folded transcripts (SI Figs 11A-B), indicating that equilibrium refolding for *alx* aptamer intermediates results in a more dynamic L1. Taken together, these data support that equilibrium refolding alters the structure and dynamics of the metal-sensing core.

**Supplementary Note 6**

*pH- and Mn-dependent changes in mntP aptamer structure during co-transcriptional expression platform folding*. We observed similar reactivity trends in *mntP* expression platform folding at pH 7.2 as pH 8.5 (SI Fig. 12B-C). For example, at pH 7.2, we also observed the transient spike in A11 reactivity at transcript length ~205 nt; however, both the amplitude of this spike and the difference ±Mn is smaller compared to pH 8.5 (SI Fig. 12D). Other curious ligand- and pH-dependent trends stood out for *mntP* nucleotides in L3, L1, and the switch helix P1.1 during expression platform folding. A48 in L3 drops in reactivity at transcript length ~200 nt for all conditions followed by an increase in reactivity at pH 7.2 +Mn and pH 8.5 +Mn compared to the respective –Mn conditions (SI Fig. 12D). This indicates that a global, Mn-dependent structural rearrangement occurs in the aptamer at this point, consistent with the reactivity trend described for A11. Since A11 and A48 are stacking partners (Fig. 6C), it makes sense that their reactivity changes are coupled upon disruption of the cross-helix A-minor interaction. A50 in L3, which coordinates Mn, also exhibits ligand-dependent reactivity differences for the duration of expression platform folding (SI Fig. 12D).

L1 nucleotides besides A11 undergo ligand-dependent structural changes at the riboswitching point. Similar to A11, A14 reactivity spikes towards the end of expression platform folding –Mn (SI Fig. 12D); however, the increase in reactivity occurs ~5-10 nt before the spike in A11 reactivity, indicating that the A14-G100 base-pair in L1 is broken before the cross-helix A-minor is disrupted. A101 in L1 remains generally more reactive at pH 8.5 ±Mn vs. pH 7.2 ±Mn from transcript lengths ~120-190 nt and demonstrates high reactivity in general despite its predicted base-pairing with U13 (SI Fig. 12D). Curiously, at transcript length ~200 nt, a drop in A101 reactivity is observed for all conditions, with the +Mn condition trending higher for the remainder of expression platform folding (SI Fig. 12D). When L1 is disrupted –Mn, A101 is predicted to be single-stranded vs. remaining base-paired with U13 in the Mn-stabilized aptamer. However, past transcript length ~220 nt, A101 is generally more reactive +Mn (SI Fig. 12D), indicating that the Watson-Crick face of A101 is transiently exposed +Mn at the end of expression platform folding, perhaps due to differences in the expression platform tertiary structure. A103 in L1 remains generally more reactive –Mn at pH 7.2 and 8.5 for the entirety of expression platform folding (SI Fig. 12D). At transcript length ~200 nt, A103 reactivity increases across all conditions, with the absolute reactivity values being higher –Mn (SI Fig. 12D). A103 is predicted to base-pair with G9 in the Mn-stabilized aptamer vs. single-stranded –Mn (Fig. 6C); thus, higher A103 reactivity -Mn is consistent with these predictions. Notably, A14, A101, and A103 are reactive towards DMS in general across all conditions tested, indicating that L1 transiently melts during expression platform folding in a pH- and Mn-dependent manner.

Lastly, we analyzed co-transcriptional Mn- and pH-dependent reactivity differences in the P1.1 switch helix, the key structural riboswitching element. C2 and A3 at the base of the P1.1 switch helix undergo a steady increase in reactivity from transcript lengths ~180-195 nt, followed by a reactivity drop ~200 nt and subsequent increase across all conditions tested (SI Fig. 12D). There is no clear Mn-dependent trend observed for C2 and A3, suggesting that a disrupted P1.1 is transiently sampled even in the presence of Mn. C104 spikes in reactivity at transcript length ~195 nt and the amplitude of this spike is higher –Mn (SI Fig. 12D), supporting that the C104-G8 base-pair in the P1.1 helix is disrupted –Mn before C104 alternatively pairs with G203 in the expression platform (Fig. 6C). A106 exhibits high reactivity at pH 7.2 in general for the entire duration of expression platform folding, despite its base-pairing U6 in P1.1 (SI Fig. 12D). Further, A106 reactivity is higher at pH 7.2 +Mn and pH 8.5 +Mn compared to their respective –Mn conditions (SI Fig. 12D). This suggests that in the presence of Mn, the Watson-crick face of A106 is more often accessible for DMS methylation. Lastly, A108 is generally more reactive at pH 7.2 during expression platform folding and demonstrates a clear pH-dependent spike in reactivity at transcript length ~190 nt, with the amplitude of the spike being higher at pH 7.2 ±Mn (SI Fig. 12D). Taken together, our observations of *mntP* L3, L1, and P1.1 reactivities during expression platform folding highlight that a global structural change occurs in the expression platform at transcript lengths ~200-205 nt, which is likely prompted by the start of strand invasion at P1.1 in the aptamer.

### Supplementary Note 7

*pH- and Mn-dependent reactivity changes in mntP equilibrium refolded intermediates*. We expanded our analysis of equilibrium refolded *mntP* intermediates to other nucleotides in the metal-sensing core, A48 and U54. At pH 8.5, A48, the A11 stacking partner, markedly drops in reactivity ±Mn at transcript length ~200 nt, with its reactivity trending higher +Mn (SI Fig. 13D). While a similar trend is observed for co-transcriptionally folded intermediates, the magnitude of the Mn-dependent reactivity difference is smaller in the equilibrium refolded context. This result indicates that the *mntP* L3 in the aptamer can still perform divalent sampling even with the fully folded expression platform. A similar trend was observed at pH 7.2, illustrating that pH affects L3 structure when the expression platform is present (SI Fig. 13D). U54 in L3 also shows a clear difference in reactivity between co-transcriptionally folded vs. equilibrium refolded intermediates. Starting at transcript length ~115 nt, U54 is more reactive +Mn at pH 8.5 and this trend is maintained for the entirety of expression platform folding (SI Fig. 13D). Although a similar trend is observed for equilibrium refolded intermediates, the magnitude of the reactivity difference is decreased, indicating that equilibrium refolding impacts L3 structure and consequently its sampling for Mn. Taken together with A48, these data support that L3 structure is altered in the equilibrium refolded context.

### Supplementary Note 8

We observed that, regardless of pH or presence of Mn, the T7-synthesized, equilibrium refolded *mntP* riboswitch assumes the same translationally inactive conformation, supported by DMS- and 1M7-constrained secondary structures (SI Fig. 14A). Nevertheless, interesting trends were extracted from the refolded full-length riboswitch chemical probing.

*pH- and Mn-dependent trends in L3 reactivity.* We observed interesting reactivity trends in the L3 loop for *mntP*. First, the absolutely conserved A50 shows less reactivity +Mn vs. –Mn when the RNA was refolded at pH 7.2, likely due to its direct coordination of Mn (SI Fig. 14B). However, there are no significant reactivity changes for A50 when RNA is refolded at pH 8.5 ±Mn, demonstrating that the flexibility of this nucleotide is limited by alkaline pH alone (SI Fig. 14B-C). In DMS-modified *mntP* RNA, A50 is less reactive +Mn at both pH (SI Fig. 14D), further supporting its direct coordination with Mn. Lastly, A48 in *mntP* L3 is less reactive towards 1M7 in general at alkaline pH but does not show a ligand-dependent reactivity difference (SI Fig. 14B-C). In contrast to 1M7 data, we observed that A48 was more reactive towards DMS +Mn at either pH (SI Fig. 14D). This observation supports that Mn binding specifically orients the Watson-Crick face of L3 such that A48 is more susceptible to DMS methylation. Together, these data support that L3 retains its ability to sample for divalents even when the RNA is refolded into an inactive conformer with a disrupted metal-sensing core.

*pH- and Mn-dependent 1M7 reactivity trends in the mntP expression platform.* In the *mntP* expression platform, several pH- and ligand-dependent reactivity differences are observed when RNA is modified with 1M7. Upon a shift to alkaline pH, A117 and A118 in P5 become less reactive at both ±Mn (SI Fig. 14C). At alkaline pH, other P5 nucleotides G140-A144 are less reactive -Mn. Interestingly, this effect becomes overall less pronounced when Mn is introduced and C139 shows a marked increase in reactivity at pH 8.5 +Mn (SI Fig. 14B-C). Finally, when Mn is present, the P5 loop from G174-G183 shows a variety of reactivity difference upon a shift to alkaline pH (Fig. 14C). Notably, G174 and C179 are both more reactive at alkaline pH compared to neutral pH when Mn is present while U176, U180, and G181 are less reactive (SI Fig. 14C). RBS nucleotides are reactive with 1M7 at all conditions, but show some interesting reactivity differences. At pH 7.2, A215 is more reactive +Mn than –Mn while G216 and G217 are less reactive +Mn (SI Fig. 14B). In contrast, A215 is also less reactive ±Mn at pH 8.5 (SI Fig 14C).

Taken together, these findings show that shifts to alkaline pH ±Mn affect the reactivity of many single stranded nucleotides in the *mntP* expression platform.

**Supplementary Note 9**

*pH- and Mn-dependent reactivity changes in alx equilibrium refolded intermediates*. Comparison between co-transcriptionally folded and equilibrium refolded *alx* intermediates revealed pH- and Mn-dependent reactivity trends for multiple nt in the *alx* expression platform that were unique to equilibrium refolded intermediates (see SI Figs. 17A-B for *alx* expression platform reactivity heatmaps, pH 7.2 and 8.5 ±Mn). A137 demonstrates a stark increase in reactivity ±Mn starting at transcript length ~150 nt, which reaches an apex at transcript length ~170 nt and then decreases until transcript length ~180 nt (SI Fig. 17C). The peak in reactivity observed for A137 ±Mn in equilibrium refolded intermediates is markedly higher than its reactivity in co-transcriptionally folded intermediates at both pH 7.2 and 8.5 (SI Fig. 17C), pointing to a distinct structural transition induced by equilibrium refolding spanning transcript lengths ~150-180 nt. Further, the amplitude of the A137 peak for equilibrium refolded intermediates is Mn-dependent with A137 being more reactive in the presence of Mn (SI Fig. 17C), indicating that Mn can influence the expression platform structure post-transcriptionally.

**Supplementary Note 10**

Like with *mntP*, the equilibrium refolded *alx* riboswitch assumes the same translationally inactive conformation regardless of pH or presence of Mn, as supported by DMS- and 1M7-constrained secondary structures (SI Fig. 20A). The following trends of interest were observed.

*pH- and Mn-dependent trends in L3 reactivity.* For *alx*, we saw that the strictly conserved A37 in L3 is always less reactive +Mn regardless of pH when modified with both 1M7 and DMS (SI Fig. 20B-D). Similar to *mntP*, this provides additional evidence that L3 can sample for divalents even when the RNA is largely refolded into the inactive conformation. When the *alx* aptamer-only was refolded, we observed that L3 becomes less flexible at pH 7.2 +Mn (SI Fig. 1D), with the exception of U38 and U39 that were shown to be flipped out of the A-minor stack in a published crystal structure(1). In contrast to the 1M7-probed aptamer at pH 7.2 +Mn, the full-length *alx* riboswitch RNA shows increased reactivity for U35 and C36 (SI Fig. 20B), likely due to a difference in L3 positioning when RNA is refolded with the expression platform.

*pH- and Mn-dependent 1M7 reactivity trends in alx expression platform.* In the *alx* expression platform, the loop nt G129-A132 in P5 are more reactive +Mn_at alkaline vs. neutral pH (SI Fig. 20C). In P6, A153 and G154 are less reactive when refolded -Mn at pH 8.5 than at pH 7.2 (SI Fig. 20C). In the presence of Mn, A153 and C155 are less reactive when refolded at alkaline pH than at neutral pH while A156 is more reactive (SI Fig. 20C). Interestingly, U169 in the P6 loop of the inactive conformer is less reactive upon a shift to pH 8.5 ±Mn while even though U170-C172 in the same loop are more reactive upon such shift, demonstrating clear pH reactivity differences between proximal nt within the same structural element (SI Fig. 20C). C171 and C172 are also more reactive when Mn is present at neutral pH (SI Fig. 20C) We observe that RBS nucleotides are overall less reactive +Mn, where A198 and A201 are less reactive at pH 7.2 and G199 and G200 are less reactive at pH 8.5 (SI Fig. 20B-C). Surprisingly, G199 and G200 are more reactive at pH 8.5 -Mn than pH 7.2 -Mn, but adding Mn abolishes this reactivity difference (SI Fig. 20C). Taken together, these RBS reactivity differences demonstrate condition-dependent fluctuations in the flexibility of these nucleotides when the RNA has been equilibrium refolded.

### Supplemental Methods

#### Proteins

Q5 High-Fidelity DNA Polymerase, Vent (exo-) DNA polymerase, Mth RNA Ligase (as part of the 5' DNA Adenylation kit), T4 RNA Ligase 2 truncated KQ, ET SSB, RNase H, RNase $I_f$, T4 DNA Ligase, SalI, HindIII, EcoRI, PstI, KpnI, BamHI, XbaI, DpnI, TIPP, NEBNext rRNA depletion kit, and HiScribe T7 High Yield RNA Kit were purchased from New England Biolabs. TURBO DNase, SuperaseIN RNase Inhibitor, and SuperScript II were purchased from Thermo Fisher Scientific. SAvPhire Monomeric Streptavidin was purchased from Sigma-Aldrich. Catalog numbers for all commercial proteins are tabulated in Supplementary Table 1. Preparation of *E. coli* RNAP, *E. coli* $\sigma^{70}$, and *E. coli* NusA were performed as described previously(7).

#### Bacterial strains

The strains used in this study are derivatives of *E. coli* K-12 and are tabulated in Supplementary Table 4.

The Δ*alx*::Kan mutations were sourced from strains in the Keio collection(8) and introduced into MC4100 *E. coli* strain via P1 transduction. Chromosomally encoded T7 RNAP was introduced to this MC4100Δ*alx* strain using P1 transduction for the *alx* and *mntP* reporter fusions and was expressed from a *lac* promoter.

#### Plasmids and cloning procedures

The plasmids used in this study are tabulated in Supplementary Table 5.

The plasmid pMIS309 carrying the template for promoter-initiated transcription on the wild-type *mntP* riboswitch sequence (Supplementary table 5) was constructed by PCR-amplifying from a plasmid template, pRA57, (Supplementary table 5) from 486 nt upstream and 42 nt downstream of the *mntP* translation start site using oligos 0257 and 0258. The PCR product was then cloned into the BamHI and EcoRI sites of pUC19. The plasmid was assembled *via* restriction-ligation using T4 DNA Ligase (New England Biolabs).

The plasmid pMIS313 carrying the template for TECprobe-VL experiments on the *alx* riboswitch (Supplementary table 5) was constructed using a two-step PCR strategy. The PCRs employed oligos with 5' overhangs that introduced the PRA1 promoter and SC1 hairpin upstream of the *alx* riboswitch and additional downstream *alx* sequence. The first PCR used pMIS301 (Supplementary table 5) as a template and oligos 0155 and 0187. The second PCR used the purified first step PCR product as a template and oligos 0156 and 0188. The second-step PCR product was cloned into the HindIII and EcoRI sites of pUC19 *via* Gibson assembly using NEBuilder HiFi DNA Assembly Master Mix (New England Biolabs).

The plasmid pMIS315 carrying the template for TECprobe-VL experiments on the *mntP* riboswitch (Supplementary table 5) was constructed using a two-step PCR strategy. The PCRs employed oligos with overhangs that introduced the PRA1 promoter and SC1 hairpin upstream of the *mntP* riboswitch. The first PCR used pRA57 (Supplementary table 5) as a template and oligos 0258 and 0259. The second PCR used the purified first step PCR product as a template and oligos 0264 and 0265. The second step PCR product was cloned into the BamHI and EcoRI sites of pUC19 *via* restriction-ligation using T4 DNA Ligase (New England Biolabs).

The plasmid pMIS349 carrying the *alx* translational reporter with the A37T mutation in the sequence encoding the *alx* riboswitch was constructed by overlap extension PCR using published methods(9) with oligos 0229,0232,0458, and 0459.

The plasmid pMIS508 carrying the template for the *alx* aptamer under the control of a T7 promoter was constructed by cloning the template sequence (Supplementary Table 5) as a gene block from Integrated DNA Technologies into the BamHI site of pUC57. The plasmid was assembled via restriction-ligation using T4 DNA Ligase (New England Biolabs).

The plasmid pMIS509 carrying the template for the *mntP* aptamer under the control of a T7 promoter was constructed by cloning the template sequence (Supplementary Table 5) as a gene block from Integrated DNA Technologies into the BamHI site of pUC57. The plasmid was assembled via restriction-ligation using T4 DNA Ligase (New England Biolabs).

The plasmid pMIS511 carrying the template for *in vitro* SHAPE/DMS-MaP of the *alx* riboswitch was constructed by cloning the template sequence (Supplementary Table 5) as a gene block from Integrated DNA Technologies into the PstI and XbaI sites of pUC57. The plasmid was assembled via restriction-ligation using T4 DNA Ligase (New England Biolabs).

The plasmid pMIS513 carrying the template for *in vitro* SHAPE/DMS-MaP of the *alx* aptamer was constructed by cloning the template sequence (Supplementary Table 5) as a gene block from Integrated DNA Technologies into the PstI and XbaI sites of pUC57. The plasmid was assembled via restriction-ligation using T4 DNA Ligase (New England Biolabs).

The plasmid pMIS518 carrying the template for *in vivo* DMS-MaP of the *alx* riboswitch was constructed by PCR amplifying the template sequence (Supplementary Table 5) from pMIS447 (BioBasic) using oligos 0005 and 0026 and cloning it into the XbaI and PstI sites of pUC19. The plasmid was assembled via restriction-ligation using T4 DNA Ligase (New England Biolabs).

The plasmids with *alx* and *mntP* reporter gene fusions under the control of a native *E. coli* promoter are pRA54 (*alx* translational reporter) and pRA57 (*mntP* translational reporter) and were constructed in prior work(10) and used as controls in this work (Supplementary Table 5). Reporter gene fusions under the control of a T7 promoter (*alx*: pMIS501 and *mntP*: pMIS517, Supplementary Table 5) were constructed via sequential PCR amplification. For *alx*, the first amplification generated 208 nucleotides of the *alx* 5'-UTR under control of a T7 promoter. This was amplified from pMIS303 using oligos 0202 and 0203, adding a PstI restriction site. For the second amplification, *alx-lacZ* was amplified from pRA54 for the translational fusion using oligos 0204 and RAV23, giving a 97 nucleotide overlap with the first PCR amplification. A third PCR step using 0202 and RAV23 combined the first two PCRs into the full length *alx* translational fusion under the control of a T7 promoter. The translational fusion (pMIS501) was cloned into the PstI and BamHI sties of a single copy plasmid pMU2386. The plasmids were assembled via restriction-ligation using T4 DNA Ligase (New England Biolabs). For *mntP*, the first amplification generated 228 nucleotides of the *mntP* 5'-UTR under the control of a T7 promoter. This was amplified from pMIS310 using oligos 0202 and 0454, adding a PstI restriction site. For the second amplification, *mntP-lacZ* was amplified from pRA57 for the translational fusion using oligos 0455 and RAV126, giving a 95 nucleotide overlap with the first PCR amplification. A third PCR step using 0202 and RAV126 combined the first two PCRs into the full length *mntP* translational fusion under the control of a T7 promoter. The translational fusion (pMIS517) was cloned into the PstI and SalI sites of a single copy plasmid pMU2386. The plasmids were assembled via restriction-ligation using T4 DNA Ligase (New England Biolabs).

#### ***In vitro*** **transcription template preparation**

Supplementary Table 3 tabulates details for the linear double-stranded DNA templates prepared for this study, including oligonucleotides used for PCR amplification and cleanup method used. Supplementary Table 6 provides all DNA template sequences used in this work.

Unmodified or terminally biotinylated DNA templates for promoter-initiated *in vitro* transcription were prepared by PCR with high-fidelity Q5 DNA polymerase (New England Biolabs), treated with DpnI (New England Biolabs) if a plasmid was the PCR template, and purified by either phenol extraction followed by ethanol precipitation or Monarch PCR & DNA cleanup kit (New England Biolabs).

DNA templates for TECprobe-VL were PCR amplified from a linear DNA template containing a terminal biotin modification (*alx*: Template 2, *mntP*: Template 5) using Vent (exo-) DNA polymerase (New England Biolabs). The PCR contained 1X ThermoPol Buffer (New England Biolabs), 200 μM dNTP Solution Mix, 250 nM forward primer, 250 nM reverse primer, 0.02 nM template DNA, 0.02 U/μl Vent (exo-) DNA polymerase, and a concentration of biotin-11-dNTPs (AAT BioQuest and Biotium) that promoted incorporation of ~3 biotin modifications in the transcribed region of the DNA template(11). The PCR was performed using the following thermal cycler protocol: 95 °C for 3 min [95 °C for 20 s, 58 °C for 30 s, 72 °C for 25 s] x 30 cycles, 72 °C for 5 min, hold at 12 °C. The PCR products were purified by phenol extraction followed by ethanol precipitation and quantified using the Quant-iT dsDNA Assay Kit (HS) (Invitrogen) with a BioTek Synergy H1 microplate reader (Agilent).

Linear DNA templates were prepared for transcription by T7 RNAP via PCR amplification of a plasmid using Q5 DNA Polymerase. For the *alx* riboswitch SHAPE/DMS-MaP template DNA, oligos 0026 and 0277 were used to amplify the template DNA from plasmid pMIS511 and DpnI was used to digest original plasmid DNA (Template 8). For the *alx* aptamer SHAPE/DMS-MaP template DNA, oligos 0026 and 0277 were used to amplify the template DNA from plasmid pMIS513 and DpnI was used to digest original plasmid DNA (Template 9). For the *mntP* riboswitch SHAPE/DMS-MaP template DNA, oligos 0297 and 0298 were used to amplify the template DNA from plasmid pMIS314 and DpnI was used to digest original plasmid DNA (Template 10).

**Synchronized single-round *in vitro* transcription from a promoter**

To monitor transcription on the *mntP* riboswitch sequence when initiated from its native promoter, we performed synchronized, single-round *in vitro* transcription assays. The template contained the *mntP* promoter (261 nt upstream of the transcription start site), the wild-type *mntP* riboswitch (nt 1-225), and nt 1-45 of *mntP* coding sequence (Template 4, Supplementary Table 3). To prepare the RNAP holoenzyme (holoRNAP), *E. coli* RNAP (2 $\mu$M) and $\sigma^{70}$ (4 $\mu$M) were combined in 1x transcription buffer (20 mM Tris-Cl, pH 8.5, 150 mM KCl, 10 mM $MgCl_2$, and 5 mM DTT) and incubated for 30 min at 37 °C. Following holoRNAP assembly, *in vitro* transcription reactions were initiated with an ApU dinucleotide and synchronized, taking advantage of the natural lack of cytosines between positions 3 and 15 of the *mntP* riboswitch. The initiation mix for *in vitro* transcription thus lacked CTP and contained 0.1 $\mu$M of the holoRNAP, 200 nM of the DNA template, 1 $\mu$M ATP, 2.5 $\mu$M CTP, 2.5 $\mu$M GTP, 150 $\mu$M ApU (Trilink), [$\alpha$-$^{32}$P]ATP (Revvity), and transcription buffer. After allowing initiation to proceed for 15 min at 37 °C, rifampicin (Fisher) was added to a final concentration of 10 $\mu$g/ml, to inhibit reinitiation. Completion of this step yielded ECs stalled at the G15 position on the *mntP* riboswitch.

Next, transcription elongation reactions were performed by combining 9 $\mu$l of ECs with 1 $\mu$l of an NTP mix containing 1 mM each ATP, UTP, CTP, and GTP (100 $\mu$M final, each NTP). All elongation reactions were performed at 37 °C. $MnCl_2$ was introduced concurrently with NTPs to a final concentration of 100 $\mu$M to test the effect of Mn on transcription elongation and pausing. The transcription factor NusA was introduced concurrently with NTPs to a final concentration of 250 nM to test its effect on transcription elongation and pausing. After beginning elongation, the reaction was quenched at desired time points with an equal volume of 2x stop buffer (90 mM Tris-borate buffer, 8 M urea, 50 mM EDTA, and 0.02% of both xylene cyanol and bromophenol blue). The time points collected were 10 s, 30 s, 60 s, 90 s, 2 min, 3 min, and 4 min. Additionally, a “chase” reaction was performed with each NTP present at a final concentration of 500 $\mu$M. Radiolabeled RNAs in the quenched samples were denatured for 2 min at 95 °C and separated using denaturing PAGE (6% 19:1 acrylamide:bisacrylamide, 8 M urea). Samples were run alongside a 5'-$^{32}$P radiolabeled ladder (MspI-digested pBR322 plasmid). The gel was exposed to a PhosphorImager screen, and the screen was scanned using a Typhoon PhosphorImager.

#### ***mntP*** **riboswitch pause mapping using scaffold-based transcription**

To precisely map pauses in the *mntP* riboswitch, transcription elongation assays using nucleic acid scaffolds were performed. Each scaffold initiated transcription upstream of the sequence window where pauses were expected to occur.

Multiple nucleic acid scaffolds were designed to map pauses on the *mntP* riboswitch sequence towards the end of aptamer domain folding and beginning of expression platform folding. The sequence window for scaffold-based pause mapping experiments was narrowed based on results from promoter-based assays. These experiments are specifically designed to map elemental pauses, meaning that the influence of local RNA structure on pause duration is omitted as a variable. Prior work has established that the elemental pause is an obligate precursor to longer-lived pauses stabilized by nascent RNA structure(12). Therefore, these minimalistic experiments are relevant for mapping pauses encountered by ECs when the full-length riboswitch RNA is present. A brief description of the scaffolds is presented here and are described pictorially in Supplementary Figure 5. The nucleotide numbering corresponds to the numbering in the full-length *mntP* riboswitch sequence. Scaffold 1 employed an 8-mer starting RNA to position RNAP upstream of nt 79-100. Scaffold 2 used a 10-mer RNA to position RNAP upstream of nt 101-113. Two separate experiments with different starting RNAs were performed on scaffold 3. The first 10-mer RNA positioned RNAP upstream of nt 114-131, and the second 10-mer RNA positioned RNAP upstream of nt 132-139. Past these defined sequence windows for pause mapping on scaffolds 1-3, the RNA emerging from RNAP could form non-native secondary structures that may have influenced pause behavior. Therefore, RNA species outside of the defined sequence windows were omitted for pause analysis.

All scaffold-based transcription elongation assays used the following protocol. First, the nucleic acid scaffold was annealed in a buffer (20 mM Tris, pH 7.2, 150 mM KCl, and 5 mM DTT) with 5 $\mu$M RNA and 10 $\mu$M template DNA (T-DNA). Then, ECs were reconstituted by combining RNAP with the nucleic acid scaffold in transcription buffer (20 mM Tris, pH 7.2, 150 mM KCl, and 5 mM DTT) and incubated for 15 min at 37 °C. Next, the non-template DNA (NT-DNA) was added and incubation continued for 10 min at 37 °C to complete EC assembly. The ratio of RNA:T-DNA:NT-DNA:RNAP was 1:2:5:3 corresponding to concentrations of 0.5, 1, 2.5, 1.5 $\mu$M, respectively. Next, the ECs were diluted fivefold, and heparin was added to the ECs at a final concentration of 0.1 mg/ml to sequester unbound RNAP. ECs were incubated with heparin for 3 min at 37 °C. To radiolabel the ECs, [$\alpha$-$^{32}$P]ATP (Revvity) was added to the ECs for 1 min at 37 °C, which extended a fraction of RNAs by 1 nt for visualization. To extend all ECs by 1 nt, ECs were incubated with unlabeled ATP at a final concentration of 2 $\mu$M for 3 min at 37 °C.

Transcription elongation reactions were performed by combining the ECs with an NTP mix containing 1 mM each ATP, UTP, CTP, and GTP (100 $\mu$M final, each NTP). $MgCl_2$ was added to a final concentration of 5 mM alongside NTP introduction. Additionally, to extend pause duration and aid with pause mapping, transcription elongation reactions were performed with three of the NTPs at a final concentration of 100 $\mu$M and the fourth NTP limiting at a final concentration of 10 $\mu$M. Samples were removed at various time points and quenched with an equal volume of 2x stop buffer. Chase reactions were performed with each NTP at a final concentration of 500 $\mu$M. In parallel, RNA sequencing ladders were generated to map pause locations. To make the ladder, the ECs were prepared as described above. The ECs were combined with ATP, UTP, CTP, and GTP to 100 $\mu$M and one 3'-dNTP (Millipore Sigma) to 1 mM. The sequencing reactions proceeded for 20 min at 37 °C and were quenched with an equal volume of 2x stop buffer. Radiolabeled RNAs from the quenched time points were heat-denatured for 2 min at 95

°C and separated using denaturing PAGE (15% 19:1 acrylamide:bisacrylamide, 8 M urea). The gel was exposed to a PhosphorImager screen, and the screen was scanned using a Typhoon PhosphorImager.
